# Computational Design of Metallohydrolases

**DOI:** 10.1101/2024.11.13.623507

**Authors:** Donghyo Kim, Seth M. Woodbury, Woody Ahern, Doug Tischer, Nikita Hanikel, Saman Salike, Jason Yim, Samuel J. Pellock, Anna Lauko, Indrek Kalvet, Donald Hilvert, David Baker

## Abstract

De novo enzyme design starts from a description of an ideal active site composed of catalytic residues surrounding the reaction transition state(s), and builds a protein structure that contains this site^1–7^. Generative AI methods such as RFdiffusion^11,12^ now enable the direct generation of proteins around active sites, but to date, such scaffolding has required specification of both the position in the sequence and the backbone coordinates of each catalytic residue, which complicates sampling. Here we introduce a generative AI method called RFdiffusion2 that overcomes these limitations and use it to design zinc metallohydrolases starting from a density functional theory description of the active site geometry. Of an initial set of 96 designs tested experimentally, the most active has a *k*_cat_/*K*_M_ of 16,000 M^−1^ s^−1^, orders of magnitude higher than previously designed metallohydrolases.^6,7,13,14^ A second round of 96 designs yielded 3 additional highly active enzymes, with *k*_cat_/*K*_M_ up to 53,000 M^−1^ s^−1^ and *k*_cat_ up to 1.5 s^−1^. The structures of the four enzymes are very different from each other and from the structures in the PDB. Each enzyme positions the reaction substrate almost perfectly for nucleophilic attack by a water molecule activated by the bound metal, and are predicted by PLACER^15^ and Chai-1^44^ to have highly preorganized active sites. The ability to generate highly active catalysts straight out of the computer, without experimental optimization, using quantum chemistry calculated active site geometries should open the door to a new generation of potent designer enzymes.^16,17^

## Main

Metallohydrolases catalyze some of the most difficult hydrolysis reactions in biology by using their bound metal ion(s) to activate a water molecule positioned adjacent to the substrate bond to be cleaved.^18–20^ Engineering new metallohydrolases is currently of considerable interest for degrading human-generated environmental pollutants for which there has not been sufficient time for efficient hydrolytic enzymes to evolve.^21–23^ Protein engineering has expanded the scope of substrates that metallohydrolases can hydrolyze, but this often requires initial promiscuous activity.^24,25^ De novo enzyme design has been used to generate new metallohydrolases^6,8,13^, but these have had quite low activity and efficiency requiring extensive directed evolution to match those of native enzymes.^8^ Given a description of an ideal metallohydrolase active site, de novo metalloenzyme design seeks to find or generate a protein scaffold that positions the catalytic residues, metal(s), and substrate/transition state with high accuracy.^26,27^ RFdiffusion has been used successfully to scaffold sites, but the search has been complicated by the requirement for the specification of the sequence positions and conformations of the catalytic residues.^11,12,28^

We reasoned that a generative AI design method which only required the specification of sidechain functional group positions around a reaction transition state, capable of sampling over all possible sequence positions and conformations of these residues, could more readily satisfy complex catalytic constraints.^16,17,29,30^ We set out to develop such an approach, and use it to design new metallohydrolases starting from a quantum chemistry generated active site description with a bound metal cofactor.

To enable sequence position and sidechain rotamer-agnostic enzyme design, we developed a generative AI flow-matching model called RFdiffusion2.^33^ RFdiffusion2 extends the RFdiffusion capabilities of generating scaffolds that position a set of functional residues (a “motif”) in two key ways. First, it enables atomic substructure scaffolding: RFdiffusion can only scaffold backbone-level motifs (with the sidechain and backbone atom N-C_α_-C=O positions specified), while RFdiffusion2 can scaffold arbitrary atom-level motifs (any subset of amino acid heavy atoms). This is important for enzyme design because it allows users to specify only the key functional group positions interacting with the reaction transition state, rather than the full sidechain and backbone conformation. Second, RFdiffusion2 enables sequence-position-agnostic scaffolding: RFdiffusion requires specification of the primary sequence positions of the motif residues, but RFdiffusion2 can scaffold motifs whose primary sequence positions are unknown. RFdiffusion2 replaces diffusion with flow matching^31,32^ and achieves sequence position-agnostic atomic substructure scaffolding by providing randomly selected native atomic coordinates (but not their sequence positions) during training in addition to the partially noised, sequence-labeled atomic coordinates. With these improvements, RFdiffusion2 generates diverse proteins starting directly from catalytic configurations consisting of input functional group positions and substrate coordinates. Allowing the model to resolve the *a priori* unknown degrees of freedom (i.e., the primary sequence positions and sidechain rotamer conformations of the catalytic sidechains) is considerably more effective at generating self-consistent design solutions than randomly sampling (the space is far too large to exhaustively sample) those degrees of freedom before inference, as was necessitated with RFdiffusion. A detailed description of RFdiffusion2 training and benchmarking results for a wide range of active site scaffolding problems is described in a separate reference.^33^

As an initial test of RFdiffusion2, we chose to design a zinc metallohydrolase for a fluorogenic ester, 4-methylumbelliferyl phenylacetate (4MU-PA), as a model reaction (Fig. 1A). We began by using density functional theory (DFT) to identify the geometry of the rate-determining Zn(II)-OH nucleophilic attack on the substrate ester. Four distinct catalytic arrangements based on the stereochemistry of the tetrahedral intermediate and the nature of the oxyanion hole were considered (Fig. 1B and Fig. S3; for further details see SI, section 3.1). These calculations provide the coordinates of the three Zn(II)-binding imidazole rings, the metal, and the transition state. Our previous RFdiffusion approach required the backbone coordinates and residue positions as inputs; this required upfront sampling of rotameric states and sequence position for each histidine. This cannot be done exhaustively: even with relatively coarse sampling around the sidechain chi angles χ_1_, χ_2_, and the backbone torsion angle ψ there are on the order of 10^18^ possible choices for the sidechain conformations and sequence placements of our full catalytic site (Fig. 1C and Fig. S4). Whereas each RFdiffusion run has to be initialized with a specific (and generally randomly selected) choice from this enormous set of combinations, RFdiffusion2 as described above searches the entire space in each trajectory.

**Fig. 1.**
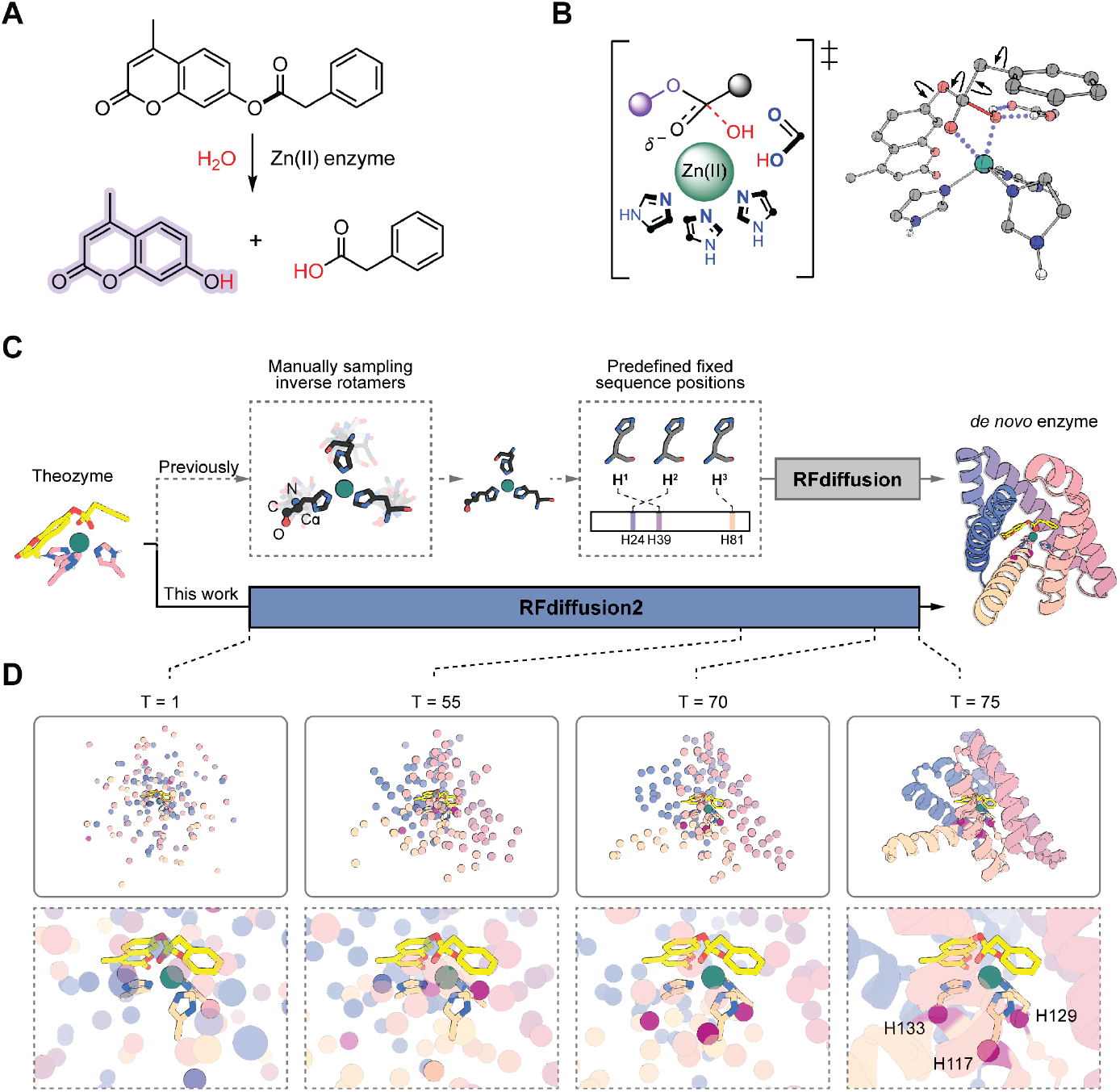
RFdiffusion2 design method. (**A**) Hydrolysis of 4MU-PA yields phenylacetic acid and a fluorescent coumarin product. (**B**) Example theozyme for Zn(II)-OH nucleophilic attack of the 4MU-PA ester. 2D representation (left) and 3D DFT model (right). Arrows on the 3D model represent sampled conformational flexibility. (**C**) Comparison of scaffold generation around an input theozyme using previous backbone centric RFdiffusion (top row) versus interaction functional group centric RFdiffusion2. RFdiffusion requires explicit upfront sampling of side chain conformations and residue sequence positions, whereas RFdiffusion2 only requires the transition state complex and the catalytic side chain functional groups, implicitly sampling sequence space and rotameric space during inference. (**D**) Snapshots of the global structure and active site from model X_T_ during an RFdiffusion2 inference trajectory. The coordinates of the input transition state complex and catalytic functional groups stay fixed during inference while the overfall backbone structure and sequence positions and unspecified atoms of the catalytic side chains are sampled by RFdiffusion2. The C_α_ atoms that host the catalytic histidines at the end of the trajectory are retrospectively highlighted as red spheres; these C_α_ atoms are not predetermined but rather move into the frame to host the fixed sidechains as the global structure forms around the motifs.

RFdiffusion2 inference trajectories were used to build protein scaffolds housing the DFT-generated minimal active site configurations, referred to as theozymes.^2,34^ Several snapshots from a representative trajectory are shown in Fig. 1D, transforming random noise on the left into the final backbone on the right (Movie S1). The C_α_ atoms of each residue (shown in colored spheres representing final sequence position) are initially sampled from a Gaussian distribution, while the target functional atom positions (shown in sticks) stay fixed. As the trajectory proceeds from left to right, the global structure takes shape around the motifs, with the fixed histidine sidechains eventually connecting to C_α_ atoms of the protein backbone at sequence positions of the network’s choosing. 5120 RFdiffusion2 inference trajectories were carried out starting from different random seeds and for each of the resulting protein scaffolds, sequences were generated using ProteinMPNN.^35^ The catalytic geometry and interactions with the transition state of those designs for which the AlphaFold2^36^ (AF2) predicted structure was close to the design model were further optimized using iterative LigandMPNN^37^ and constrained Rosetta repacking and minimization^38^ (Fig. S1; see SI, section 3.1). Designs containing a proposed general base positioned to activate the water molecule (i.e., Glu/Asp/His within H-bonding distance of the Zn(II)-bound water) and sidechain hydrogen bonds stabilizing the transition state oxyanion (if applicable), and that AF2 predicted to adopt the target structure, were characterized with PLACER^15^ to assess active site preorganization (see below). 96 designs were selected for experimental characterization based on predicted active site geometry and preorganization (Fig. S5; see SI, section 3.1).

Linear DNA fragments encoding the 96 designs were cloned into a plasmid encoding a C-terminal Strep-tag and used to transform *E. coli*, and the resulting proteins were purified using Streptactin affinity chromatography. 86/96 designs were expressed and soluble as judged by SDS-PAGE analysis of the eluants (Fig. S6). Purified designs were supplemented with zinc sulfate, and hydrolysis of 4MU-PA was monitored by fluorescence. Five designs (A1, A8, B9, C4, and F7) had activity well above background (Fig. 2B and Fig. S7). Sequence-verified single clones for each of these design hits were expressed and purified by affinity chromatography followed by size-exclusion chromatography to obtain pure, monomeric protein fractions (Fig. S8 and Fig. S9). Michaelis-Menten kinetic characterization of the purified variants revealed a *k*_cat_/*K*_M_ of 16000 ± 2000 M^−1^ s^−1^ for the most active A1 design, and *k*_cat_/*K*_M_ values in the range of 35-140 M^−1^ s^−1^ for the other four designs (Fig. 2, C,D; Fig. S10; Table S1). For comparison, the *k*_cat_/*K*_M_ of previously designed metallohydrolases ranged from 3 to 60 M^−1^ s^−1^ (Table S3).^8^ A1 is also quite a robust enzyme, retaining activity for at least 1000 turnovers (Fig. 3E). Additionally, A1 differs considerably from previously described structures: the most similar structures identified through TM alignment with the PDB and AFDB have template modeling (TM) scores^39^ of 0.41 and 0.49, respectively, and do not have analogous arrangements of catalytic residues (Fig. S16). We refer to A1 as zinc metalloesterase 1 (ZETA_1) throughout the remainder of the text.

**Fig. 2.**
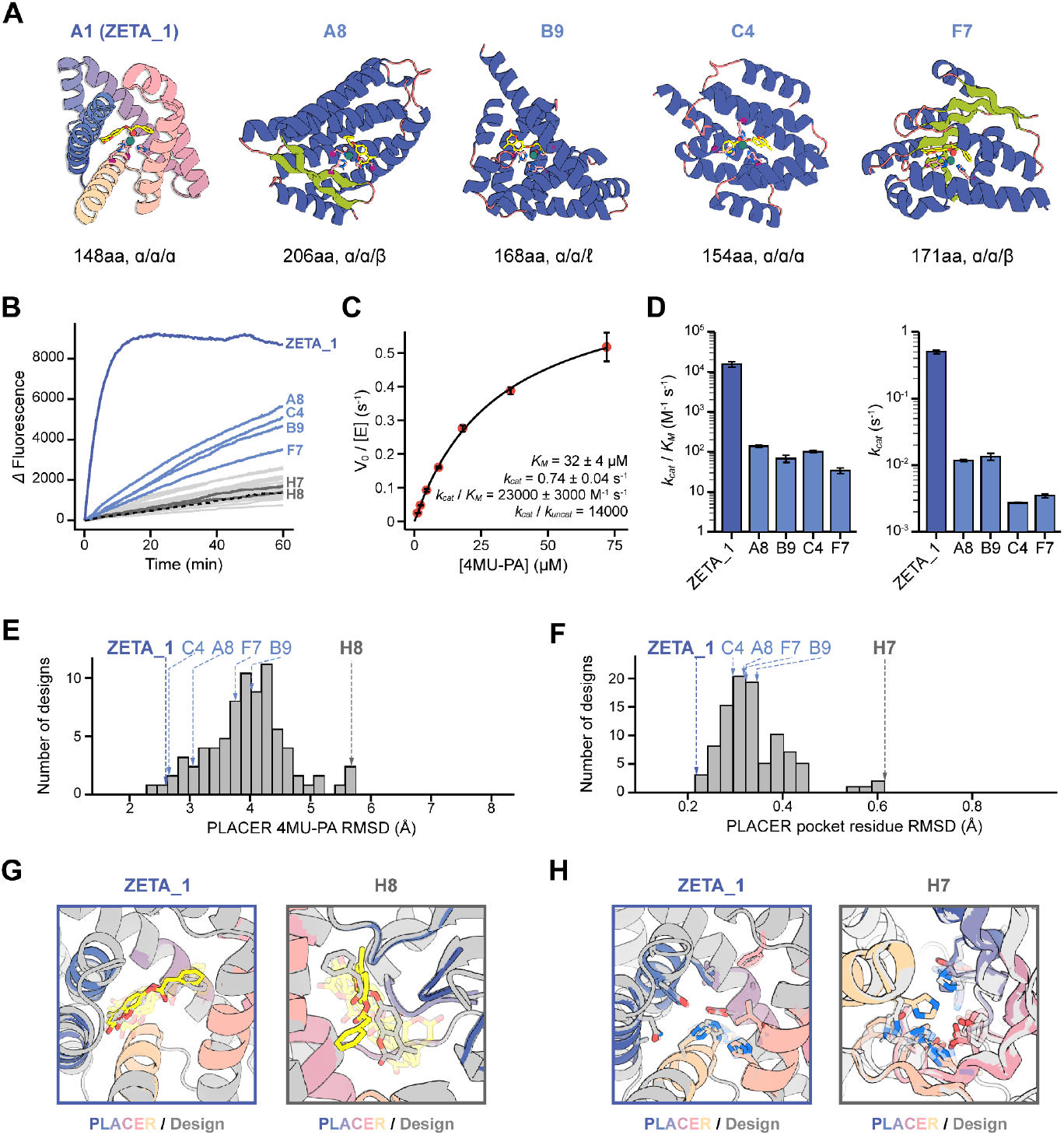
Activity characterization and PLACER preorganization assessment. (**A**) Design models of most active designs. Sequence length and the secondary structure of each catalytic histidine are indicated below. (**B**) Reaction progress curves. Dotted black line is the buffer background. (**C**) Michaelis-Menten characterization of A1, hereafter referred to as ZETA_1. The y-axis is the initial rate divided by the total enzyme concentration. (**D**) Michaelis-Menten parameters of most active designs. (**E**,**F**) Distribution of PLACER active site pre-organization ensemble metrics for the ordered designs. Average design-prediction substrate RMSD (E), and the average catalytic and binding residue design-prediction RMSD (F) across all predicted ensembles generated by PLACER for each design. (**G**) For ZETA_1, the substrate position in PLACER ensembles is close to the design model, while in the inactive design H7 the substrate position fluctuates widely. (**H**) For ZETA_1, the sidechains surrounding the active site are largely fixed in positions close to the original design model, while in the inactive H8 design, the sidechain positions vary considerably. Note that only the first 5 randomly generated, unranked ensemble predictions are shown for PLACER in (G,H).

**Fig. 3.**
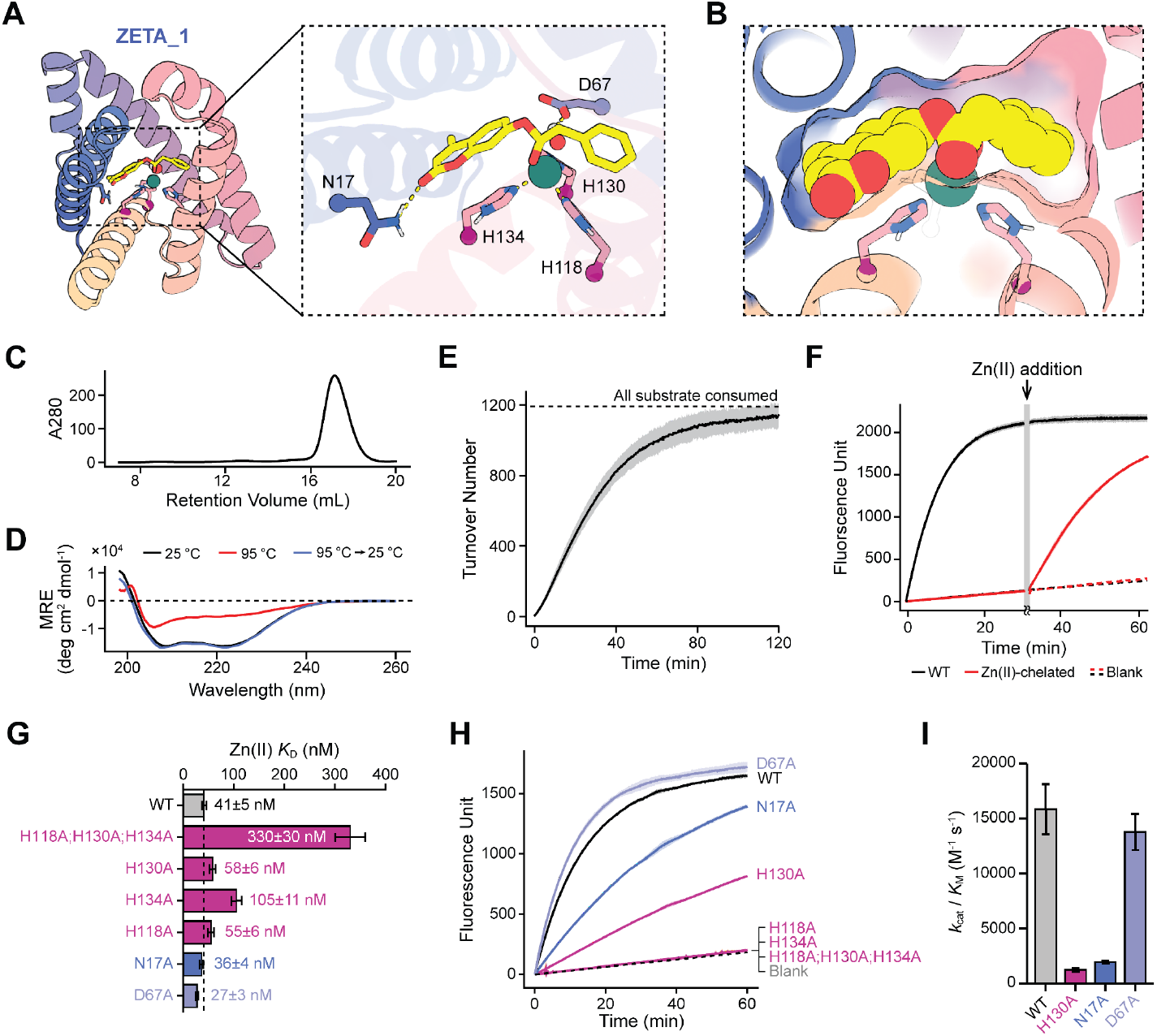
Characterization of ZETA_1 activity. (**A**,**B**) ZETA_1 design model with magnification of the active site showing (**A**) the hypothesized catalytic residues and (**B**) a surface view of the pocket revealing high shape-complementarity to the substrate. (**C**) Size-exclusion chromatogram of ZETA_1 showing a single peak corresponding to monomeric protein. (**D**) Circular dichroism spectra for ZETA_1 at 25 °C (black), 95 °C (red), and a cycle from 25 °C to 95 °C to 25 °C (blue). The spectra suggest that ZETA_1 has an alpha-helical secondary structure and that it can refold after heating and partial unfolding. (**E**) [Product]:[enzyme] turnover curve measuring the lower bound of the total turnover of ZETA_1 demonstrating full hydrolysis of 4MU-PA and >1000 turnovers within 2 hours. Note that the background reaction has been subtracted from the spectra so that every turnover can be attributed to the enzyme (Fig. S11; more details in SI, section 3.3). (**F**) Reaction progress curves for the Zn(II) *holo*- and zinc-chelated *apo*-ZETA_1 proteins showing zinc-dependent activity. Adding excess Zn(II) to the *apo*-ZETA_1 sample after 30 minutes reestablishes the activity demonstrating that zinc is essential for the catalytic mechanism of ZETA_1. (**G**) Zinc affinity of the wild-type ZETA_1 and its knockout mutants, measured as the dissociation constant (*K*_D_) where a lower value indicates tighter binding. (**H**) Fluorescence progress curves comparing the activity of the wild-type ZETA_1 to its different knockout mutants. (**I**) Michaelis-Menten *k*_cat_/*K*_M_ for the active ZETA_1 mutants compared to the wild-type.

Design ZETA_1 not only has remarkably high activity but was also the top-ranked design in our *in silico* ranking, hence its original A1 label. The structure in the absence of substrate was predicted to be very close to the design model by AlphaFold2 (Fig. S12 and Fig. S15A), and the designed active site of ZETA_1 was predicted to be highly preorganized by PLACER, with the catalytic sidechains fixed in place and the substrate held closely in its designed position, adjacent to the proposed Zn(II) position. PLACER is a deep neural network that, given a protein backbone containing a substrate, fully randomizes the positions of the substrate and all side chains within a 600-atom sphere, and then generates new coordinates for these groups^15^; repeated PLACER trajectories generate an ensemble of possible sidechain conformations and small molecule docks. Design ZETA_1 stood out from the other designs in both the extent of catalytic site pre-organization (the catalytic sidechains were largely fixed in space in catalytically competent conformations) and the positioning of the substrate/transition state in the active site (in the ZETA_1 ensemble the substrate remained largely fixed in space in the active site, whereas in the inactive designs H7 and H8 it fluctuated considerably) (Fig. 2, E-H; Movies S2-S5). Seven designs based on the same ZETA_1 backbone family were initially filtered out during the design selection phase as they had suboptimal PLACER metrics; we retrospectively expressed and purified these designs and found that they had very low or no activity, further highlighting the utility of PLACER ensemble calculations for identifying active designs (Fig. S14). These findings suggest that combining global structure prediction with detailed PLACER modeling of the active site provides a powerful approach to assessing the catalytic machinery and substrate binding geometry for design selection (Fig. S13).

The ZETA_1 active site consists of a primarily hydrophobic pocket with three histidines binding Zn(II) with their N_ε_ atoms, an aspartate as a potential general base, and an asparagine that forms a hydrogen bond to the coumarin ring (Fig. 3, A,B). As in the original theozyme model used to generate ZETA_1, the Zn(II) ion also acts as an oxyanion hole, stabilizing the forming negative charge at the transition state; there are no nearby sidechain H-bond donors (Fig. S15). Zinc is absolutely critical for ZETA_1 activity: extraction of bound Zn(II) by dialysis in the presence of the chelator 1,10-phenanthroline completely eliminated activity, and activity was subsequently restored by addition of zinc to the solution (Fig. 3D). Zinc titration experiments indicated a binding affinity (*K*_D_) for Zn(II) of 41±5 nM which is comparable to previous designed zinc enzymes^27,40^, but higher than native zinc hydrolases which typically have *K*_D_ <10 nM.^20,41–43^

We carried out mutagenesis experiments on the proposed catalytic residues to evaluate their contributions to zinc binding and catalysis (Fig. 3, G-I). In the design model, N17 positions the substrate by hydrogen bonding with the ester carbonyl of the coumarin moiety and could stabilize the developing negative charge on the leaving group; the N17A mutation led to a 8.1-fold decrease in *k*_cat_/*K*_M_ (Fig. S18). Mutation of all three metal coordinating histidine residues to alanine simultaneously (H118A;H130A;H134A), as well as two of the three single histidine-to-alanine substitutions (H118A and H134A), completely inactivated the enzyme, as expected. Mutating the third Zn(II)-coordinating residue to alanine (H130A) only resulted in a decrease of 13-fold in *k*_cat_/*K*_M_, while mutation of the proposed general base D67 to alanine had little effect on *k*_cat_/*K*_M_ and increased zinc binding affinity. These results suggest that H134/H118/H130 and H134/H118/D67 may be competing Zn(II)-binding sites due to the close proximity of the coordinating side chains of H130 and D67, corroborated by Chai-1^44^ predictions of the protein-Zn(II)-substrate complex (Fig. S15 B,C); the D67A mutation may confine the zinc to the originally designed coordination sphere with the three histidines, which is more catalytically competent. In the H130A mutant, D67 likely coordinates Zn(II) and maintains binding, albeit in a less optimal binding geometry, lowering the zinc affinity and enzyme activity.

Guided by these observations, we started from new DFT theozymes containing the catalytic base, and generated protein structures scaffolding these theozymes using a newer version of RFdiffusion2 trained from random weight initialization on a 3-fold larger data set, whereas previous versions were fine-tuned from structure prediction weights (Fig. 4A; see SI, section 3.2). Designs for which Chai-1 predictions of the protein-Zn(II)-substrate phosphoanalog complex closely matched the design models with high confidence were found by PLACER to have highly pre-organized active sites (Fig. S20). 96 such designs spanning 37 RFdiffusion2 generated backbones were selected for experimental characterization (Fig. S21). 85/96 designs were expressed and soluble (Fig. S22), and 11 spanning 3 different folds had substantial zinc-dependent 4MU-PA hydrolysis activity (Fig. 4, B,C). Michaelis-Menten analysis revealed that 5 designs had a *k*_cat_/*K*_M_ >10^4^ M^−1^ s^−1^ and 6 designs had a *k*_cat_/*K*_M_ >10^3^ M^−1^ s^−1^ (Fig. 4D and Fig. S23; Table S1). The most active designs for each backbone had a *k*_cat_/*K*_M_ = 53,000 ± 5000 M^−1^ s^−1^ (ZETA_2), *k*_cat_/*K*_M_ = 19,000 ± 2000 M^−1^ s^−1^ (ZETA_3), and *k*_cat_/*K*_M_ = 1,100 ± 200 M^−1^ s^−1^ (ZETA_4) (Fig. 4, F-H). ZETA_2 has a *k*_cat_ = 1.5 ± 0.1 s^−1^, a 3-fold increase over the *k*_cat_ of ZETA_1. RFdiffusion2 enables specification of the position of the substrate relative to the designed protein center of mass; for ZETA_2 and ZETA_3, the protein was centered near the PA and 4MU, respectively, resulting in opposite substrate binding modes in the design models (i.e., the 4MU is exposed in ZETA_2 and the PA is exposed in ZETA_3) (Fig. S24).

**Fig. 4.**
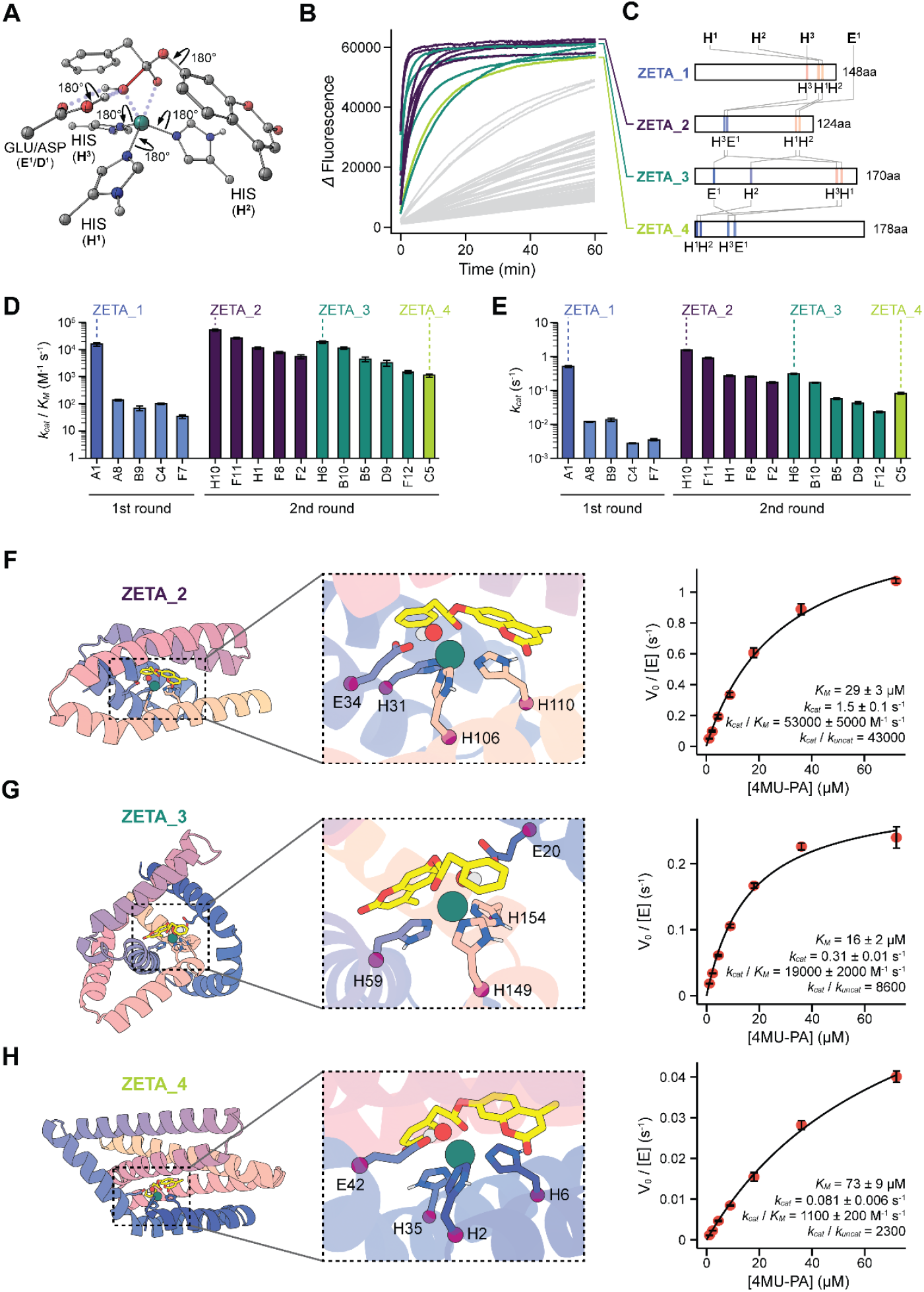
Characterization of second round designs. (**A**) Second round DFT theozymes containing the catalytic base. Zn(II) is coordinated by N_ε_ atoms in these theozymes and thus C_β_ is explicitly modeled. (**B**) Reaction progress curves colored by scaffold family. (**C**) Sequence length and catalytic residue positioning for the top design in each scaffold, named ZETA_2-4. These designs differ from each other and from ZETA_1. (**D**,**E**) Michaelis-Menten (**D**) *k*_cat_/*K*_M_ and (**E**) *k*_cat_ for the 11 second round hits with colors corresponding to scaffolds using the scheme in (B,C). Activities are higher, on average, than the first round of designs (blue); y-axes are logarithmic. (**F-H**) Design model with active site zoom-in for (**F**) ZETA_2, (**G**) ZETA_3, and (**H**) ZETA_4. Michaelis-Menton plots and parameters are on the right.

The success rate in the second design campaign was considerably higher than the first campaign (11/96 vs 1/96 designs with *k*_cat_/*K*_M_ >10^3^ M^−1^ s^−1^), supporting the conclusions from the first round analysis. The structures of ZETA_1-4 are quite different from each other and previously known metallohydrolases. The sequence positions of the catalytic residues in each of these enzymes are also very different, highlighting the diversity of RFdiffusion2 generated design solutions.

## Conclusion

We demonstrate that RFdiffusion2 can generate highly active metallohydrolases directly from active site configurations obtained by quantum chemistry calculations. The zero-shot design of an enzyme (ZETA_1) with a *k*_cat_/*K*_M_ >10^4^ M^−1^ s^−1^ – the top-ranked *in silico* design out of a total of 96 tested straight from the computer (with no experimental optimization) – is a considerable advance for de novo enzyme design which in the past has required extensive design screening and directed evolution to achieve activities at this level.^8,16^ The robustness of our design strategy is demonstrated by the zero-shot design of 3 additional enzymes from a new set of 96 designs tested straight from the computer: ZETA_2 and ZETA_3 with a *k*_cat_/*K*_M_ >10^4^ M^−1^ s^−1^, and ZETA_4 with a *k*_cat_/*K*_M_ >10^3^ M^−1^ s^−1^. ZETA_1-4 have structures very different from each other and from previously known structures. The catalytic efficiencies of ZETA_1-3 are within the range observed for native metallohydrolases (*k*_cat_/*K*_M_ = 10^4^~10^8^ M^−1^ s^−1^) and greater than all previously designed metallohydrolases prior to optimization (Table S3).^46–50^ Experimental characterization of ZETA_1 provides strong evidence to suggest that it functions as designed, utilizing a bound Zn(II) ion to activate a water molecule for nucleophilic attack and to stabilize the resulting oxyanion intermediate and flanking transition states. The PLACER and Chai-1 ensemble results suggest the key to obtaining *k*_cat_/*K*_M_ >10^4^ M^−1^ s^−1^ is precise substrate placement relative to the activated water and the Zn(II) ion; going beyond this through positioning of the general base such that it cannot reconfigure to interact with the bound ion (in the case of ZETA_1), and incorporating further sidechain oxyanion stabilization could increase activity into the range of the most active native metallohydrolases.

Our RFdiffusion2-PLACER design approach has many advantages over previous de novo enzyme design methods and should be broadly applicable for generating efficient catalysts for a wide diversity of chemical reactions. By enabling direct scaffolding of sidechain functional groups, rather than backbone N-C_α_-C=O coordinates as required in enzyme design calculations with RFdiffusion^1,28^, RFdiffusion2 bypasses the need for explicit enumeration of catalytic sidechain rotamer conformations and placement of the catalytic residues along the linear sequence–this enables each design trajectory to sample from the enormous space of possibilities rather than being confined to a small, and perhaps unrealizable, subspace. Assessment of active site preorganization and substrate/transition state positioning with PLACER and Chai-1 proved remarkably effective at identifying the most active designs; this result, together with similar observations from de novo designed serine hydrolases^1^ and retroaldolases^15^, suggests that PLACER and related deep learning approaches will be widely useful for design ranking. In our laboratory, we have found the RFdiffusion2-PLACER design approach described here to yield catalysts for several uni- and bimolecular reactions, and we look forward to seeing what the broader design community can generate with these tools which we are making freely available and open source.

## Acknowledgments

We thank Y. Wang, A. Hunt, J. Gershon, J. Min, C. Demakis, A. Shida, and Y. Kipnis for discussions about zinc hydrolases, A. Swartz, P. Lund-Anderson, N. Ennist, Y. Hsia, S. Sharma, and S. Honda for valuable discussions about metal binding, small molecule binding, and general discussions about protein design, D. Evans for guidance with performing quantum chemistry, M. Bauer for software assistance, K. VanWormer, R. Ticzon, H. Nunez-Ortega for managing the wet-lab resources, and L. Goldschmidt for maintaining the computational resources at the Institute for Protein Design. This work was funded by the National Science Foundation Graduate Research Fellowship Program, Grant No. DGE-2140004 (S.M.W), The Howard Hughes Medical Institute (D.B., I.K.), The Audacious Project at the Institute for Protein Design (A.L., N.H.), The Open Philanthropy Project Improving Protein Design Fund (I.K., S.J.P.), the ETH Zurich (D.H.), The Bill and Melinda Gates Foundation INV-010680 (D.K., W.A.), and the Schmidt Futures Foundation (A.L., S.J.P., S.S.).

## Author Contributions

S.M.W., I.K., and D.B. conceptualized the metallohydrolase design project. W.A., DT and D.B. conceptualized RFdiffusion2. W.A., J.Y., D.T., and S.S. contributed to development of RFdiffusion2. W.A. trained all versions of RFdiffusion2 used in the study. S.M.W. and I.K. performed DFT calculations. S.M.W., I.K., and D.K. developed the computational design pipeline. S.J.P. and A.L. contributed ideas, code, and functions. S.M.W. and D.K. produced and characterized the designs and analyzed the results. N.H. performed the Zn(II)-binding affinity experiments. I.K., S.J.P., A.L., and D.H. provided guidance for optimizing experimental protocols and helped interpret key results. D.B. and D.H. supervised the research. The manuscript and supplemental information were drafted by S.M.W., D.K., I.K., and D.B. All authors reviewed and commented on the manuscript.

### Data and Materials Availability

All data are available in the main text or as supplementary materials. Design scripts and RFdiffusion2 will be made available on GitHub upon publication of the manuscript. PLACER is publicly available on GitHub at the link: https://github.com/baker-laboratory/PLACER.

