## Supplementary Information for "Computational Design of Metallohydrolases"

### 1. NOMENCLATURE

Enzymes

We interchangeably refer to the best enzyme from the 1st design campaign as A1 *and* zinc metalloesterase 1 (ZETA_1) throughout the supplemental information. The name A1 came from its commercially ordered gene position in a 96-well eBlock (Well ID: A1). However, for the purposes of standardization, we would appreciate it if any future references would use **ZETA_1** as the official name.

The 2nd design campaign featured 11 major hits among 3 completely different scaffolds. After characterizing the kinetics of all 11 hits, we picked the design with the highest catalytic efficiency ($k_{cat}/K_M$) within each of the 3 scaffold families to represent them. These designs were given the names ZETA_2, ZETA_3, and ZETA_4 (ordered in terms of catalytic efficiency, where ZETA_2 has the highest). Note that ZETA_2 has the highest catalytic efficiency of every design in this manuscript, even surpassing ZETA_1.

ZETA_1-4 all have unique scaffolds made from scratch around theozymes with RFdiffusion2. As mentioned above, for the purposes of standardization, we would appreciate it if these enzymes were individually referred to as **ZETA_1**, **ZETA_2**, **ZETA_3**, and **ZETA_4** (or ZETA_1-4 if referring to all) in any future references.

Computational Protein-Ligand Ensemble Predictor Method

The name for the software that predicts ensembles of sidechain conformations and ligand conformations is now called PLACER (Protein-Ligand Atomistic Conformational Ensemble Resolver). It is publicly available on GitHub under this name. This software was formerly known as ChemNet, and its current preprint name still refers to it with this name. However, it is in the process of being published under its new name, PLACER.

### 2. GENERAL MATERIALS

General solvents and buffers

Milli-Q water was obtained in-house using a Q-POD® Ultrapure Water Remote Dispenser from Millipore Sigma. Nuclease-free water was commercially bought from Invitrogen™. Dimethylsulfoxide (DMSO) was obtained from Sigma Aldrich.

All general buffer components (i.e., HEPES, NaCl, Tris-HCl, EDTA) were bought from Millipore Sigma. All buffers were prepared in Milli-Q water and filtered through a 0.2 µm pore size Nalgene™ Rapid-Flow™ Sterile Disposable Filter Unit with a PES membrane. Further mentions of 'bottle filtration' will refer to this 0.2 µm pore size filter.

Materials for biology

Synthetic linear DNA fragments encoding the genes for each reverse-translated protein design with overhangs for golden gate assembly were ordered in a 96-well eblock® from IDT, shipped dry, and stored at -20 °C. Custom plasmid vectors were ordered from GenScript and stored at -20 °C. Golden gate assembly of the linear gene fragments into the custom plasmid vectors was done using the NEBridge® Golden Gate Assembly Kit (BsaI-HF® v2) from New England Biolabs. Competent BL21(DE3) *E. coli* cells for protein expression and competent NEB 5-alpha *E. coli* cells for plasmid production were commercially bought from New England Biolabs and stored at -80 °C. Super optimal broth with catabolite repression (SOC) media was purchased from New England Biolabs for recovery after heat-shock transformation.

Lysogeny broth (LB) media for *E. coli* inoculation was made from tryptone (Sigma Aldrich; 10 g/L), NaCl (Millipore Sigma; 10 g/L), and Bacto™ Yeast Extract (ThermoFisher Scientific; 5 g/L) dissolved in Milli-Q water. Terrific broth II (TB-II) media for protein production was bought as a solid powder from MP Biomedicals and dissolved at a concentration of 50 g/L in Milli-Q water. All media were prepared under flame, sterilized through bottle filtration, and immediately used or stored at 4 °C before use. TB-II and LB were either autoclaved or sterilized by bottle filtration before use. Kanamycin sulfate from *Streptomyces kanamyceticus* was bought as a powdered bioreagent from Millipore Sigma. Isopropyl β-D-1-thiogalactopyranoside (IPTG, dioxane-free) to induce protein expression was bought from Fisher Scientific. Agar plates with kanamycin were made in-house for colony isolation. Glycerol (≥99.0% grade) for glycerol stocks was bought from Sigma Aldrich.

Axygen® 96-well clear round bottom 2 mL polypropylene deep well plates for cultures are bought from Corning (P-DW-20-C-S). Breathe-Easier® breathable films for culture plates were bought from Andwin Scientific. Axygen® aluminum sealing films for general sealing purposes were bought from Corning (PCR-AS-600).

Materials for protein purification

Lysozyme, DNase I, and Polymyxin B sulfate used in *E. coli* cell lysis were bought from Sigma Aldrich (L6876, DN255G, P0972), respectively. Pierce® protease inhibitor tablets were bought from Thermo Scientific™. Filter polypropylene microplates with 96 wells (800 µL/well), long drip spouts, and 25 µm polyethylene frit for small-scale affinity chromatography were purchased from Agilent. Econo-Pac® 25 mL gravity flow-through chromatography columns for low-throughput affinity chromatography were purchased from BioRad. Strep-Tactin® resin for affinity chromatography was purchased from IBA Lifesciences. Desthiobiotin used in protein elution was purchased from IBA Lifesciences. Strep-Tactin® regeneration buffer containing HABA (10x concentration) was also purchased from IBA Lifesciences. Amicon® 3 kDa and 10 kDa MWCO centrifugal concentrators for protein spin-concentration were purchased from Millipore Sigma. Size exclusion chromatography was performed on an ÄKTA pure™ chromatography system from Cytiva using a Superdex 75 Increase 10/300 GL column also purchased from Cytiva.

Materials for gel electrophoresis

SDS-PAGE gel electrophoresis was performed with 15- or 26-well precast AnykD gels purchased from Bio-Rad. Tris, glycine, and sodium dodecyl sulfate were obtained from Sigma Aldrich. Laemmli buffer was purchased from Bio-Rad. Precision Plus Protein™ Dual Xtra

Prestained Protein Standards was purchased from Bio-Rad to serve as a ladder of protein molecular weight standards in gels. Electrophoresis was performed in either a Mini-PROTEAN Tetra Cell (15-well gel) or Criterion™ Cell (26-well gel), and powered by a PowerPac basic unit, all purchased from Bio-Rad. Gel staining was performed using an eStain L1 protein staining device purchased from GenScript with the eStain L1C Staining Kit also purchased from GenScript. Gels were imaged on a ChemiDoc XRS+ system purchased from Bio-Rad.

Materials for enzyme characterization

4-Methylumbelliferyl phenylacetate (4MU-PA, commercially known as phenyl-acetic acid 4-methyl-2-oxo-2H-chromen-7-yl ester) for the first round of activity screening was acquired from Millipore Sigma. 4MU-PA for the second round of activity screening and all kinetics and characterization experiments was custom synthesized by WuXi. 4-Methylumbelliferone (4MU) for making product concentration standards was purchased as a solid from Millipore Sigma. Full area 96-well flat bottom assay plates with a non-binding surface made from black polystyrene, used in the 1st round of screening measurements, were purchased from Corning. Half area 96-well flat bottom assay plates with a non-binding surface made from black polystyrene, used in all other experiments, were purchased from Corning. Fluorescence measurements were performed with The BioTek Synergy Neo2 Hybrid Multimode Reader RUO.

Zinc sulfate was purchased from Sigma Aldrich. 1,10-Phenanthroline for zinc chelation from the enzymes was purchased from Sigma Aldrich. Slide-A-Lyzer® 3.5 kDa MWCO dialysis cassettes were purchased from ThermoFisher Scientific. Mag-Fura-2, tetrapotassium salt was purchased from ThermoFisher Scientific for experiments determining zinc binding affinity. Quartz spectrophotometer cells with a pathlength of 10 mm were purchased from Starna Cells, Inc. UV-visible absorbance measurements were performed using a JASCO V-750 Double Beam UV-Visible Spectrophotometer. Quartz cuvettes (110 - Macro Cells) with a 1 mm pathlength for circular dichroism were purchased from Hellma USA Inc. Circular dichroism measurements were performed using a JASCO J-1500 Circular Dichroism Spectrophotometer.

### 3. METHODS

#### 3.1. Computational Design of Metallohydrolases

<u>Design pipeline overview</u>

We chose to design a zinc metallohydrolase for a fluorogenic ester, 4-methylumbelliferyl phenylacetate (4MU-PA), as a model reaction that would allow for sensitive screening and kinetic characterization of active enzymes and direct benchmarking with previous computationally designed zinc esterases (Fig. 1A). Rational theozymes for the rate-limiting hydroxide attack transition state in the proposed hydrolysis reaction were constructed and optimized using density functional theory. We modeled the nucleophilic attack from either side of the substrate in addition to stabilizing the oxyanion with either two external hydrogen bonds or via the zinc ion alone before creating additional low energy rotamers of the substrate to model different modes of binding (Fig. S3). Note that each theozyme contained three histidines coordinated to the zinc ion and a carboxylic acid to represent a protonated general base that deprotonated the attacking water molecule.

Protein scaffolds were generated around the designed theozymes using RFdiffusion2 with the catalytic histidine imidazoles fixed in place during generation while allowing for delta or epsilon nitrogen coordination with the zinc (Fig. S1B and Fig. S4). The general base and oxyanion stabilizing hydrogen bonds were left to be found during later sequence generation to explore additional backbone and sequence space alike due to the flexibility of residues that can accomplish these tasks. Competent scaffolds were resampled with RFjoint2 inpainting to modify loop structures and lengths outside of the active site to generate extra diversity for each scaffold (Fig. S1C). ProteinMPNN was used to generate global, foldable sequences for each scaffold which were verified by AlphaFold2 (Fig. S1D). Successfully folded designs then underwent iterative active site sequence design and constrained fast relaxing (i.e., energy minimization) with LigandMPNN and Rosetta, respectively (Fig. S1E). First, the primary sphere of residues directly interacting with the ligand (< 5Å) was redesigned before the scaffold was gently relaxed and the sidechains were repacked. Residues that were identified to make hydrogen bonds to the substrate and general bases within 3Å of the theozyme hydroxide position were kept. The process was then repeated 5 times with the allowed scaffold movement during relaxation being increasingly constrained with each cycle. Finally, the secondary sphere of residues responsible for interacting with the primary sphere residues to preorganize the active site (5Å < X < 8Å) was redesigned with LigandMPNN. The original catalytic histidines were fixed for the entirety of sequence design.

Several hundred thousand resulting design sequences across hundreds of unique scaffolds were then filtered for the presence of a general base and hydrogen bonds to the oxyanion hole for the theozymes modeled with them. Successfully filtered design sequences were then passed through AlphaFold2 to examine the predicted structure confidence and agreement with the designed structure. Structurally verified designs were then scored by ChemNet for substrate and catalytic sidechain preorganization and agreement with the designed active sites. Rosetta was used to further score the designs on additional metrics and a custom-made, weighted function combining the relevant metrics from ChemNet and Rosetta was used to identify the best sequence for every remaining unique scaffold.

Theozyme construction

The initial theozymes modeling the rate-limiting nucleophilic zinc-hydroxide attack of the 4MU-PA ester were manually constructed in a 3D chemical modeling software. Each theozyme consisted of (1) a single zinc(II) ion, (2) three imidazole rings representing histidine sidechains coordinating to the zinc ion, (3) the 4MU-PA ester substrate, (4) a hydroxide ion coordinated to the zinc ion representing the deprotonated nucleophilic water molecule, and (5) an acetic acid group interacting with the hydroxyl group, representing a protonated catalytic general base (e.g., aspartate or glutamate). Furthermore, we considered two modes for oxyanion stabilization: (1) Zn(II)-O interaction, and (2) external hydrogen bond oxyanion hole (Fig. S3 C,E). The Zn(II)-O oxyanion stabilization uses the zinc ion to stabilize the forming oxyanion in addition to stabilizing the nucleophilic hydroxide, requiring no extra elements to be modeled. On the contrary, the external hydrogen bond oxyanion stabilization has the oxyanion facing away from the zinc and uses two additional H-bonded methanol molecules to mimic a potential oxyanion hole. These two methanol molecules represent a generic form of an oxyanion hole and help locate the transition state geometry more akin to what it would look like in the presence of an oxyanion hole by polar groups in an enzyme active site. Each of these two classes of theozymes had the nucleophilic attack modeled from each side of the planar 4MU-PA ester bond before forming the chiral tetrahedral intermediate (i.e., *R* or *S* attack, Fig. S3A-B). Thus, four unique theozymes were modeled in total as transition states.

All transition state optimization calculations were performed using Gaussian 16 software.[1] Structural optimizations and frequency calculations were performed with the B3LYP-D3 method along with the 6-31G(d) basis set and the SDD ECP on the Zn(II) atom. D3 dispersion correction was applied using the Becke-Johnson damping function.[2] Solvent effects of water were included using the CPCM solvation model during geometry optimization. Frequency calculations were performed to confirm whether the structure is a minimum or a transition state. Intrinsic reaction coordinate (IRC) analysis was used to confirm that the obtained transition states connect the correct minima.

Conformational flexibility of the transition states was sampled based on rotatable bonds in the 4MU-PA substrate (Fig. S3C-F). The conformers were sampled as frozen coordinates with no geometry optimization, in order not to affect the transition state geometries. Energies of the conformers were evaluated using the GFN2-XTB semiempirical QM method.[3] This procedure was performed using a Python script *frozen-conf-xtb*, available on GitHub.[4] We generated 10 conformers for each transition state, all within 5 kcal/mol of each other, and with a minimum difference between conformers being at least 20 degrees for at least one rotatable bond. These conformers do not necessarily represent local minima in the conformational energy surface but rather capture structural diversity among low energy conformers. Depending on the specific protein environment each of these conformers may be more or less preferred, and furthermore, Rosetta minimization is capable of adjusting these rotatable bonds within a small range.

We generated unique Rosetta ligand params files from each of the transition state geometries. Each params file defines the ligand, containing the 4MU-PA substrate atoms, the Zn(II) atom, and the hydroxyl group. Rosetta ligand params file was created from the produced conformers by first converting the conformer multiXYZ files to mol2 files using Open Babel, and thereafter converting the mol2 file to params using the `$ROSETTA_MAIN/source/scripts/python/public/molfile_to_params.py` script provided in Rosetta.

<u>Theozyme conformer generation & enumeration of histidine coordination modes</u>

We used the obtained transition state geometries to construct RosettaMatch constraint files that define the six degrees of freedom between the ligand and the catalytic sidechains. For example, the interaction with histidine is defined by the ZN1-NE bond, O1-ZN1-NE angle, ZN1-NE-C angle, C2-O1-ZN1-NE torsion, O1-ZN1-NE-C torsion, and ZN1-NE-C-ND torsion. Note that 'ZN1' refers to the atom name of zinc in the structure in Fig. S4, not zinc in oxidation state 1. $N_\delta$ and $N_\varepsilon$ coordination was sampled by defining the histidine root atom (NE or ND) using the atom type NHis, which allows either of the not-protonated nitrogen atoms in either of the tautomers to be sampled. Lastly, sampling the flip of the histidine ring was enabled by setting the rotation periodicity of the O1-ZN1-NE-C torsion to 180 degrees (Fig. S4).

We used the constraint files to generate input motifs for RFdiffusion2. We used the Invrotzyme tool[5] to generate all samples of the different histidine coordination modes and flipped states. We only sampled allowed histidine placements, as defined by the constraint file, while ignoring all residue inverse rotamers (backbone position and sidechain $\chi_1$ and $\chi_2$ angles). The motifs were generated by running a command such as:

```
invrotzyme.py --cstfile theozyme.cst --params LIG.params --tip_atom …
```

<u>Backbone generation and resampling</u>

We used RFdiffusion2 to generate backbones to host each of the unique combinations of histidine placements, together with the hydrolysis reaction transition state. For simplicity, we omitted the general base at this stage and left its placement to the subsequent sequence design step. RFdiffusion2 was run by using the following command:

```
{RFdiffusion2}/run_inference.py --config-name config.yaml --config-path ./
```

Where the configuration file `config.yaml` specifies the motif parameters and other diffusion run settings. Here, `contig_atoms` specifies the fixed atoms in each input residue, with A1, B2, and C3 being the three individual histidines; PSZ is the name of the input ligand; `input_pdb` provides a path to a PDB file containing the ligand and the three histidines.

**config.yaml**

```
defaults:
  - base

diffuser:
  T: 75

guidepost_bonds: True

inference:
  align_motif: False
  num_designs: 2
  output_prefix: /path/to/RFd2_output/zn_hydrolase_PSZ_H_H_H_1_64_1_FM_model50
  input_pdb: /path/to/theozymes/PSZ_H_H_H_1_64.pdb
  final_step: 1
  ligand: PSZ
  contig_as_guidepost: True
  remove_guideposts_from_output: True
  infer_guidepost_positions: True
  guidepost_xyz_as_design: True
  recenter_xt: True
  recenter_xt_dropt: 3

contigmap:
  contigs: ['130-275,A1-1,B2-2,C3-3']
  length: '130-275'
  contig_atoms: '{"A1":"NE2,CD2,CG,CB,ND1,CE1", "B2":"NE2,CD2,CG,CB,ND1,CE1",
                "C3":"NE2,CD2,CG,CB,ND1,CE1"}'
```

From the generated protein backbones we selected globular proteins with desirable binding pocket features based on metrics describing ligand burial, substrate exposedness, lack of clashes between the ligand and protein backbone, loop content, distance of termini from the ligand, radius of gyration, and the length of the longest helix.

We further diversified identified protein backbones with RoseTTAFold joint inpainting (RFjoint2)[6] by re-generating loop regions of the diffused backbones with variable length loop substitutions. The loop regions were selected based on geometric distance from the ligand heavy atoms and the motif residue. Sequence design was enabled on all non-motif positions during the inpainting process. For each input scaffold, at least 4 diversified backbones were generated with both of these methods.

Scaffold sequence design

We reasoned that due to the possible flexibility and sequence diversity in the enzyme active site, it would be beneficial to first identify sequences that are more likely to fold correctly, and only then focus on designing the protein-ligand interactions. Therefore, we first used ProteinMPNN[7] to generate amino acid sequences optimal for each of the selected backbones. We generated 20 sequences for each backbone while keeping the catalytic histidines fixed. Three sampling temperatures (T = 0.1, 0.3, 0.6) were used, with 10 sequences generated for T = 0.1, 5 sequences generated for T = 0.3, and 5 sequences generated for T = 0.6. This was performed while omitting the introduction of cysteine and methionine residues (--omit_AA CM) and down-weighting the introduction of alanine residues (--bias_AA A: -0.75). Generated sequences were analyzed using single-sequence AlphaFold2 prediction (monomer pTM model 4 with 6 recycles) to identify the sequences that do fold into the desired shape. Successful sequences were first selected based on AF2 metrics of pIDDT ≥ 60, $C_\alpha$ RMSD ≤ 3.0 Å, PAE ≤ 15, pTM ≥ 0.5, and recycle convergence ≤ 1.5 Å.

The sequences that passed these AF2 filters were then further processed and filtered for their catalytic histidine positional agreement with the design models. Selected AF2-predicted models were superimposed to the corresponding design models to calculate the Euclidean distances between specific pairs of atoms on matching catalytic residues (e.g., the distance between AF2 H130 $C_\alpha$ - design model H130 $C_\alpha$). We calculated RMSDs from these measured distances for provided sets of atoms in each catalytic residue (e.g., RMSD of H130 $C_\alpha$ + $C_\beta$) before calculating the RMS between these RMSDs for all the catalytic residues of one type (e.g., RMS of $C_\alpha$ + $C_\beta$ RMSDs for H118, H130, and H134). An example command calculating the backbone atom RMSD (`bb_rmsd`), backbone + $C_\beta$ RMSD (`cb_bb_rmsd`), imidazole ring RMSD (`imidazole_rmsd`), and $C_\alpha$ + $C_\beta$ RMSD (`ca_cb_rmsd`) for each matching catalytic histidine is given below:

```
sidechain_rmsd_and_info_af2_matching_res.py --scorefile scores.sc --ref_pdbs_dir
af2_inputs/ --params LIG.params --output_file updated_scores.sc
--atom_groups '{"HIS": [{"atoms": ["N", "CA", "C", "O"], "label": "bb_rmsd"},
{"atoms": ["N", "CA", "C", "O", "CB"], "label": "cb_bb_rmsd"}, {"atoms": ["CG", "CE1",
"ND1", "NE2", "CD2"], "label": "imidazole_rmsd"}, {"atoms": ["CA", "CB"], "label":
"ca_cb_rmsd"}, {"atoms": ["ND1", "NE2"], "label": "imid_sc_nitrogen_rmsd"}]}'
```

After processing with the above script, the sequences were required to pass at least one additional filter of catalytic histidine backbone (or +$C_\beta$) (`bb_rmsd` or `cb_bb_rmsd`) RMS of RMSDs ≤ 2.0 Å, catalytic histidine $C_\alpha$ + $C_\beta$ (`ca_cb_rmsd`) ≤ 2.0 Å, catalytic histidine imidazole (`imidazole_rmsd`) RMS of RMSDs ≤ 3.0 Å, or catalytic histidine imidazole tip nitrogens (`imid_sc_nitrogen_rmsd`) RMS of RMSDs ≤ 3.0 Å.

Iterative active site sequence design with FastMPNNdesign

To optimize the amino acid sequence around the substrate and the catalytic residues we used a protocol of iteratively performing ligandMPNN[8] sequence generation, Rosetta sidechain packing, and backbone minimization, with an implementation analogous to Rosetta FastDesign[9], hereby referred to as FastMPNNdesign. The designable positions were selected based on distance from the ligand (substrate, Zn(II), OH) heavy atoms; residues with $C_\alpha$ atom within 8 Å from any ligand heavy atom were considered for design. Geometric constraints were applied to the Zn(II)-HIS interactions during repacking and minimization using the Rosetta AddOrRemoveMatchCsts mover. To facilitate the introduction of a general base near the Zn(II)-bound hydroxide, we identified backbone positions within 5.0 Å from the hydroxyl proton, or 7.0 Å if the distance between the proton and $C_\beta$ was less than that of $C_\alpha$, and applied an increased bias (+4.0) towards GLU, ASP, and HIS at those positions during ligandMPNN sequence generation.

#### Analysis and filtering of designed sequences

Designs were selected based on metrics describing protein-ligand contacts (Rosetta ddG, contact molecular surface, SASA, presence of a general base, H-bonds to/from acceptors/donors). An additional round of ligandMPNN sequence design was performed at the positions flanking the binding pocket, with only those in the window of 9-13 Å from the ligand enabled for redesign. For each successful design, we generated 10 additional sequences at temperatures 0.1 and 0.2, with 5 sequences coming from each temperature.

The structures of the resulting sequences were predicted using AlphaFold2 (monomer pTM model 4 with 6 recycles) and filtered with AF2 metric cutoffs set at plDDT ≥ 72.5, $C_\alpha$ RMSD ≤ 1.75 Å, PAE ≤ 7.25, pTM ≥ 0.725, and recycle convergence ≤ 0.75 Å. The passing structures were further processed and filtered for their catalytic histidine positional agreement with the design models using the `sidechain_rmsd_and_info_af2_matching_res.py` script as described above. After processing with the script, the sequences were required to pass at least one filter of catalytic histidine backbone (or +$C_\beta$) (`bb_rmsd` or `cb_bb_rmsd`) RMS of RMSDs ≤ 0.85 Å, catalytic histidine $C_\alpha$ + $C_\beta$ (`ca_cb_rmsd`) ≤ 0.85 Å, catalytic histidine imidazole (`imidazole_rmsd`) RMS of RMSDs ≤ 1.25 Å, or catalytic histidine imidazole tip nitrogens (`imid_sc_nitrogen_rmsd`) RMS of RMSDs ≤ 1.25 Å.

The remaining sequences then had their *apo*-AF2 predicted structures relaxed with Rosetta FastRelax,[10] before calculating the spatial aggregation propensity (SAP) score, net charge, and hydrophobic exposure SASA of the relaxed structures. The relaxed sequence designs were then filtered for their 'expressibility' where they were first required to have SAP score ≤ 75 or the net charge excluding histidines ≥ 20 and hydrophobic exposure SASA ≤ 1500. The passing sequences were then clustered by their original design structure and sequences containing repeating amino acids occurring 5 times or more in a row and/or hydrophobic residues in the set AVILMFYWP occurring 7 times or more in a row were removed. For each cluster, at least one sequence was kept by adding some leniency to the aforementioned selection if all sequences were flagged by keeping the sequence with the highest absolute AF2 pTM to AF2 $C_\alpha$ RMSD ratio.

#### Enzyme sidechain ensemble analysis with ChemNet

The AF2-predicted models of selected designs were superimposed to their corresponding design models, the ligand placed into the AF2 model, and the resulting *holo*-structures relaxed with Rosetta FastRelax. Subsequent ChemNet input files were prepared either by (1) removing the zinc and hydroxide, or (2) removing the hydroxide and substrate. Thus, ChemNet ensemble predictions were performed on the substrate-enzyme and zinc-enzyme complexes for each design. The generated 40 ChemNet models were analyzed for the following metrics: average substrate RMSD, average histidine RMSD, average histidine pRMSD, average pocket residue RMSD and pRMSD, and worst pocket residue pRMSD. Furthermore, we analyzed the 5 highest pLDDT (1D confidence) frames for substrate RMSD and average plDDT.

Filtering and analysis after ChemNet

Dataframes containing scores from FastMPNNdesign, AF2, Rosetta, and ChemNet were merged based on design identity for the few thousand remaining designs spanning approximately 130 unique RFdiffusion2 scaffolds. First, one simple random design sequence for every candidate RFdiffusion2 scaffold was manually screened to exclude implausible scaffolds (e.g., completely buried substrate, buried coumarin, and exposed phenylacetate). Thereafter, we used a broad combination of redundant and nonredundant *in silico* metrics to rank and identify the best design(s) for each remaining scaffold. This approach was taken as opposed to using strict cutoffs due to the lack of known standalone predictive metrics for metallohydrolase activity; additionally, we wanted to maximize structural diversity in the final set of 96 designs as opposed to sequence sampling, hence the clustering by RFdiffusion2 scaffold. Thus, we sought to explore diverse ranges for each metric in our final order while also capturing designs that were hypothesized to be most well-balanced.

To this end, the remaining designs were ultimately clustered by the parent RFdiffusion2 scaffold. After clustering, a selection of AF2, ChemNet, and Rosetta metrics hypothesized to be potentially relevant indicators of preorganization, active site configuration, and predicted structure confidence were normalized and used for a custom-weighted scoring function to score and rank designs within each cluster. Note that the metrics were split accordingly before normalization into groups where higher metric values are preferred (e.g., AF2 plDDT) or lower metric values are preferred (e.g., AF2 $C_\alpha$ RMSD). The chosen weights were an arbitrary initial guess that were meant to roughly balance the cumulative influence of grouped metrics from AF2, ChemNet, and Rosetta. The 'composite score' calculated as the output of the scoring function was simply the weighted sum of all normalized metrics preferring lower values, plus the weighted sum of inverse-normalized metrics preferring higher values. Thus, the lowest composite score corresponded to the best-ranked design.

By this logic, the 3 designs with the lowest composite scores were selected from each RFdiffusion2 scaffold cluster (for clusters with 3 or fewer candidate sequences, all were retained), applying the metrics and weights shown below. All other designs were filtered out. Following this filtering step there were only 3 or fewer sequences left for each candidate scaffold, leaving approximately 260 sequences spanning 119 RFdiffusion2 scaffolds. The optimal sequence for each RFdiffusion2 scaffold cluster was chosen as the one with the lowest AF2 $C_\alpha$ RMSD from its design. All other sequences were filtered out, leaving a single sequence per scaffold. Only scaffolds with top-ranked sequences among the 96 lowest AF2 $C_\alpha$ RMSD values were subsequently ordered.

```
## Columns to Normalize and Invert (i.e., higher value = better) ##
columns_to_normalize_and_invert = [
'pTMscore_af2_out','zinc_plddt_pde_top5','zinc_plddt_top5','zinc_plddt_pde_pnear',
'zinc_plddt_pnear','subst_plddt_top5','subst_plddt_pde_top5',
'subst_frac_good_kabsch', 'subst_plddt_pnear', 'subst_plddt_pde_pnear',
'O2_hbond_fastmpnn','total_hbonds', 'O2_hbond', 'O3_hbond', 'O5_hbond',
'electrostatic_complementarity_p', 'electrostatic_complementarity_s',
'hbonds_to_metal_binding_residues']
...
```

```
normalized_df[col] = 1 - (input_df[col] - min_val) / (max_val - min_val)

## Columns to Normalize without Inversion (i.e., lower value = better) ##
columns_to_normalize = [
'rmsd_to_input_af2_out', 'zinc_rmsd_ligand', 'zinc_rmsd_std_cat_rms',
'subst_rmsd_ligand', 'subst_u_cat_rms', 'zinc_rmsd_std_ligand',
'zinc_rmsd_pde_top5', 'zinc_rmsd_top5', 'zinc_rmsd_cat_rms', 'zinc_u_cat_rms',
'subst_rmsd_std_ligand', 'subst_rmsd_top5', 'subst_rmsd_pde_top5',
'subst_rmsd_cat_rms', 'subst_rmsd_std_cat_rms', 'SAP_score', 'violations',
'all_cst_fastmpnn', 'cat_HIS_ca_cb_ONLY_rms_of_rmsds_af2_out',
'cat_HIS_imidazole_rms_of_rmsds_af2_out',
'cat_HIS_imidazole_tip_Ns_rms_of_rmsds_af2_out', 'ligand_rmsd_post_holo_relax',
'Ca_rmsd_after_fastrelax', 'longest_cont_apolar_seg',
'hydrophobic_exposure_sasa_in_design', 'fa_elec_fastmpnn', 'fa_intra_elec_fastmpnn',
'hbond_sc_fastmpnn', 'total_score_fastmpnn']
...
normalized_df[col] = (input_df[col] - min_val) / (max_val - min_val)
```

```
## Composite Score Calculation ##
# Each cumulative family of metrics (i.e., Rosetta, ChemNet, and AF2) is
# approximately equal.

weights = {
    ## AF2 Metrics ##
    'rmsd_to_input_af2_out': 0.125,
    'pTMscore_af2_out': 0.125,
    'cat_HIS_ca_cb_ONLY_rms_of_rmsds_af2_out': 0.0625,
    'cat_HIS_imidazole_rms_of_rmsds_af2_out': 0.0625,
    ## Zinc-Enzyme ChemNet Ensemble Metrics ##
    'zinc_rmsd_ligand': 0.05,
    'zinc_rmsd_std_ligand': 0.01,
    'zinc_plddt_pde_top5': 0.01,
    'zinc_rmsd_pde_top5': 0.01,
    'zinc_rmsd_top5': 0.01,
    'zinc_plddt_top5': 0.01,
    'zinc_rmsd_cat_rms': 0.01,
    'zinc_rmsd_std_cat_rms': 0.01,
    'zinc_u_cat_rms': 0.01,
    'zinc_plddt_pde_pnear': 0.01,
    'zinc_plddt_pnear': 0.01,
    ## Substrate-Enzyme ChemNet Ensemble Metrics ##
    'subst_rmsd_ligand': 0.05,
    'subst_rmsd_std_ligand': 0.01,
    'subst_rmsd_top5': 0.01,
    'subst_plddt_top5': 0.01,
    'subst_rmsd_pde_top5': 0.01,
    'subst_plddt_pde_top5': 0.01,
    'subst_rmsd_cat_rms': 0.01,
    'subst_rmsd_std_cat_rms': 0.01,
    'subst_frac_good_kabsch': 0.01,
    'subst_plddt_pnear': 0.01,
    'subst_plddt_pde_pnear': 0.01,
    ## Rosetta Metrics Dealing with Expressibility & Active Site Features ##
    'O2_hbond_fastmpnn': 0.05,
    'SAP_score': 0.05,
    'violations': 0.05,
    'all_cst_fastmpnn': 0.05,
    'longest_cont_apolar_seg': 0.025,
    'hydrophobic_exposure_sasa_in_design': 0.025,
    ## Rosetta Metrics from FastMPNNdesign ##
    'ligand_rmsd_post_holo_relax': 0.05,
    'Ca_rmsd_after_fastrelax': 0.05,
```

```
    'O2_hbond': 0.025,
    'O3_hbond': 0.025,
    'O5_hbond': 0.015,
    'electrostatic_complementarity_p': 0.015,
    'electrostatic_complementarity_s': 0.015,
    'hbonds_to_metal_binding_residues': 0.015,
    'fa_elec_fastmpnn': 0.01,
    'fa_intra_elec_fastmpnn': 0.01,
    'hbond_sc_fastmpnn': 0.01,
    'total_score_fastmpnn': 0.01
}

## Calculate the Composite Score ##
normalized_df['composite_score'] = sum(normalized_df[col] * weight for col, weight
in weights.items())
...
### FILTERING TO 3 FILES PER GROUP IF MORE THAN 3 ###
filtered_groups = []
for group, data in pre_filtered_df.groupby('flow_matching_group'):
    if len(data) > 3:
        data = data.nsmallest(3, 'composite_score')
```

Structural similarity search of the PDB and AFDB

To evaluate the structural novelty of the A1 (ZETA_1) design, we employed FoldSeek,[11] a tool that enables rapid and accurate structure searches. We compared the A1 design against the PDB100, a clustered subset of the Protein Data Bank containing entries deposited before January 2024, and the AFDB50,[12] a clustered database with 50% sequence identity (version 4, encompassing 54 million structures). We reported the TM-score for the top match. The top hit against PDB100 was 8GYN, which corresponds to the Tumor Necrosis Factor Alpha-Induced Protein 8-Like Protein 1 in zebrafish. In contrast, the top hit against AFDB50 was A0A176QCW7, an unknown protein predicted to function as a biotin transporter.

#### 3.2. Changes in the 2nd Design Campaign of Metallohydrolases

Note: This is referred to as the "2nd/second design campaign" and "round 2" interchangeably in the main text and supplemental information.

Observations and trends from the 1st design campaign

We made several observations from the design structures of the hits from the 1st design campaign which informed our refined strategy for the 2nd design campaign. We describe these observations below:

(1) All of the hits except for F7 came from the class of DFT theozymes using Zn(II) to stabilize the oxyanion, rather than 2 protein hydrogen bonds. Perhaps this was because it was difficult to precisely position 2 protein hydrogen bonds, either from sidechains and/or backbones, to the substrate oxyanion hole during sequence design after the scaffold had already been generated. Even in the case of F7, the design model only predicts 1 hydrogen bond from a serine residue to the oxyanion, not 2 as intended by the DFT theozyme. We allowed these cases, like F7, to pass since it was extremely rare to get a well-behaved design with 2 hydrogen bonds in place stabilizing the oxyanion.

(2) Excluding F7, the 4 hits using Zn(II) to stabilize the oxyanion were derived from the DFT theozyme of the *S*- enantiomer of the hydroxide attack. That is, the design models of A1 (ZETA_1), A8, B9, and C4 all contain the *S*- enantiomer of the hydroxide attack. F7 models the *R*- enantiomer of the hydroxide attack, but note that it was designed for a different mode of oxyanion stabilization.

(3) The design models of every hit coordinate the Zn(II) with the $N_{\varepsilon}$ atom of the His residues, with the exception of B9 which has 2/3 His residues coordinating Zn(II) with their $N_{\varepsilon}$ atom and the other His residue coordinating with its $N_{\delta}$ atom.

(4) The scaffold for A1 (ZETA_1) was generated around its theozyme with the exact conformation from DFT (i.e., NOT from one of the extra conformers rapidly generated from the original DFT model using GFN2-XTB semiempirical QM). However, during FastMPNNdesign, Rosetta chose a different conformer with the 4MU flipped approximately 180 degrees. Nonetheless, this is almost an equivalent geometry in terms of the interactions that were optimized during DFT such as pi-stacking with the His residues. The models of the other hits also had the substrate approximately in the geometry that was optimized out of DFT, rather than one of the more drastically-different conformers. We interpreted this to suggest that designing around the exact DFT theozyme, instead of a different conformer of the DFT theozyme, is optimal and that any conformational sampling should be done at the DFT step (i.e., creating different conformer input models for DFT).

(5) The position of the proposed general base in the design models of the hits was suboptimal in comparison to how it was positioned during DFT or in comparison to native enzymes like astacin, thermolysin, or carboxypeptidase. Additionally, we experimentally

confirmed that the general base was dysfunctional in A1 (ZETA_1) after knocking it out to alanine and measuring the kinetics, which were not significantly different from the wild-type (Fig. 3I). Although we modeled the general base during DFT, we did not include it in the motif during RFdiffusion2, reasoning that Glu/Asp/His could accomplish this function and thus it could be found during sequence design. This suggested to us that this strategy was not ideal and that the position of the general base was potentially more delicate than expected.

(6) The sequence lengths of the hits were all relatively short compared to what was sampled during design. We generated scaffolds that were between 130 amino acids and 275 amino acids in length, but 4/5 of the hits were designed to be <175 amino acids in length (A8 was only 206 amino acids). Notably, A1 (ZETA_1) was the shortest enzyme of the hits, only 148 residues in sequence length. Many of the methods may perform better with smaller proteins which could have biased the set of proteins that survived the design pipeline to be smaller on average. Nonetheless, the ideal catalyst should generally be as small as possible to optimize material efficiency, assuming the desired catalytic properties of the catalyst stay the same.

<u>Theozyme preparation for the 2nd design campaign using only new DFT models</u>

We used the observations stated above to narrow our theozyme search in a second round of DFT and enhance our design pipeline for the 2nd design campaign. Accordingly, we restricted our new DFT transition state search to: (1) theozymes in which the oxyanion is stabilized by Zn(II); (2) the *S*-enantiomer of the hydroxide attack; and (3) histidine sidechains explicitly coordinating Zn(II) through their $N_\varepsilon$ atom, thus adding $C_\beta$ to each histidine in the DFT input models (Fig. 4A). We built an DFT input theozyme of this description using a 3D chemical modeling software, with the A1 (ZETA_1) design model active site as reference, and performed transition state optimization as described in section 3.1.

Additionally, we wanted to perform any conformational sampling at the DFT step, where unique conformers could be optimized and verified to be a valid transition state, especially since we were working with only one specific theozyme. We modeled every combination of 180° flips for (1) each histidine sidechain, (2) the 4MU leaving group in the substrate, and (3) the carboxylate general base (see annotated arrows on Fig. 4A) and performed transition state optimization for each model. In total, 32/32 optimized transition state geometries were successfully obtained from DFT for the combinations of conformational samples. These optimized transition state geometries from DFT were directly turned into pdb files. Furthermore, the general base was represented as either glutamate or aspartate. This amounted to 64 pdb theozymes which served as the inputs to RFdiffusion2.

<u>Generating scaffolds with an updated version of RFdiffusion2</u>

The updated version of RFdiffusion2 can optionally intake point coordinates which direct the model on where it should aim to generate the scaffold's center of mass. These coordinates are specified as an 'ORI token' in the theozyme input pdbs, a unique line with a format such as:

```
HETATM   80  ORI ORI X  10      X_CORD  Y_CORD  Z_CORD  1.00  1.00           X
```

Where `X_CORD`, `Y_CORD`, and `Z_CORD` specify the point coordinate (x,y,z) of the ORI token directing RFdiffusion2 on where the generated protein center of mass should be approximately located. The advantage of this strategy is that a user can place ORI tokens around parts of the substrate or theozyme which they believe should be more surrounded by protein, effectively controlling which part of the theozyme should be more buried (and conversely, which part of the theozyme should be more solvent-exposed). We suspected that it might be important for the 4MU leaving group to be more solvent exposed, suggesting that the ORI token be placed around the PA part of the substrate, but we decided not to fully commit to that strategy in favor of one that would yield a greater diversity of scaffold designs.

To this end, we placed a central ORI token near the ester of the substrate and used a simple script to generate a cloud of 71 additional ORI token coordinates in a 4.5 Å radius around the central point, totaling 72 ORI tokens. The ORI tokens were approximately homogeneously distributed throughout the sphere, not just on the surface. The result was a wide diversity of ORI tokens which would specify many points for generating protein centers of mass with respect to the theozyme, resulting in many different binding orientations. This ended up being the case for ZETA_2 and ZETA_3, which had ORI tokens on opposite sides of the substrate (Fig. S24).

We performed small-scale RFdiffusion2 testing of these 72 ORI token positions with a single theozyme and manually checked the outputs. We decided to remove 8 of the ORI tokens, although in retrospect we were not convinced that this step was worth the computational effort. This left us with a final set of 64 ORI tokens. This set of filtered ORI tokens was inserted the 64 theozyme pdbs; each theozyme pdb file was replicated for every ORI token, where a single ORI token was inserted into each copy, generating the complete set of combinations between the 64 theozyme pdbs and all ORI tokens. These served as the final inputs to RFdiffusion2. Specifically, there were 64 theozymes ($2^5$ flip conformer combinations x 2 possible general bases (Glu/Asp)) x 64 filtered ORI tokens = 4096 unique combinations as inputs to RFdiffusion2.

RFdiffusion2 was set up and executed as described in section 3.1, with some minor changes to the yaml file (example shown below). Notably, we specified the functional group for the general base as part of the motif in this 2nd second design campaign, instead of leaving the general base to be found during sequence design. We hypothesized that this would better guarantee its precise positioning since it was being fixed in its DFT geometry from the start of the pipeline. Additionally, we reduced the possible scaffold length range from 130–275 to 110–190 amino acids. Specifically, for each of the 4096 RFdiffusion2 inputs, commands were created such that exactly 2 structures would be produced: 1 structure with a length between 110-150 amino acids and 1 structure with a length between 150-190 amino acids. Thus, a total of ~8000 unique scaffolds were generated.

**config.yaml (2nd Design Campaign with Updated RFdiffusion2)**

```yaml
defaults:
  - aa_v2

diffuser:
  T: 100
  aa_decode_steps: 40

guidepost_bonds: True

inference:
  align_motif: False
  num_designs: 1
  output_prefix: /path/to/RFd2_output/theozyme_X_GLU_lig_SZA_ORI_Y__model_RFD140
  input_pdb: /path/to/theozymes/theozyme_X_GLU_lig_SZA_ORI_Y.pdb
  final_step: 1
  ligand: SZA
  contig_as_guidepost: True
  idealize_sidechain_outputs: True
  remove_guideposts_from_output: True
  infer_guidepost_positions: True
  guidepost_xyz_as_design: True
  center_motif: True
  recenter_xt: False
  update_seq_t: False
  write_trb: True

contigmap:
  contigs: ['110-150,A1-1,B2-2,C3-3,D4-4']
  length: '110-150'
  contig_atoms: '{"A1":"NE2,CD2,CG,CB,ND1,CE1", "B2":"NE2,CD2,CG,CB,ND1,CE1",
                 "C3":"NE2,CD2,CG,CB,ND1,CE1", "D4":"OE1,OE2,CG,CD"}'
```

Filtering with Chai-1 predictions of the protein-Zn(II)-substrate phosphoanalog

We had the advantage of incorporating Chai-1 as a computational filter in our 2nd design campaign, encouraged by the remarkable agreement between the A1 (ZETA_1) design model and the predicted Chai-1 ensembles, which also predicted the D67 and H130 Zn(II) binding competition. However, we had concerns about using Chai-1 to predict the protein-Zn(II)-substrate ensemble and filtering on this basis: this strategy could enrich design selection for tight substrate binders, which are not necessarily good enzymes. Enzymes perform catalysis through binding their reaction transition state(s) tighter than the substrate or product(s). Therefore, it would be ideal to use Chai-1 to predict ensembles of the protein-Zn(II)- and something that resembles the transition state of the reaction, then filtering on this basis. However, we were also concerned about using Chai-1 to predict the protein-Zn(II)-tetrahedral carbon intermediate ensemble since it will have never seen an ester tetrahedral during training because it is an unstable, brief intermediate and therefore non-crystallizable.

Thus, we decided that the ideal Chai-1 ligand prediction would (1) be nearly identical to our substrate, (2) resemble the transition state geometry and charge, and (3) be well-represented in its training data, especially in the context of Zn(II). Indeed, crystallographers and enzymologists previously solved problems (1) and (2) when trying to capture metallohydrolase structures in their catalytic geometry[29] and measure the binding affinity of the transition state: they used phosphate transition state analogs (phosphoanalogs) of the ester or amide substrate, typically phosphodiesters. Phosphates are tetrahedral, and phosphodiesters specifically, typically have a net negative charge of -1 around neutral pH, closely mimicking the geometry and charge of the transition state of a hydrolysis reaction in a metallohydrolase active site. Furthermore, there are many crystal structures of zinc hydrolases bound to these transition state analogs (e.g., PDB: 1TLP (pdb_00001tlp), 1QJI (pdb_00001qji), 5OD1 (pdb_00005od1), 2TMN (pdb_00002tmn)) because of their resemblance of the transition state, making them potent inhibitors, solving problem (3). With this information, we made Chai-1 predictions of the protein-Zn(II)-phosphoanalog of 4MU-PA (i.e., replacing the ester carbon with phosphorus) and filtered the designs from these predictions. An example of a Chai-1 command used is shown below:

```
path/to/chai/run_chai.sh --name designX --protein "['DESIGNSEQ']" --ligand "['[Zn+2]',
'Cc2cc(=O)oc3cc(OP(=O)([O-])Cc1ccccc1)ccc23']" --output_dir /path/to/out/designX
--export_arrays --export_mode json
```

Predictions and filtering was performed in two broad iterations: (1) Chai-1 predictions of the protein-Zn(II)-phosphoanalog of 4MU-PA without additional N-terminal or C-terminal tags followed by filtering, and (2) Chai-1 predictions of the protein-Zn(II)-phosphoanalog of 4MU-PA with an added MSG N-terminal sequence and added GSAWSHPQFEK C-terminal sequence, representing what the designs would be expressed as, followed by filtering. The second step was performed due to concerns about the N- or C-terminus being too close to the active site in some designs. After each step of predictions, filtering was also done in two iterations: (1) general metric cutoffs that every design had to pass with no exceptions and (2) more extreme metric cutoffs that every design had to pass unless every design from a unique scaffold family was filtered out, in which case only the best designs from that family would be kept (2-3 maximum).

Although the exact filtering cutoffs differed for the steps described above, we always filtered on the same metrics. Described herein are the metrics and the specific cutoffs for the final step of filtering. First, we filtered for the positional agreement between the Chai-1 predictions and the design model: $C_\alpha$ RMSD ≤ 1.25 Å, catalytic residue RMSD ≤ 1.5 Å, substrate RMSD ≤ 4.0 Å, substrate "core" atoms (ester + hydroxide) RMSD ≤ 2.0 Å, and Zn(II) RMSD ≤ 1.25 Å. Note that Chai-1 predicts 5 ensembles by default; we filtered such that 4/5 (80%) of the Chai-1 predictions had to pass all of these filters. Additionally, we filtered on the Chai-1 confidence metrics for its predictions: average protein atom (chainA) pLDDT ≥ 80, average substrate atom (chainB) pLDDT ≥ 85, Zn(II) (chainC) pLDDT ≥ 85, catalytic residue atom pLDDT ≥ 85, substrate "core" atoms (ester + hydroxide) pLDDT ≥ 80, ipTM ≥ 0.7, pTM ≥ 0.875, protein-substrate average PAE (chainA_chainB_pae_avg) ≤ 6, protein-Zn(II) average PAE (chainA_chainC_pae_avg) ≤ 6, and finally substrate-Zn(II) average PAE (chainB_chainC_pae_avg) ≤ 6. Again, note that Chai-1 predicts 5 ensembles by default and we required 4/5 (80%) of the Chai-1 predictions had to pass all of these filters. At this step, if all designs were filtered out for a scaffold family, we kept the designs with the lowest substrate RMSD and catalytic residue RMSD in that scaffold family (2 maximum). This left us with designs spanning 37 unique scaffolds.

#### 3.3. Experimental Biology & Biochemistry

Cloning and transformation

Genes encoding the designed proteins with overhangs for Golden Gate assembly were ordered from IDT as dry eBlocks™. The DNA sequences were codon-optimized for increased *E. coli* expression, and ordered as follows:

```
(5') ATACTACGGTCTCAAGGA-<design>-GGTTCCCGAGACCGTAATGC (3')
```

The DNA fragments were cloned into a custom vector (pDTstrep1; 5684 base pairs) encoding a kanamycin-resistance gene for positive selection, a T7 promoter, and a C-terminal Strep-tag®, following a published protocol.[13] The final expressed proteins were obtained as `MSG-design-GSA-WSHPQFEK`, where WSHPQFEK is the C-terminal Strep-tag® sequence for Strep-Tactin® affinity chromatography purification.

Before golden gate assembly, 20 μL of nuclease-free water was added to each well of the dry eBlock before the plate was covered with an aluminum seal, vortexed, and centrifuged (1000g x 30s) to suspend the linear DNA fragments at a concentration of 10 ng/μL (approximately 23 nM for the largest fragments of 700 bp). Golden gate assembly was performed using the NEBridge® Golden Gate Assembly Kit (BsaI-HF® v2) containing (1) a solution of BsaI type IIS restriction enzyme and T4 DNA ligase, and (2) a T4 ligase buffer with 1 mM ATP. The reactions were set up at a 5 μL scale such that DNA fragment concentration was at least 8.8 nM for the largest DNA fragments (700 bp) and the cloning vector concentration was 4.4 nM. First, a master mix of the cloning reaction components (GG assembly kit, T4 buffer, cloning vector, and $ddH_2O$) was prepared. For each 5 μL reaction, the master mix consisted of 1.8 μL ddH2O, 0.5 μL T4 buffer, 0.25 μL GG assembly kit, and 0.54 μL cloning vector (from 41 nM stock), mixed in the given order. The master mix was prepared for 96 reactions in a cooled microcentrifuge tube and then spread out in 3.1 μL aliquots to a cooled PCR plate. Lastly, 1.9 μL solution of the DNA fragment (~23 nM stock in case of 700 bp fragment) was added to each well and carefully mixed with the pipette tip. The plate was sealed, and incubated in a thermocycler (37 °C) for 20 minutes, followed by a 5 minute incubation at 60 °C to heat-inactivate the enzymes. The resulting plasmid products of Golden Gate assembly were then used for transformation; excess reaction mixture was stored at -20 °C for future use.

Bacterial transformation was performed over ice using competent BL21(DE3) *E. coli* cells. A new PCR plate was set into a cooled metal block on ice and 2.0 μL of the Golden Gate plasmid products from the previous steps were transferred into their respective wells with a multichannel pipette. This PCR plate was briefly sealed and centrifuged (1000g x 30s) to get the plasmids at the bottom of each well. Next, tubes of competent *E. coli* stored at -80 °C were allowed to thaw and swiftly added in 7 μL aliquots to each well with a repeater pipette. The PCR plate was rapidly centrifuged (100g x 5-10s) to ensure sufficient mixing of the cells and plasmids at the bottom of each well and the plate was returned to the metal cooling block on ice for 30 minutes. Thereafter, the plate was transferred to a 42 °C metal block for 17~20s to heat shock the *E. coli* and returned to ice for 2 minutes. 100 μL of pre-warmed SOC recovery media was added into each well of the PCR plate and a breathable film was used to cover the plate. The plate was incubated in a microplate shaker (37 °C, 1200 rpm) for 1 hour. 900 µL aliquots of sterile LB media with kanamycin (50 µg/mL) were added to each well in a 2 mL deep round-well incubation plate under flame, followed by 100 µL of the transformed *E. coli* in SOC media. The deep round-well plate was covered with a breathable film and incubated in a microplate shaker (37 °C, 1200 rpm) for 16-18 hours.

Successfully cloned designs had opaque wells while unsuccessfully cloned designs had transparent wells the following day. Glycerol stocks were made for every clone by mixing 125 µL of 50% v/v glycerol in Milli-Q water with 125 µL of the overnight culture in a new PCR plate. The PCR plate was covered with an aluminum seal and stored at -80 °C. Note that the obtained clones are not sequence-verified at this stage and could be polyclonal or even wrong due to mistakes during DNA synthesis. After preparing the glycerol stocks, the remaining cultures were used for expression, as described below.

Small-scale protein expression with IPTG

Proteins were expressed at a 4x1 mL culture scale in sterile terrific broth II (TB-II) media supplemented with 50 µg/mL of kanamycin. Four 2 mL deep round-well plates were loaded with 1 mL of the TB-II+kanamycin media under flame. 150 µL of the overnight cultures in LB media were transferred into each respective well for each of the four plates. These plates were covered with breathable film and incubated in a microplate shaker (37 °C, 1200 rpm) for 90 minutes to promote sufficient cell growth. Thereafter, 11.5 µL of 100 mM IPTG (isopropyl β-D-1-thiogalactopyranoside) was added into every well of the four plates (1 mM final [IPTG]) under flame to induce expression. The plates were resealed with the breathable films and incubated in the microplate shaker (37 °C, 1200 rpm) for 2 hours before being transferred to a room temperature microplate shaker (25 °C, 1200 rpm) to incubate for an additional 20-24 hours. The following day, the plates were centrifuged (4,198 g, 8 min, 25 °C) to collect the cell pellets. The resulting plates containing the cell pellets were covered with an aluminum seal and stored at -80 °C until purification.

Protein purification for screening

The frozen cell pellets were thawed and lysed, and the expressed protein was purified by affinity chromatography with Strep-Tactin® resin following an adapted published procedure.[14] First, three buffers were prepared with Milli-Q water as follows: **W1 buffer** (100 mM Tris-HCl, 150 mM NaCl, 1 mM EDTA, pH 8.0), **W2 buffer** (100 mM Tris-HCl, 150 mM NaCl, pH 8.0), and **E buffer** (40 mM HEPES, 50 mM NaCl, 2.5 mM desthiobiotin, pH 8.0). Additionally, a **lysis buffer** was made from the W2 buffer where it was supplemented with 1 mg/mL lysozyme, 0.5 mg/mL polymyxin B sulfate, 10 µg/mL of DNAse I, and 1 Pierce® protease inhibitor tablet per 50 mL of W2 buffer. All buffers were stirred until the reagents were completely dissolved, and sterile-filtered through a 0.2 µm pore size bottle-filter.

Four plates of cell pellets (i.e., 4 mL of culture for each design) were simultaneously thawed and 600 µL of **lysis buffer** was added to each well in one plate and transferred to the other three plates to combine all of the same-design pelleted cultures in the fourth plate (e.g.,

A1 plate 1 → A1 plate 2 → A1 plate 3 → A1 plate 4). Cells were suspended by repeatedly pipetting the suspension in and out of the tip. The final plate containing the combined cultures in lysis buffer was covered with an aluminum seal and placed in a shaking microplate incubator at 37 °C, 1200 rpm for 60 minutes and then transferred to a 60 °C metal bead bath for 20 minutes. The plate was then centrifuged (4,198 g, x 30 minutes; 14 °C) to pellet the insoluble lysate aggregates before column purification.

Affinity chromatography was performed by adding 200 µL of a 50% Strep-Tactin® resin suspension to each well in a 96-well 25 µm fritted plate such that 1 CV = 100 µL of resin. The resin was equilibrated with 500 µL of **W1** buffer 3 times (3 x 5 CV of W1 buffer) while using a vacuum manifold to pull the liquid through the plate. The bottom spouts of the fritted plate were sealed with parafilm, and 600 µL of the soluble lysate supernatant was gently added to each well in the fritted plate. The fritted plate was covered with an aluminum seal and inverted several times to resuspend the resin. The resuspended resin was allowed to settle at room temperature for 15 minutes before the parafilm and aluminum seal were removed. The protein-loaded resin was washed 2 times with 500 µL of **W1** buffer (2 x 5 CV of W1 buffer) and 1 time with 500 µL of EDTA-free **W2** buffer (1 x 5 CV of W2 buffer) while using a vacuum manifold to pull the liquid through. Finally, the bound protein in each column was eluted into a 2 mL deep 96 well plate by adding 200-300 µL of **E** buffer (1 x 2-3 CV of E buffer) and eluting by centrifugation (1,500 g, 2 min). The eluted protein plate was covered with an aluminum seal and stored at 4 °C until further use. The eluted protein solutions were analyzed by SDS-PAGE to determine the presence of soluble protein. The results of the SDS-PAGE analysis are presented in Fig. S5. We found that 89/96 designs yielded soluble protein band(s). For some proteins, multiple bands were present, possibly because of the presence of multiple species in the obtained eBlocks™ DNA fragments.

The used Strep-Tactin® resin was recovered by first washing each well in the plate with 100-200 µL of E buffer (1 x 1-2 CV of E buffer), 500 µL of W1 buffer (1 x 5 CV of W1 buffer), and 300 µL of W2 buffer (1 x 3 CV of W2 buffer). Finally, the resin was regenerated by adding 3x500 µL of regeneration buffer (purchased as 10x concentrated and diluted in Milli-Q water) (3 x 15 CV of regeneration buffer) until a dark red color was homogeneous through the resin. Note that the resin was resuspended several times via pipetting during the cleaning and regeneration steps to remove aggregates and ensure homogeneity. The regenerated resin wells in the fritted plate were stored under 40-60% v/v ethanol and water to prevent bacterial growth; the plate was covered with an aluminum seal on the top and parafilm on the bottom spouts before being stored at 4 °C until further use.

Screening fluorescence assay (1st round of 96 designs)

The activity screening assay was performed in a 96-well (full-area) fluorescence plate to screen for the hydrolysis of the fluorogenic ester 4MU-PA (Sigma), monitored by a plate-reader using an excitation/emission wavelength of 365 nm and 445 nm, respectively. Briefly, each reaction was run at 30 °C using 6 µL of eluted enzyme containing 200 µM of supplemented zinc sulfate and 54 µL of a substrate solution composed of 10 µM of 4MU-PA in reaction buffer (40 mM HEPES, 50 mM NaCl, pH 8.0) with 5.0% v/v DMSO. Thus, the final reaction in each well contained the eluted enzyme (1/10th the eluted concentration), 20 µM of zinc sulfate, 9 µM of 4MU-PA, and approximately 4.5% v/v DMSO in the reaction buffer.

In a new PCR plate, 48 µL of eluted protein and 2 µL of the 5 mM zinc sulfate solution were combined (final [Zn(II)] = 200 µM). The plate was covered with an aluminum seal, briefly centrifuged (1,000g, 15 s), and incubated at 37 °C for 15 minutes. Afterward, 6 µL of the enzyme-zinc solution from each well was transferred into its corresponding well in a 96-well (full area) fluorescence reaction plate. A stock solution of 100 mM 4MU-PA (Sigma) in DMSO was prepared and used to make a 10 µM working solution of the substrate in reaction buffer with 5.0% DMSO v/v. 54 µL of the substrate working solution was quickly added to every well in the reaction plate using a multichannel pipette. The reaction plate was placed into the plate reader, shaken for 3s at 282 cpm, and fluorescent progress curves monitored for one hour at 30 °C. Note that the enzyme concentration was not standardized in the screening assay, as this was only used to identify hits to characterize in more detail.

Screening fluorescence assay (2nd round of 96 designs)

The activity screening assay was performed in a 96-well (half-area) fluorescence plate to screen for the hydrolysis of the fluorogenic ester 4MU-PA (WuXi), monitored by a plate-reader using an excitation/emission wavelength of 365 nm and 445 nm, respectively. Briefly, each reaction was run at 25 °C using 10 µL of eluted enzyme containing 100 µM of supplemented zinc sulfate and 90 µL of a substrate solution composed of 111 µM of 4MU-PA in reaction buffer (40 mM HEPES, 50 mM NaCl, pH 8.0) with 5.55% v/v DMSO. Thus, the final reaction in each well contained the eluted enzyme (1/10th the eluted concentration), 20 µM of zinc sulfate, 100 µM of 4MU-PA, and approximately 5% v/v DMSO in the reaction buffer.

In a new PCR plate, 49 µL of eluted protein and 1 µL of the 5 mM zinc sulfate solution were combined (final [Zn(II)] = 100 µM). The plate was covered with an aluminum seal, briefly centrifuged (1,000g, 15 s), and incubated at room temperature for 15 minutes. Afterward, 10 µL of the enzyme-zinc solution from each well was transferred into its corresponding well in a 96-well (half area) fluorescence reaction plate. A stock solution of 100 mM 4MU-PA in DMSO was prepared and used to make a 111 µM working solution of the substrate in reaction buffer with 5.55% DMSO v/v. 90 µL of the substrate working solution was quickly added to every well in the reaction plate using a multichannel pipette. The reaction plate was placed into the plate reader, shaken for 3s at 282 cpm, and fluorescent progress curves monitored for one hour at 25 °C. Note that the enzyme concentration was not standardized in the screening assay, as this was only used to identify hits to characterize in more detail.

Isolation and sequence-verification of colonies

Identified screening hits and designs of interest were characterized in greater detail and verified through DNA sequencing and electrospray ionization mass spectrometry (ESI-MS; mass spectrometry methods below). Single clones of identified hits were obtained by the following procedure. A stab from the corresponding polyclonal glycerol stock was diluted into 200 µL of sterile water, and 2 µL of this was further diluted into another 200 µL of water. 20 µL of the diluted cell suspension was spread on an LB-agar plate supplemented with kanamycin, and grown at 37 °C overnight. The following day, 1-3 of the colonies were picked by careful swabbing with a pipette tip and transferred to individual PCR tubes containing 100 µL Milli-Q water. Then, 94 µL of each isolated colony in water was used to inoculate a 1 mL LB culture containing 50 µg/mL of kanamycin for overnight growth and the rest of the isolated colony in water was used for DNA sequencing (described below). The overnight LB+kanamycin cultures were incubated at 37 °C and 225 rpm until the following day when they were visibly opaque (~16-20 hours of growth). Glycerol stocks were made for each of the isolated clones that had verified DNA sequences containing the gene of interest. In the case of a single design having 2+ isolated clones that were DNA-sequence verified, one was chosen at random for glycerol stocks and future experiments. Note that all of these steps were performed using sterile technique under flame.

The DNA sequencing of isolated colonies was performed using Sanger sequencing. ColonyPCR with T7 and T7-term primers was performed to amplify the expressing region of the DNA contained in the isolated cells. The PCR reaction mixture comprised 2 µL of the single colony suspended in nuclease-free water, 1 µL of 10 µM T7 and T7 terminal primers designed to flank the target sequence, 12.5 µL of GoTaq Green 2X Master Mix (which includes Taq DNA polymerase and the accompanying buffer), and 8.5 µL of additional nuclease-free water. The thermal cycling conditions were as follows: an initial lysis and denaturation step at 95°C for 3 minutes, followed by 31 cycles of denaturation (95 °C, 30 s), annealing (58 °C, 30 s), and extension (72 °C, 1 min), culminating in a final extension step at 72 °C for 5 minutes. The success of the PCR reactions was verified by DNA gel electrophoresis using 1.2% agar/TAE gel with SybrSafe dye. Subsequently, 10 µL of unpurified PCR products were sent for sequencing at Genewiz/Azenta, and the results were compared against the design sequences.

Large-scale protein expression with IPTG

DNA sequence-verified, single colony glycerol stocks were used to inoculate 5 mL overnight cultures in LB media with 50 µg/mL of kanamycin that were incubated at 37 °C and 225 rpm. After 16-20 hours, the 5 mL cultures were added to 50 mL of TBII+kanamycin media in 250 mL baffled Erlenmeyer culture flasks, and incubated for 90 minutes at 37 °C and 225 rpm. After the time elapsed, 55 µL of 1 M IPTG solution was added to each of the 55 mL cultures to induce expression. The flasks were incubated at 37 °C and 225 rpm for an additional 2 hours before the temperature was lowered to 25 °C where the flasks were incubated for 20-24 hours to promote sufficient expression. The following day, the cultures were transferred into 50 mL falcon tubes, and cell pellets were collected by centrifugation (4,198g, 8 min, 25 °C). The cell pellets were stored at -80 °C until purification. Note that all steps involving inoculation, media transfers, and IPTG additions were performed with sterile technique under flame.

Protein purification for characterization

The frozen cell pellets were thawed, lysed, and purified by (1) affinity chromatography with Strep-Tactin® resin and (2) size-exclusion chromatography. First, three buffers were prepared with Milli-Q water as follows: **W1 buffer** (100 mM Tris-HCl, 150 mM NaCl, 1 mM EDTA, pH 8.0), **W2 buffer** (100 mM Tris-HCl, 150 mM NaCl, pH 8.0), and **E buffer** (40 mM HEPES, 50 mM NaCl, 2.5 mM desthiobiotin, pH 8.0). Additionally, a **lysis buffer** was made from the W2 buffer where it was supplemented with 100 µg/mL lysozyme, 10 µg/mL of DNAse I, and 1 Pierce protease inhibitor tablet per 50 mL of W2 buffer.

The thawed cell pellets were resuspended in 20 mL of lysis buffer and mechanically lysed by ultrasonication (IBA Lifesciences) at 80% amplitude with 10s on/off pulse cycles for 5 minutes of total “on” time. Note that this was all performed in an ice bath to prevent significant heating of the lysate. The lysates were then centrifuged (4,198 g, 30 min, 4 °C) to collect the supernatant.

Plastic 25 mL gravity flow-through chromatography columns were prepared with 6-10 mL of new Strep-Tactin® resin in a 50% suspension such that 1 CV = 3-5 mL of resin. The columns were equilibrated 2 times with 2.5 CV of W1 buffer followed by 1 addition of 1 CV of W2 buffer. Note that this step and the following steps for affinity chromatography were carried out in a 4 °C room using only gravity flow-through. Once the columns were equilibrated, the lysate supernatant was gently added over the columns and allowed to completely flow through. The columns were then washed 3 times with 1 CV of W1 buffer, 2 times with 1 CV of W2 buffer, and 1 time with 0.5 CV of E buffer. Next, the bound protein in each column was eluted into a falcon tube with 5 additions of 0.5 CV of E buffer. The collected elutants were set aside for further purification by size-exclusion chromatography and the columns were washed 1 time with 2 CV of E buffer before being regenerated by 3 additions of 5 CV of regeneration buffer (obtained as 10x concentrated and diluted in Milli-Q water). Note that the resin was resuspended several times during regeneration so that the columns had a homogenous bright red color after regeneration; the regenerated columns were overlaid with 40-60% v/v ethanol and water to prevent bacterial growth, capped at both ends, and stored at 4 °C for future use.

The collected protein eluants were spin-concentrated to a final volume of 1.0~1.1 mL using a 3 kDa MWCO centrifugal concentrator. The concentrated elutants were then purified by size-exclusion chromatography on an ÄKTA pure™ chromatography system using a Superdex 75 Increase 10/300 GL column with a running buffer of 40 mM HEPES, 50 mM NaCl, pH 8.0. The resulting monomeric fractions of the purified samples were collected and immediately used in downstream experiments. Protein molecular weight was confirmed with electrospray ionization mass spectrometry.

<u>Kinetic analysis</u>

The Michaelis-Menten kinetics of each purified design of interest were determined by incubating the protein samples with zinc sulfate and the fluorogenic substrate 4MU-PA (WuXi) while monitoring the fluorescent signal of the product formation at 25 °C with an excitation/emission wavelength 365 nm and 445 nm, respectively. Before the kinetics, the purified protein samples were diluted to 10x of their desired concentration in the reaction using **reaction buffer** (40 mM HEPES, 50 mM NaCl, pH 8.0). Additionally, zinc sulfate dissolved in Milli-Q water was added to each of these 10x concentrated purified protein stocks to reach a desired [zinc]:[enzyme] ratio. Note that the numeric details for the concentrations of protein and zinc are described in the final paragraph. The samples containing protein and zinc sulfate at 10x the reaction concentration were incubated at 25 °C for 15 minutes. Additionally, stock solutions for the 4MU-PA substrate (WuXi) and 4MU fluorescent product (Sigma) were made by dissolving each of the solids in DMSO.

Two sets of serial dilutions of the 4MU product (1.4 - 45 μM and 0.9 - 30 μM) were made in 95% reaction buffer and 5% DMSO to construct a fluorescence calibration curve, where each point was measured at 25 °C in a 96-well half area plate using 100 μL. Every point was measured in triplicate and averaged over 4 minutes of measurements. The calibration curve was fitted with a linear regression equation to convert fluorescence into the product concentration of 4MU. New calibration curves were made before every series of experiments.

After constructing a standard curve, serial dilutions of the 4MU-PA substrate (1.1 - 144 μM reaction concentration) were made in 94.5% v/v reaction buffer and 5.5% v/v DMSO at concentrations 1.11x the desired concentration in the reaction to account for a 0.9x dilution with the 10x concentrated protein-zinc sample. Kinetic reactions were performed by mixing 10 μL of the 10x concentrated protein-zinc sample with 90 μL of the prepared substrate solution in a 96-well half area plate. The reactions were allowed to shake for 3s at 282 cpm immediately after mixing the enzyme and substrate. Each reaction was monitored for 4 minutes at 25 °C and all substrate concentration points were measured in triplicate. Note that the DMSO was 5.0% v/v in the final reaction mixture. The resulting curves were analyzed by first subtracting the fluorescence of the background reaction at each substrate concentration point for the matching zinc concentration in the reaction buffer. The resulting fluorescence curve was manually set to start at y=0 fluorescence units such that a linear regression (y = mx + b) could be calculated for the linear initial velocity with b=0 (i.e., only the slope, m, needed to be determined). The calculated slopes for the initial velocities at each substrate concentration point were transformed to be expressed in terms of [4MU product]/[enzyme] using the calibration curve and known enzyme concentration. The steady-state parameters $k_{cat}$ and $K_M$ were determined by nonlinear regression fitting all data points to the Michaelis-Menten equation: $v_o/[E]_o = k_{cat}[S]/(K_M + [S])$, where $v_o$ denotes the initial velocity, $[E]_o$ the enzyme concentration, and [S] the substrate concentration. Standard deviations were calculated from the standard error of the fit considering each triplicate value as an individual data point.

To determine the zinc buffer-catalyzed reaction rate, background reactions were carried out without the enzyme at 25 °C in the reaction buffer containing zinc at varying concentrations, and initiated by 4MU-PA substrate (1.1 - 72 μM). The uncatalyzed rate constant was determined from the initial rates of two independent measurements by linearly fitting all data points to the equation $v_0 = k_{uncat}[S]$ using nonlinear regression.

From the 1st design campaign, the kinetics for A1 (ZETA_1) were carried out at $[E]_o$ = 100 nM, 40:1 [Zn(II)]:$[E]_o$ (4 μM zinc sulfate in reaction), and 1.1 - 72 μM 4MU-PA substrate. The kinetics for the active A1 mutants (D67A, H130A, and N17A) were carried out at $[E]_o$ = 100 nM (D67A) or $[E]_o$ = 500 nM (H130A and N17A), 40:1 [Zn(II)]:$[E]_o$, and 1.1 - 72 μM 4MU-PA substrate to be consistent with the measurement of the wild-type A1. The kinetics for C4 were carried out at $[E]_o$ = 3 μM, 10:3 [Zn(II)]:$[E]_o$ (10 μM zinc sulfate in reaction) due to precipitation of this design above this zinc concentration, and 1.1 - 144 μM 4MU-PA substrate. The kinetics for all of the other characterized hits (A8, B9, F7) were carried out at $[E]_o$ = 2 μM, 10:1 $[Zn(II)]:[E]_o$ (20 μM zinc sulfate in reaction), and 1.1 - 144 μM 4MU-PA substrate. Experiments were performed in biological duplicates, with the results being in close agreement. We herein report representative results for each design/variant.

From the 2nd design campaign, the kinetics for ZETA_2 (well position H10) and ZETA_3 (well position H6) were carried out at $[E]_o$ = 100 nM, 10:1 $[Zn(II)]:[E]_o$ (1 μM zinc sulfate in reaction), and 1.1 - 72 μM 4MU-PA substrate. The kinetics for ZETA_4 (well position C5) were carried out at $[E]_o$ = 500 nM, 10:1 $[Zn(II)]:[E]_o$ (5 μM zinc sulfate in reaction), and 1.1 - 72 μM 4MU-PA substrate. Experiments were performed in biological duplicates, with the results being in close agreement. We herein report representative results for each design/variant.

Total turnover number (lower bound estimation)

The lower bound for the total turnover number of A1 (ZETA_1) was estimated using an activity assay monitoring the fluorescent signal generated by the hydrolyzed coumarin product, which was converted into product concentration by a standard curve and taken as a ratio over the enzyme concentration to yield turnover number. The 4MU-PA substrate was loaded at a concentration just below its precipitation limit under the experimental conditions and the enzyme concentration was set as low as possible, while still achieving a high rate above background, to maximize the number of possible turnover events. Purified A1 enzyme was prepared to a stock concentration of 1.0 μM in reaction buffer (40 mM HEPES, 50 mM NaCl, pH 8.0) and supplemented with 40 μM zinc sulfate dissolved in Milli-Q water. Additionally, a stock solution of 4MU product was made in anhydrous DMSO and subsequently used to make serial dilutions of 4MU (4.7 μM - 150 μM) in 95% v/v reaction buffer and 5% v/v DMSO for standard curves. The 4MU-PA substrate was also dissolved in anhydrous DMSO to make a stock solution; this was used to prepare an aqueous solution of 4MU-PA substrate (150 μM reaction concentration) in 94.5% v/v reaction buffer and 5.5% v/v DMSO at concentrations 1.11x the desired concentration in the reaction to account for a 0.9x dilution with the 10x concentrated A1-zinc stock.

After these solutions were made, 100 μL of the 4MU product standards were added to a 96-well half area plate. Reactions were then prepared in the plate by mixing 10 μL of the A1-zinc stock with 90 μL of the prepared substrate solution such that the final A1 concentration was 100 nM, the final zinc sulfate concentration was 4 μM, and the final substrate concentration was 150 μM. Additionally, note that the DMSO was 5.0% v/v in the final reaction mixture. The non-enzymatic background reaction in aqueous reaction buffer with 5.0% v/v DMSO and 4 μM zinc sulfate was also prepared in this plate. After the plate of standards, backgrounds, and reactions on a 100 μL scale was prepared, the plate was immediately covered with a transparent optical film, to prevent evaporation over several hours, and allowed to shake for 5s at 282 cpm. The fluorescent signal of each well in the plate was monitored for 3 hours at 25 °C using an excitation/emission wavelength of 365 nm and 445 nm, respectively. All standards, background reactions, and enzyme-catalyzed reactions were measured in triplicate.

The fluorescence progress curves corresponding to product generation for the background and enzyme-catalyzed reactions were averaged and plotted with the averaged fluorescence signal lines corresponding to the calibration standards of 4MU (Fig. S11A). The calibration curve was constructed by measuring the fluorescence signal of 4MU at each set concentration by averaging the signals during the course of the reaction monitoring experiment, in triplicate (Fig. S11B). A nonlinear calibration curve was observed >75 μM product, which we found to be independent of instrument gain (i.e., the calibration curve always had the same shape), likely due to the common phenomenon of fluorescence self-quenching at high fluorophore concentrations.[15–17] Note that all other experiments only required measuring relatively low fluorescence (i.e., in the linear range) and thus all used linear calibration curves. To overcome this experiment-specific challenge, we used previously reported nonlinear calibration curve fitting methods for interpolation, specifically higher-order polynomial fitting.[18–22] After converting the fluorescence signal into [4MU] using a 4th-order polynomial fit, the y-axis was transformed into units of 4MU product concentration (Fig. S11C). This allowed for the background reaction to be directly subtracted from the enzyme-catalyzed reaction so that any product generated could be attributed to enzymatic catalysis. Turnover number was then calculated based on the ratio of [product]:[enzyme] for the background-subtracted reaction using 100 nM of enzyme to produce the final plot (Fig. 3E).

It should be noted that the conclusion from this experiment (i.e., A1 achieving >1000 turnovers) is not affected by disregarding the interpolation using the nonlinear fit and exclusively considering the discrete fluorescence-[4MU] relationship of the calibration samples (Fig. 11A). By the end of the A1-catalyzed reaction, the fluorescence signal has reached a known value corresponding to 150 μM 4MU, and the background reaction has reached a known value corresponding to 38 μM 4MU. Converting this difference into enzyme turnover number ultimately yields a conservative lower bound estimate of the total turnover number since all of the substrate was hydrolyzed and it was not possible to further enhance the [product]:[enzyme] ratio; we observed limited capacity to raise the initial substrate concentration without precipitation or to reduce the enzyme concentration without substantially lowering the activity-to-background ratio. Thus, A1 (ZETA_1) may catalyze more turnovers.

Preparing the *apo*-enzyme by zinc chelation, and the zinc readdition experiment

The chelation and removal of zinc from the purified A1 (ZETA_1) protein and its mutants were accomplished by dialyzing the sample in a chelation buffer containing 1,10-phenanthroline, a potent zinc chelator, based on a previous protocol.[23] Briefly, 180 mg of 1,10-phenanthroline was dissolved in 500 mL of reaction buffer (40 mM HEPES, 50 mM NaCl, pH 8.0), reaching a final concentration of 2 mM in the **chelation buffer** (40 mM HEPES, 50 mM NaCl, 2 mM 1,10-phenanthroline, pH 8.0). Note that the buffer required heating and vigorous stirring over a stir plate to fully dissolve the 1,10-phenanthroline; the chelation buffer was cooled to 4 °C before dialysis. Wild-type A1 and its mutants were individually loaded into 3.5 kDa MWCO dialysis cassettes and dialyzed against chelation buffer at 4 °C in a beaker with a volume exceeding 400x the sample volume. The beaker had gentle stirring and the dialysis was allowed to take place for 12-24 hours before the dialyzed chelation buffer was exchanged for fresh chelation buffer where the dialysis was allowed to continue for an additional 12-24 hours. After this point, the samples were completely removed of zinc. Part of the wild-type *apo*-A1 sample was directly used in the activity assay and zinc readdition experiment, while the *apo*-mutants and leftover wild-type *apo*-A1 sample were redialyzed in reaction buffer to remove the 1,10-phenanthroline for accurate zinc binding affinity characterization (described in a separate methods section below).

The concentration of the wild-type *apo*-A1 sample in chelation buffer was measured in triplicate (70 μM) and diluted in chelation buffer to a stock concentration of 1.0 μM before the activity assay, which was performed at 100 nM *apo*-A1 concentration (containing ~200 μM 1,10-phenanthroline); wild-type A1 was also prepared at a stock concentration of 1.0 μM and supplemented with 40 μM of zinc sulfate dissolved in Milli-Q water as a control. Additionally, a stock solution of 4MU-PA (WuXi) was prepared in anhydrous DMSO and subsequently used to make an aqueous solution of 4MU-PA substrate (1.8 μM reaction concentration) in 94.5% v/v reaction buffer and 5.5% v/v DMSO at concentrations 1.11x the desired concentration in the reaction to account for a 0.9x dilution with the 10x concentrated *apo*-A1 stock. Kinetic reactions were performed by mixing 10 μL of the 10x concentrated *apo*-A1 stock sample with 90 μL of the prepared substrate solution in a 96-well half area plate where the fluorescent signal of the generated product was monitored at an excitation/emission wavelength of 365 nm and 445 nm, respectively. The reactions were shaken for 3s at 282 cpm immediately after mixing the enzyme and substrate. Each reaction was monitored for 30 minutes at 25 °C and all substrate concentration points were measured in triplicate. Note that the DMSO was 5.0% v/v in the final reaction mixture. The background reactions without added enzyme were also recorded for 4MU-PA hydrolysis in the presence of 4 μM zinc sulfate and without zinc sulfate to match the conditions of the wild-type A1 reaction and *apo*-A1 reaction, respectively.

After the reactions proceeded for 30 minutes, the monitoring was briefly stopped for 2 minutes and 1 μL of 10 mM zinc sulfate dissolved in Milli-Q water was added to the *apo*-A1 and zinc-absent background reactions, achieving a final zinc sulfate reaction concentration of approximately 100 μM. The reactions were again shaken for 3s at 282 cpm before they were monitored for an additional 30 minutes at 25 °C. Note that a large excess of zinc sulfate was added to the *apo*-A1 reaction compared to the wild-type A1 reaction because the *apo*-A1 reaction still had a considerable amount of 1,10-phenanthroline, approximately 200 μM after the 10-fold dilution in the reaction mixture. Thus, it was necessary to add enough zinc to overcome the 1,10-phenanthroline binding, which typically occurs at a 2:1 and 3:1 stoichiometric ratio since zinc has 6 possible coordination sites, so that there was enough zinc to bind in the *apo*-A1 enzymes.[24] However, it should be noted that there is a possibility that 1,10-phenanthroline was binding zinc in the active site and inhibiting the *holo*-A1 (i.e., zinc-bound) structure such that the readdition of excess free zinc simply out-competed the *holo*-A1 zinc for 1,10-phenanthroline binding; therefore, this would stop the inhibition of the *holo*-A1, as this phenomenon has been reported for some native metalloenzymes.[24–26] Nonetheless, the outcome of demonstrating the zinc dependence of A1 (ZETA_1) remains the same.

<u>Zinc binding affinity of A1 (ZETA_1) and its mutants</u>

The Zn(II) ion binding of the design A1 (ZETA_1) and its mutants was evaluated using a competition binding assay with the ratiometric dye Mag-Fura-2 as reporter and competitor.[27] The significant change in absorbance ratio at 323/342 nm (absorbance maximum of metal-bound state to isosbestic point) was used for this analysis. First, the dissociation constant of the dye ($K_D$(Mag-Fura-2, Zn(II)) = 12 ± 2 nM) was determined in competition with nitrilotriacetic acid (NTA); it was measured that $K_D$(NTA, Zn(II)) = 830 pM under the experimental conditions of $I$ = 0.05 M, pH 8.0, and 25 °C. Afterward, the dissociation constants for the proteins were determined in competition with Mag-Fura-2. For each measurement, the proteins, Mag-Fura-2, and/or NTA were each utilized at a 10 µM concentration in a 10 µL buffered solution with varying Zn(II) ion concentrations. The samples were incubated for 3 days in the dark at 25 °C to ensure binding equilibrium. Duplicate measurements were performed for each experiment.

Circular dichroism (CD) spectroscopy

To perform CD spectroscopy, it was necessary to exchange from the HEPES-based buffer to a low-salt, Tris-based buffer for accurate measurement at low wavelengths; otherwise, the machine voltage would jump too high and noise the measurements at these wavelengths. To exchange buffers, the enzymes were dialyzed overnight at 4 °C in 25 mM Tris, 25 mM NaCl, pH 8.0 aqueous buffer.

CD spectra were recorded for the purified A1 (ZETA_1) enzyme (60 µM) in aqueous buffer (25 mM Tris, 25 mM NaCl, pH 8.0) using a JASCO J-1500 Circular Dichroism Spectrophotometer and a 1 mm pathlength cuvette. First, spectra were captured for the sample 25 °C from 190 nm to 260 nm with a bandwidth of 0.1 nm. The sample was then heated to 95 °C in 0.1 °C steps with 5 s averaging time at a rate of 1 °C min-1. Spectra were then captured for the sample at 95 °C using the same parameters as before. Finally, the sample was allowed to cool to 25 °C over about two hours before additional spectra were captured with the same parameters.

SDS-PAGE gel electrophoresis and staining

SDS-PAGE gel electrophoresis was performed with 15- or 26-well precast BioRad AnykD gels. SDS-PAGE samples were prepared by mixing equal volumes of the desired protein sample and 2x-concentrated Laemmli buffer for denaturation. The prepared SDS-PAGE samples were then loaded into the gel in 6 µL aliquots and electrophoresis was performed in SDS-PAGE buffer (25 mM Tris, 192 mM Glycine, and 0.1% (w/v) sodium dodecyl sulfate) at 180 V for 28 minutes until the lowest bands traveled sufficiently through the gel. Note that ladders (Dual Xtra) were also loaded into the gels for reference. After electrophoresis, the gels were removed from their cassette, rinsed with water, and stained for 9 minutes using the eStain L1C protein staining kit. The gels were again rinsed in water and immediately imaged.

Electrospray ionization (ESI) mass spectrometry

MS data for the identified hits and A1 (ZETA_1) mutants were acquired on an Agilent 1200series LC G6230B TOF LC-MS with an AdvanceBio RP-Desalting column (A: H2O with 0.1% Formic Acid, B: Acetonitrile with 0.1% Formic Acid). Final protein concentrations were adjusted to 1-2 mg/mL in 40 mM HEPES, 50 mM NaCl, pH 8.0. Subsequent data deconvolution was performed in Bioconfirm using a total entropy algorithm. All data are presented in Supplementary Table 2.

Statistics

All bar graphs presented in this manuscript show the bar height as the mean and the error bars as 1 standard deviation. All box plots presented in this manuscript show the box extending from the first quartile (Q1) to the third quartile (Q3) of the data, with a line at the median (Q2); the whiskers extend from the box to the farthest data point lying within 1.5x the interquartile range (IQR) from the box. All individual points plotted on curves represent the mean value with error bars representing 1 standard deviation. All curves derived from fluorescent data with respect to time display the mean value as the central line for a given time with the shadow representing 1 standard deviation in either direction.

#### 4. CODE AVAILABILITY

A manuscript detailing the training and *in silico* benchmarking of the deep learning active site scaffolding method introduced in this paper is currently being finalized; we will provide this to the reviewers when ready. Code and neural network weights associated with this manuscript will be made publicly available on GitHub upon publication of this manuscript. PLACER is publicly available on GitHub at the link: https://github.com/baker-laboratory/PLACER.

#### 5. SUPPLEMENTAL FIGURES

Fig. S1. Schematic overview of the computational design of de novo metallohydrolases with the pipeline trajectory of A1 (ZETA_1).

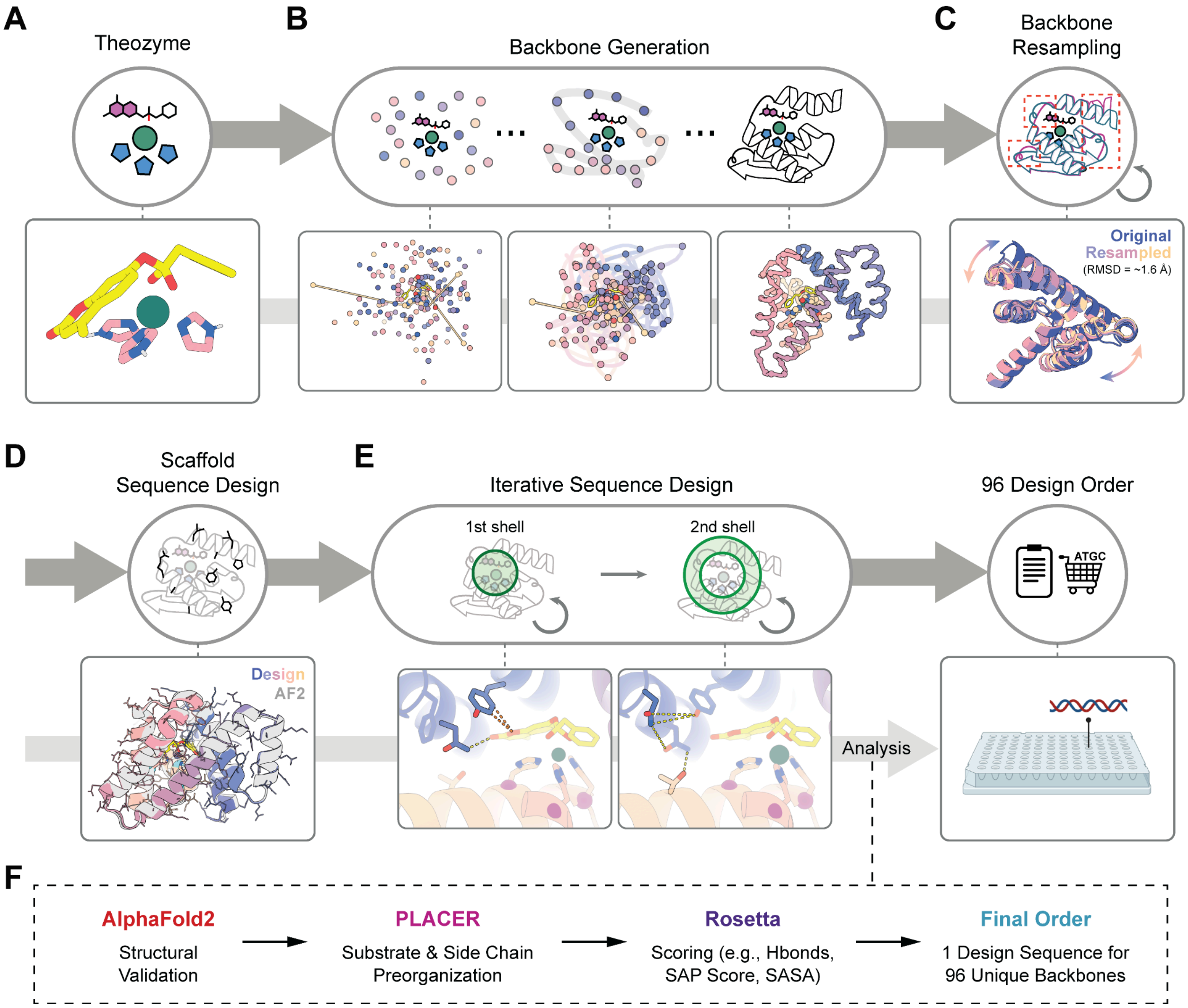

**Fig. S1.** Schematic overview of the computational design of de novo metallohydrolases with the pipeline trajectory of A1 (ZETA_1). (**A-F**) Visual illustrations of the key steps in the computational design pipeline with examples showing A1 at each stage in the pipeline.

Fig. S2. Detailed flow chart of the design and experimental pipeline for de novo metallohydrolases.

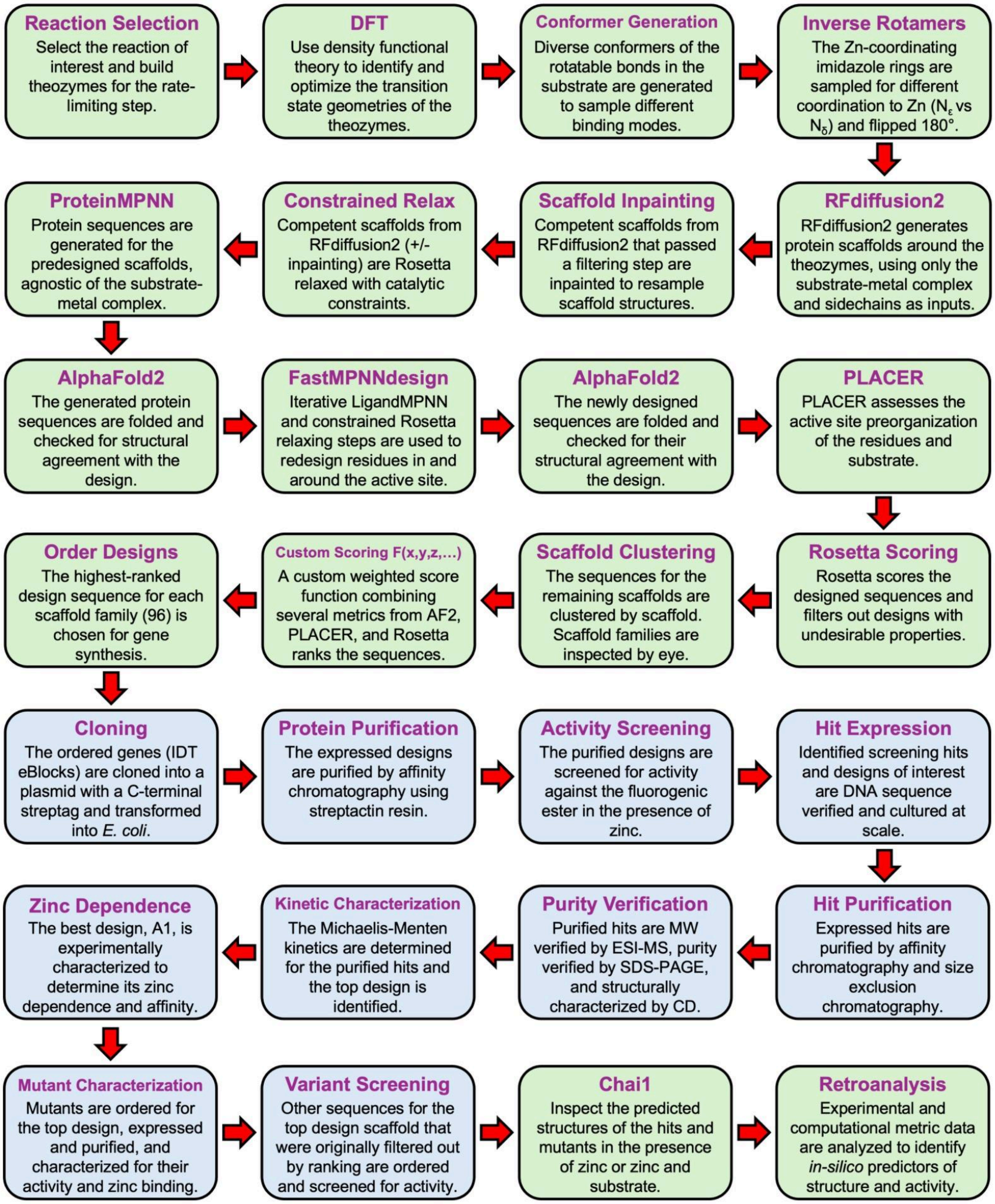

**Fig. S2.** Detailed flow chart of the design and experimental pipeline for the computational design of de novo metallohydrolases. Note that the green color represents steps done *in silico* while the blue color represents experimental steps done in the wet lab.

Fig. S3. Theozyme input generation for RFdiffusion2 and the challenge of enumerative catalytic motif scaffolding.

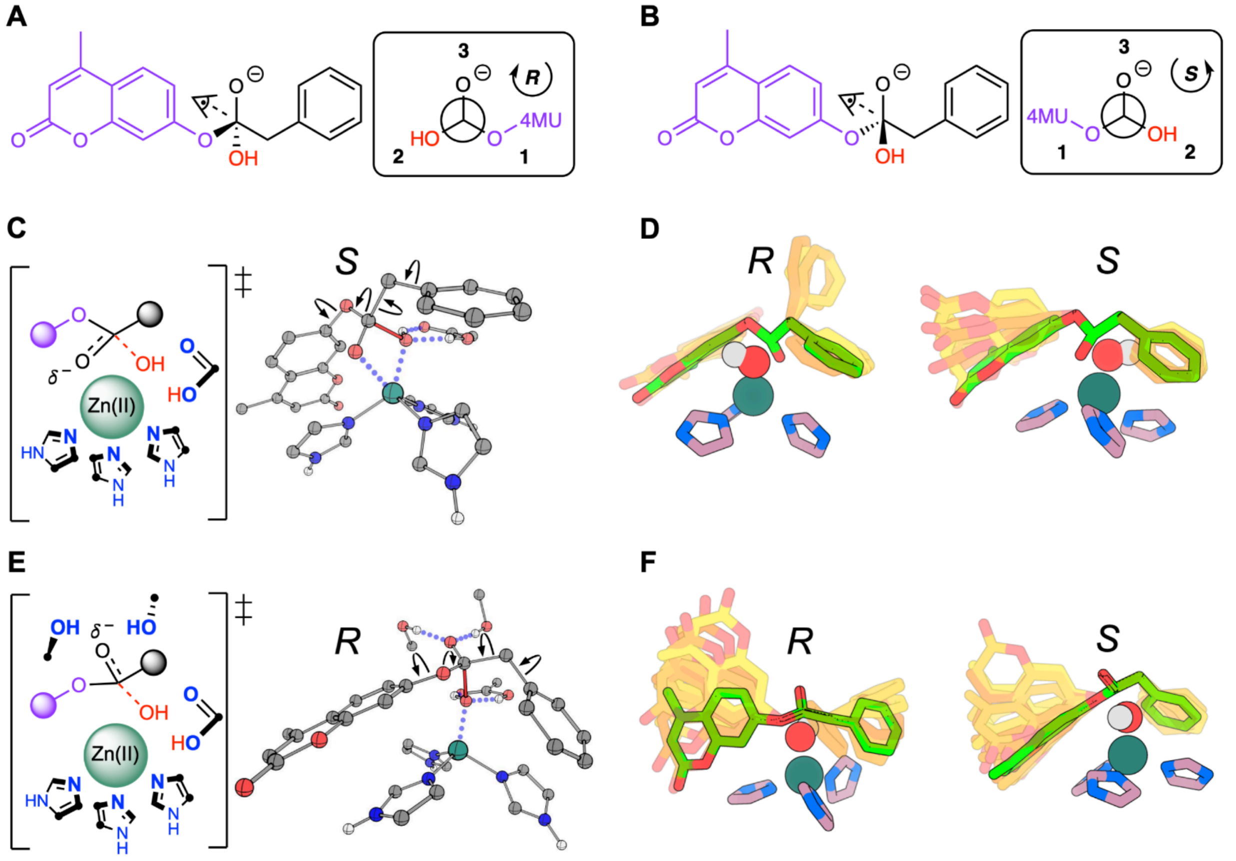

**Fig. S3.** Theozyme inception and transition state optimization with density functional theory followed by substrate conformer generation. (**A,B**) Tetrahedral intermediate enantiomers. (**C,E**) Example theozymes for zinc-hydroxide nucleophilic attack of the 4MU-PA ester modeling (**C**) oxyanion stabilization via zinc or (**E**) oxyanion stabilization via protein hydrogen bonds. 2D representation (left) and 3D DFT model (right). Arrows on the 3D model represent sampled conformational flexibility. (**D,F**) Substrate conformers (yellow spectrum) from each DFT model (green). Note that the general base, and the oxyanion hole in the case of the (E,F), were not modeled during the scaffold generation with RFdiffusion2; these were left to be found during sequence design due to the flexibility of residues that can accomplish those tasks.

### Fig. S4. Catalytic constraints and enumerative catalytic motif scaffolding around theozymes for RFdiffusion2 compared to RFdiffusion.

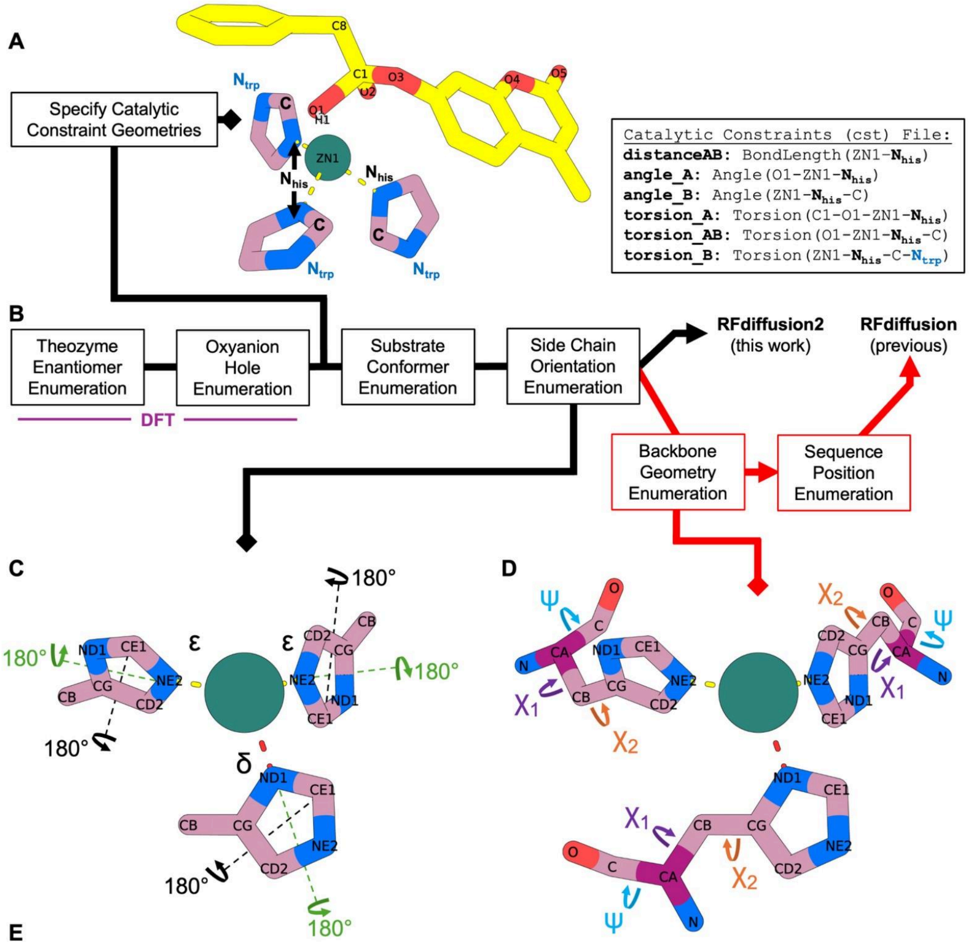

E

| |
|---|
| **4** theozymes x **10** conformers x **2³** coordination permutations x **2³** ring flips<br>= **2560 RFdiffusion2 Inputs**<br><br>**OR**<br>(assuming 100 AA protein with only 10 rotamers each for χ1, χ2, and ψ)<br><br>**2560** x **10³**(χ1) x **10³**(χ2) x **10³**(ψ) x **100!/(100-3)!** sequence positions<br>= **>10¹⁸ RFdiffusion Inputs** |

**Fig. S4.** Catalytic constraints and enumerative catalytic motif scaffolding around theozymes for RFdiffusion2 compared to RFdiffusion. (**A**) Example theozyme from DFT with relevant ligand atoms labeled and Rosetta constraint file labeling for the histidine sidechains. Catalytic constraint files specifying the coordination geometry of the histidines were created at this step for each of the 4 theozymes from DFT. Note that $N_{his}$ is a general Rosetta label that accommodates epsilon and delta coordination geometries with zinc. (**B**) Theozyme sampling enumeration pipeline with diverging paths for RFdiffusion2 and RFdiffusion. (**C**) Sidechain enumeration sampling different combinations of zinc coordination and imidazole orientations. Note that there are only 64 permutations of coordination and $C_\beta$ positioning. (**D**) Backbone enumeration for RFdiffusion. This is required in addition to the sampling in (**C**). (**E**) Calculation of the total number of possible RFdiffusion2 and RFdiffusion inputs. It should be noted that not all combinations of rotamers and sequence positions will be viable, somewhat reducing the total number, but to nowhere near the tiny enumeration space needed for RFdiffusion2.

Fig. S5. Structural panel of the 96 ordered designs spanning 96 unique backbones.

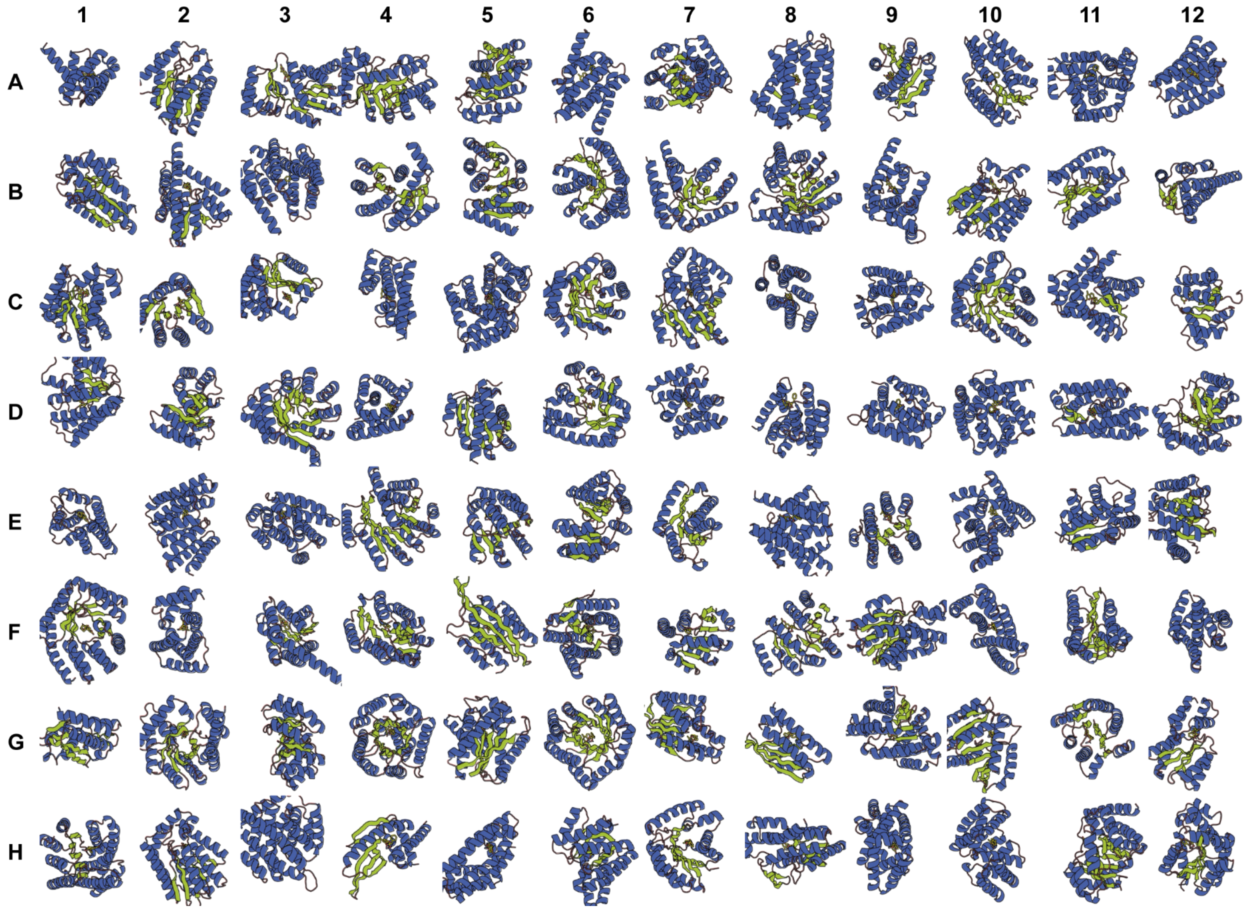

**Fig. S5.** Structural panel of the 96 ordered designs spanning 96 unique backbones. Each structure corresponds to the well it was ordered in based on its *in silico* ranking (A1 = best ranking; H12 = worst ranking). The secondary structures are colored (green = beta-sheets; blue = alpha-helices). Structures range in length from 140 - 275 amino acids and have a range of diverse folds.

Fig. S6. SDS-PAGE gel results for 96 designs expressed and purified in parallel.

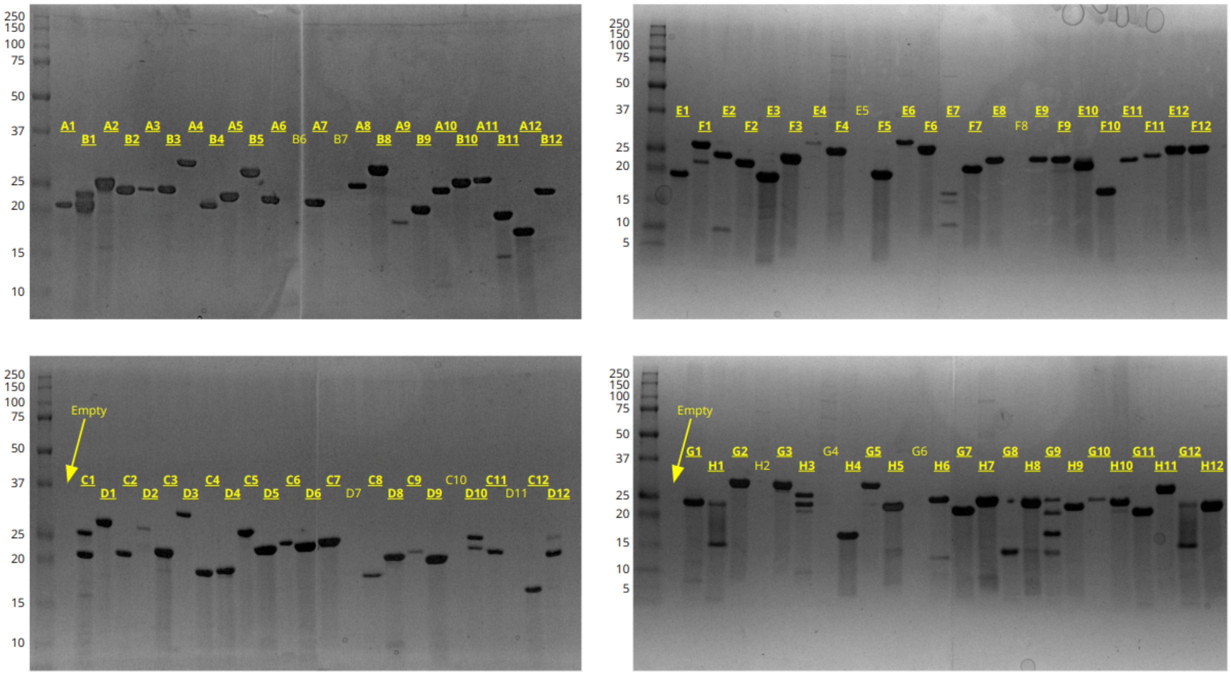

**Fig. S6.** SDS-PAGE gel results for 96 designs expressed in *E. coli* and purified by using the Strep-Tag affinity chromatography. Designs that had any visible sign of a band were counted as an expressed design (bolded and underlined text). Note that the ladder units are in kDa and that the bottom two gels had an empty spacer well. 86/96 designs show at least one band.

Fig. S7. Initial metallohydrolase activity screen with the purified designs.

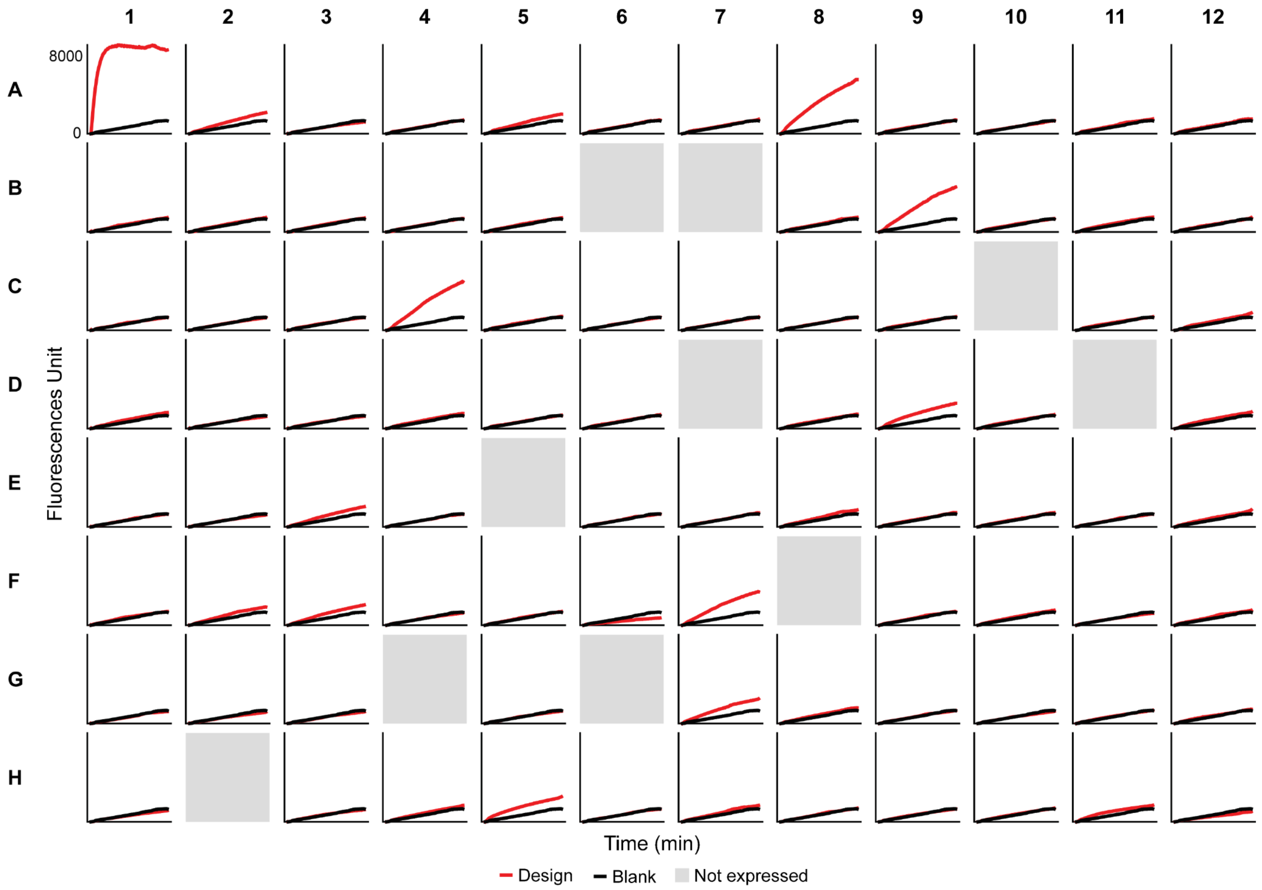

**Fig. S7.** Initial metallohydrolase activity screen with the purified designs. The hydrolysis reactions of 100 μM 4MU-PA were monitored by the fluorescence generated from the product over 1 hour. The red progress curves display the reaction in the presence of a purified enzyme (+200 μM added zinc sulfate) and the black progress curve overlaid on every plot displays the background reaction in the presence of 200 μM zinc sulfate for comparison. Note that the position of each graph corresponds to its design well position (i.e., top left is design A1) and that the grayed-out plots correspond to design wells that did not express as judged by SDS-PAGE (Fig. S5).

Fig. S8. SDS-PAGE gel results and SEC chromatograms of the purified hits.

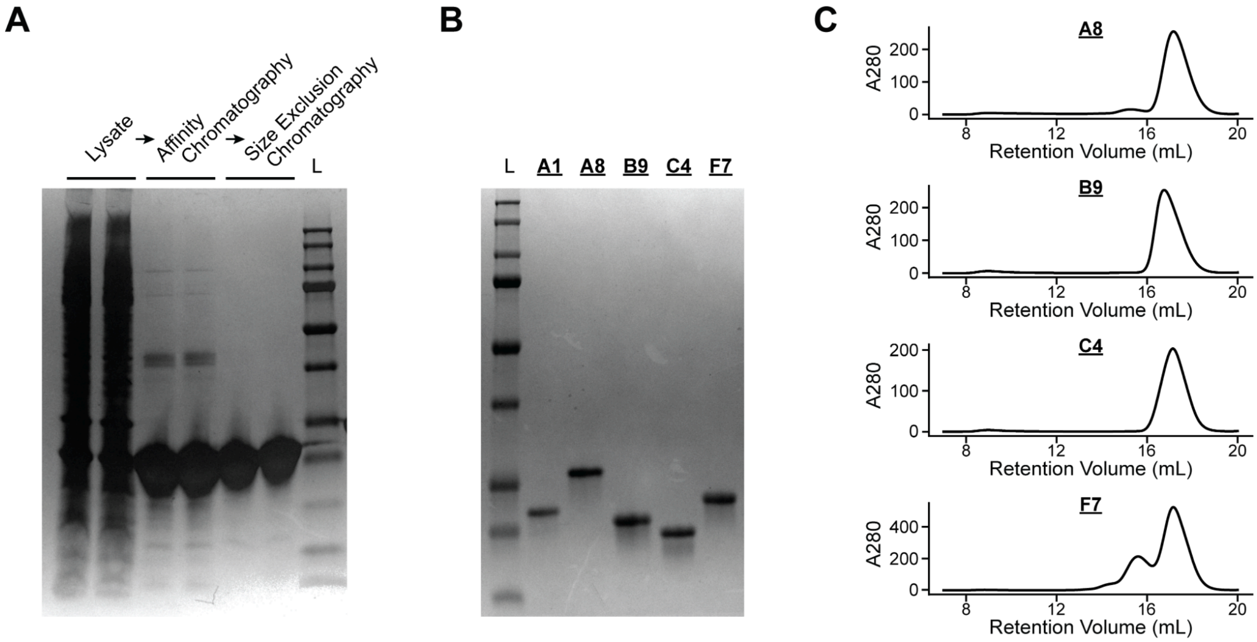

**Fig. S8.** SDS-PAGE gel results and SEC spectra for the purified hits expressed at scale. (**A**) Purposefully-overloaded SDS-PAGE gel for the A1 (ZETA_1) design in lysate (left), after affinity chromatography (middle), and after affinity chromatography and size exclusion chromatography (right) with a ladder (far right). Duplicate A1 samples were loaded at each stage. This gel conveys that even with significant protein loading, which should increase the visibility of any contaminants, we are unable to see contaminants after affinity chromatography and size exclusion chromatography, validating the purification protocol and testifying to the high purity of A1 used in further experiments. (**B**) SDS-PAGE gel for the 5 screening hits purified by affinity chromatography and size exclusion chromatography. Note that no contaminant bands are visible and all of the bands are around their expected mass as judged by the ladder position. (**C**) Size exclusion chromatography A280 traces during elution. Most proteins obtain a clean monomeric peak except F7, which appears to have a slight tendency to oligomerize. Only the fraction corresponding to the highest peak in the monomeric state was used for each design. The size exclusion chromatography A280 trace of A1 can be found in the main text (Fig. 3C).

Fig. S9. Electrospray ionization mass spectrometry for A1 (ZETA_1).

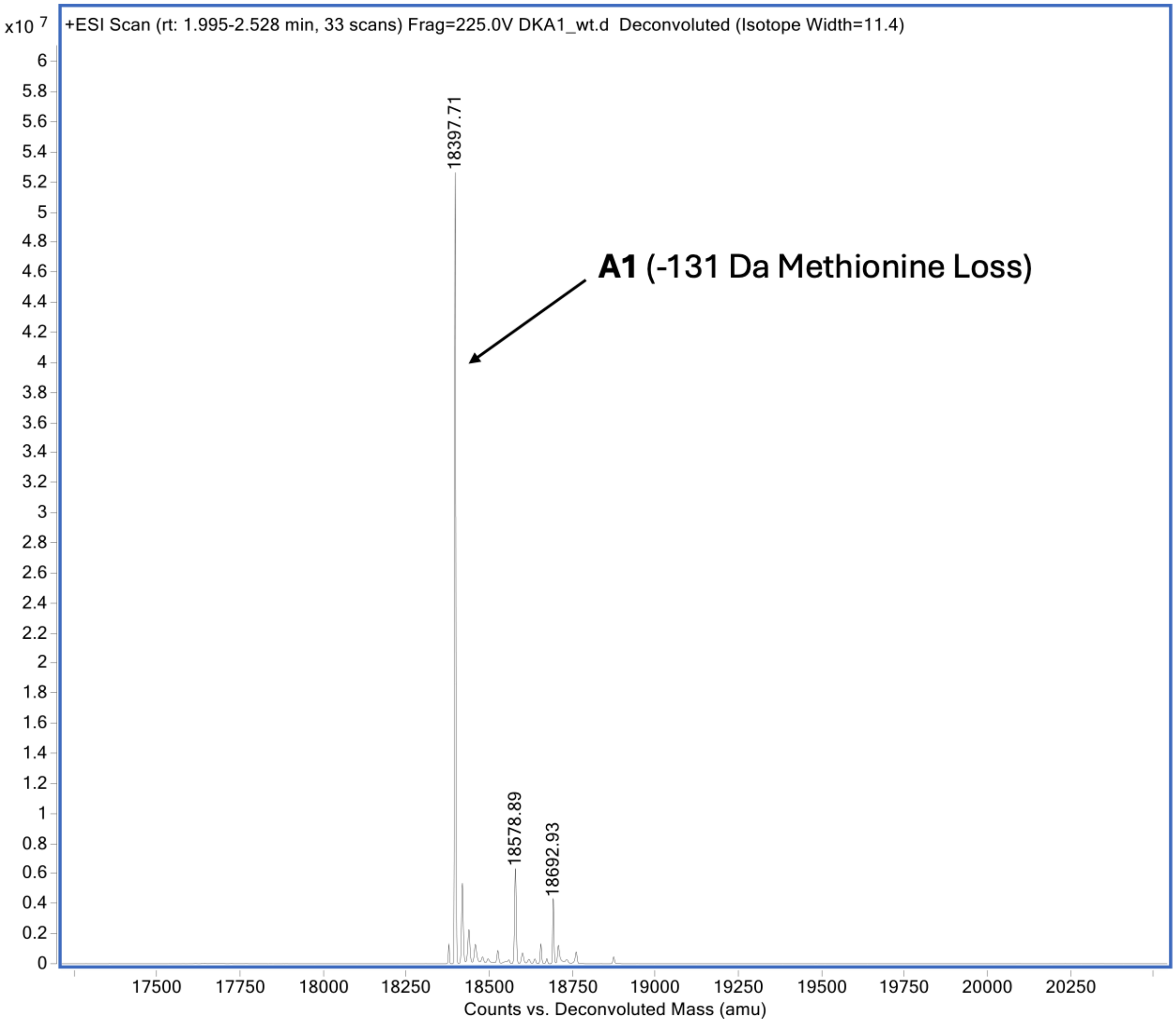

**Fig. S9.** Electrospray ionization mass spectrometry for A1 (ZETA_1). The expected molecular weight is 18528 Da. The highest peak is approximately 18397 Da, -131 Da from A1 corresponding to the loss of N-terminal methionine (-131 Da). All ESI-MS values are given in Table S2.

Fig. S10. Michaelis-Menten kinetic characterization of the 5 design hits.

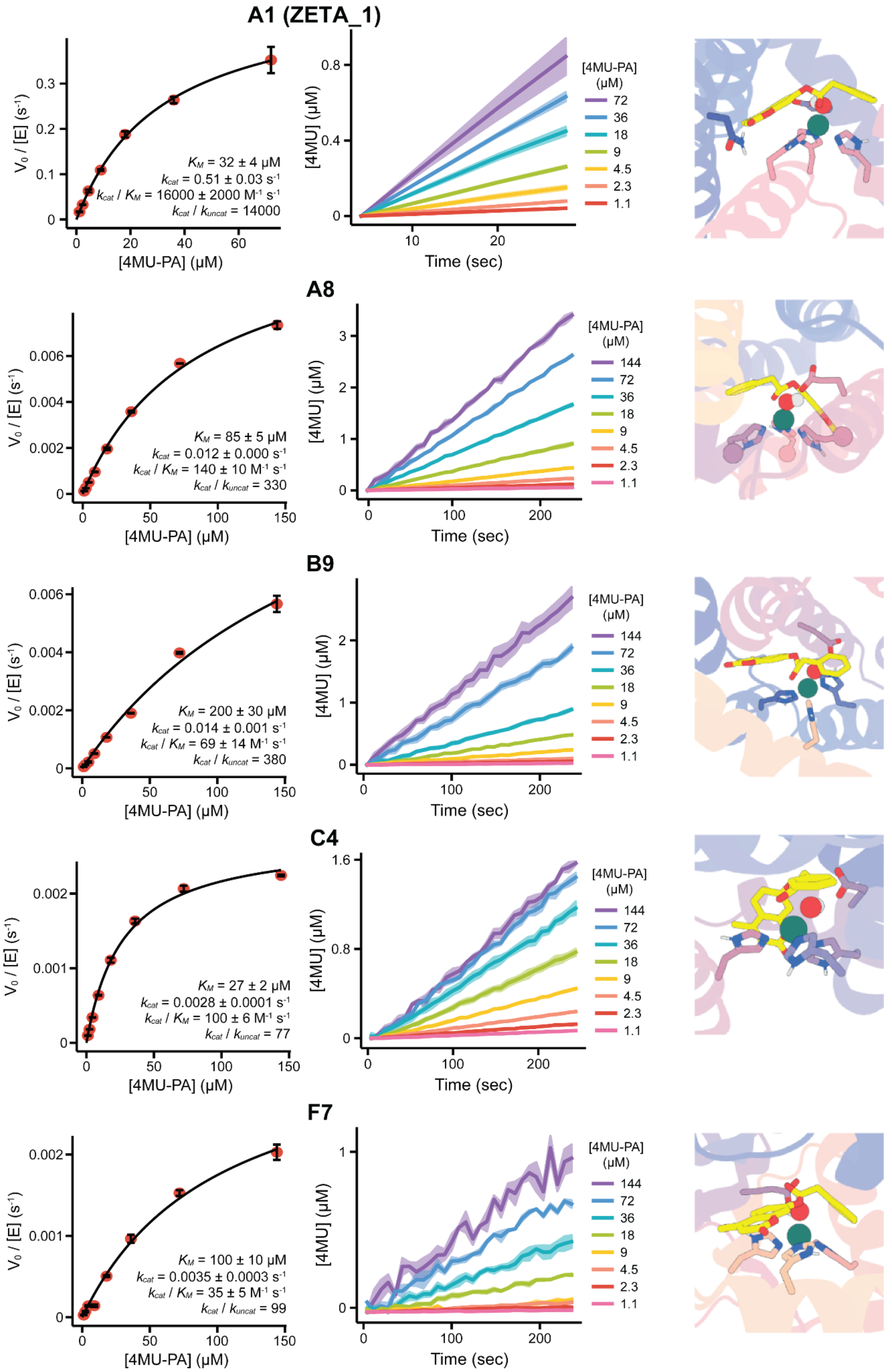

**Fig. S10.** Michaelis-Menten kinetic characterization of the 5 design hits. Michaelis-Menten plots from these data are shown on the left. The progress curves for the measured initial velocities are shown in the middle. Designed active sites with the histidines and general base are displayed on the right.

Fig. S11. Total turnover lower bound experiment for A1 (ZETA_1) and calibration curve.

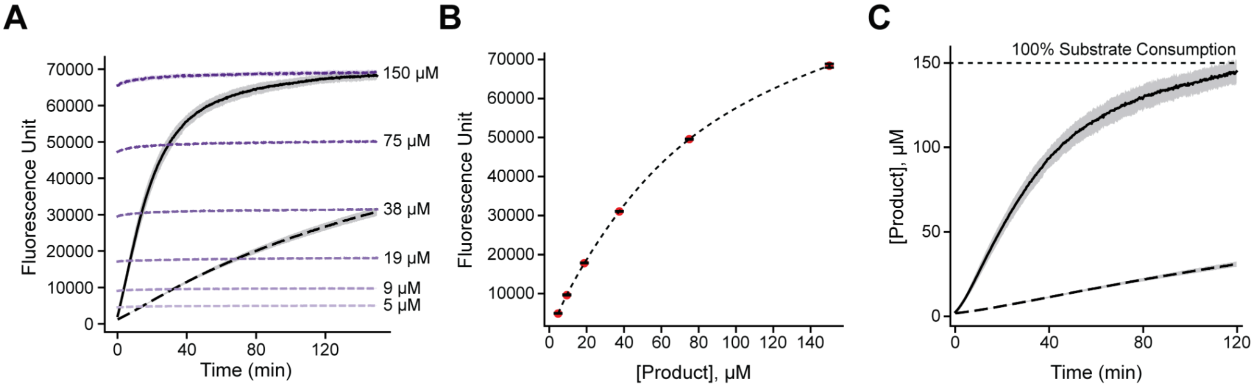

**Fig. S11.** Total turnover lower bound experiment for A1 (ZETA_1) and calibration curve. (**A**) Fluorescent progress curves for the A1-catalyzed reaction (solid black curve), background reaction with zinc (dotted black curve), and 4MU product calibration standards at labeled concentrations (rainbow lines). (**B**) Calibration curve from the 4MU product standards for the turnover experiments with a fitted 4th order polynomial regression curve for interpolation. Note that the calibration curves used for all other experiments were in the linear range and fitted with linear regression; this experiment was an exception because it required measurements of high fluorescence corresponding to highly concentrated 4MU product, which displays nonlinear behavior likely due to fluorescence self-quenching at high concentrations. The interpolation does not affect the final results of this experiment (i.e., A1 catalyzes >1000 turnovers). (**C**) A1-catalyzed reaction and background reaction with 4 μM zinc plotted in terms of product concentration after using the calibration curve to convert the fluorescence signal into product concentration. The A1 enzyme was present at 100 nM and all reactions used 150 μM of 4MU-PA substrate; enzyme turnover number is calculated by the ratio of [product]:[enzyme] after subtracting off the background reaction.

### Fig. S12. In silico metric distributions for the ordered designs.

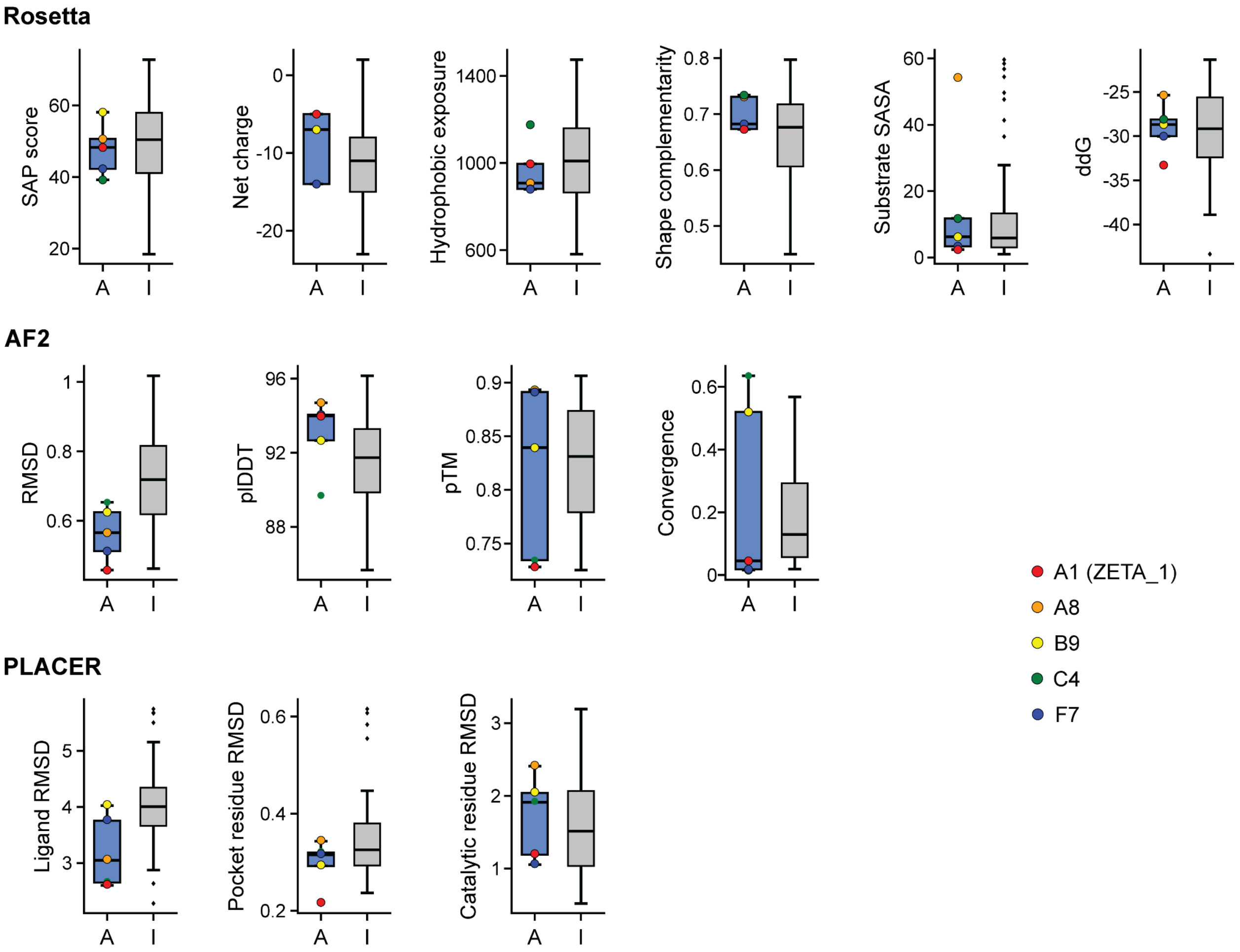

**Fig. S12.** *In silico* metric distributions for the ordered designs. ‘A’ represents the computational metric distribution for the 5 ‘Active’ designs (A1, A8, B9, C4, F7) and ‘I’ represents the computational metric distribution for the other 91 ‘Inactive’ designs, including the 10 designs that did not express. These plots compare the distributions for common and insightful metrics from Rosetta, AlphaFold2, and PLACER between the 5 active and 91 inactive designs. The box plot shows the five-quartile summary and the identified outliers are shown as small black dots for the inactive designs. The data points for the 5 active designs are shown as colored dots in each metric with the corresponding color legend shown on the right.

Fig. S13. Comparison of AlphaFold2 metrics with PLACER metrics for ordered designs.

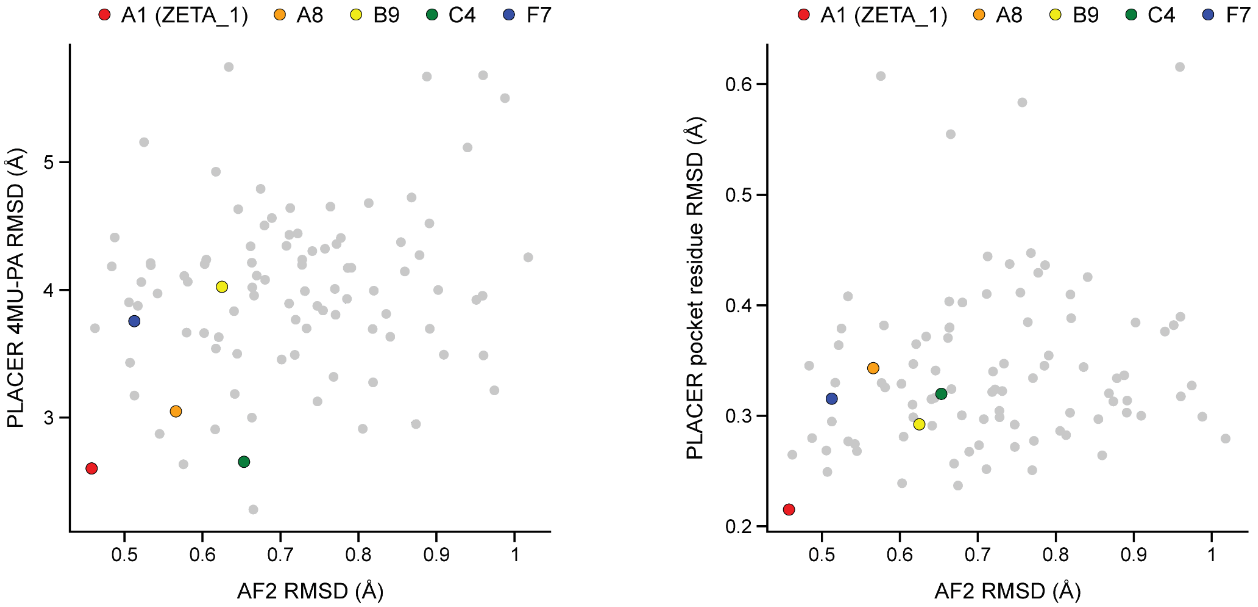

**Fig. S13.** Comparison of AlphaFold2 metrics with PLACER metrics for ordered designs. (**A**) Average substrate RMSD between the design model and the predicted PLACER ensembles (PLACER 4MU-PA RMSD) vs the global $C_\alpha$ RMSD between the design model and the AlphaFold2 prediction (AF2 RMSD). (**B**) Average active site residue RMSD between the design model and the predicted PLACER ensembles (PLACER pocket residue RMSD) vs the global $C_\alpha$ RMSD between the design model and the AlphaFold2 prediction (AF2 RMSD). All units are in Å. Note that the bottom left corner is ideal in both plots, indicating high agreement between the design and predictions from AlphaFold2 and PLACER. Interestingly, A1 (ZETA_1) is the closest to the bottom left in each of these plots, suggesting that high agreement between AlphaFold2 and PLACER may be a useful predictive metric of activity.

Fig. S14. Computational and experimental comparison of A1 (ZETA_1) against other sequences identified for that backbone during design.

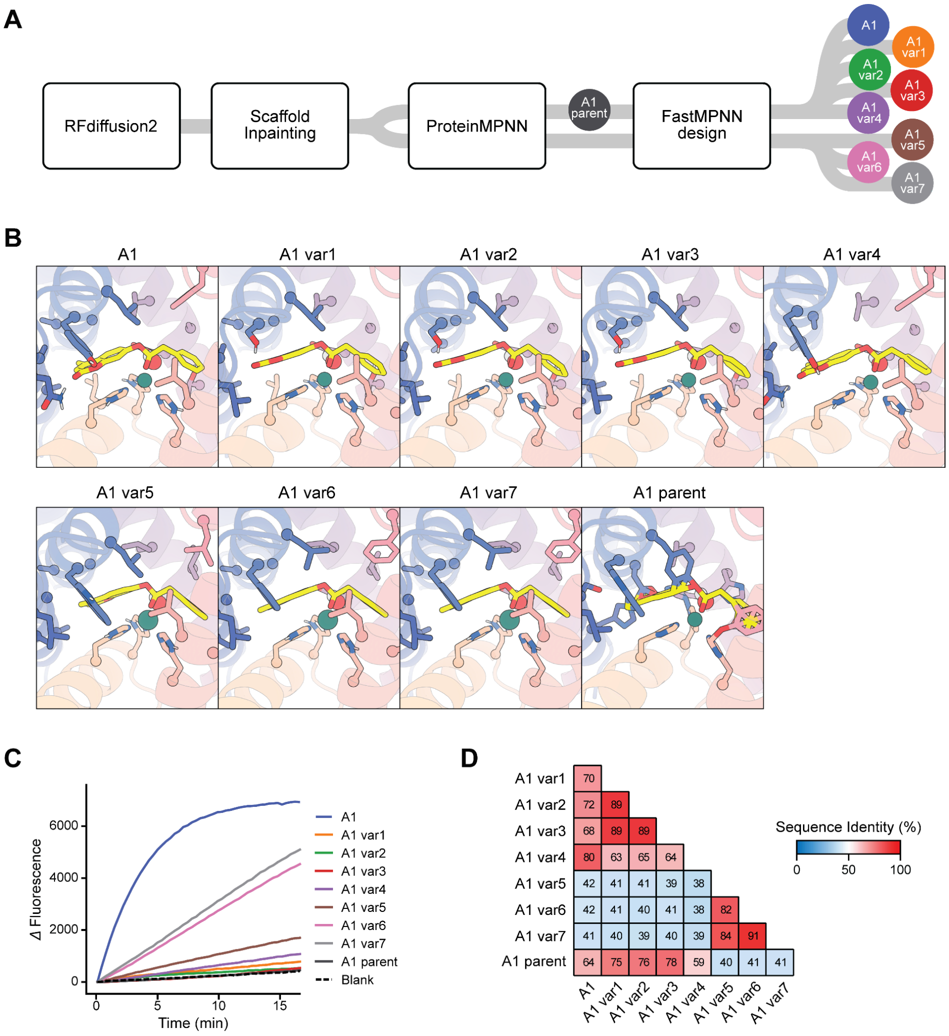

**Fig. S14.** Computational and experimental comparison of A1 (ZETA_1) against other sequences identified for that backbone during design. (**A**) High-level computational pipeline overview visualizing the conception of A1, the parent A1, and 7 variants. Note that all variants came from the same RFdiffusion2 scaffold but variants 5-7 came from a separate resample. (**B**) Comparison of the active sites between A1 and its 7 sequence variants that made it to the final stages of computational filtering. The "A1 parent" is the originally folded sequence from ProteinMPNN, which is agnostic of ligands and metals, before the active site optimization step with iterative LigandMPNN and Rosetta design, hence the clashes. (**C**) Ester hydrolase activity assay comparing A1 with its 7 variants and parent sequence demonstrating that A1 has superior activity. This provides confidence in our *in silico* filtering method, which was able to successfully identify the best sequence for the A1 backbone for the 96-design order with 96 unique backbones (1 sequence / scaffold). (**D**) Heatmap comparing the sequence similarity between A1, the A1 parent, and all of the A1 variants. Shades of blue represent lower sequence identity (<50%) between two designs and shades of red represent higher sequence identity (>50%) between two designs. Variant 4 is the most similar in sequence to A1 but still only shares 80% sequence identity, showing that this pipeline explores a highly diverse sequence space within each scaffold during design.

Fig. S15. Predicted structures of the A1 (ZETA_1) design.

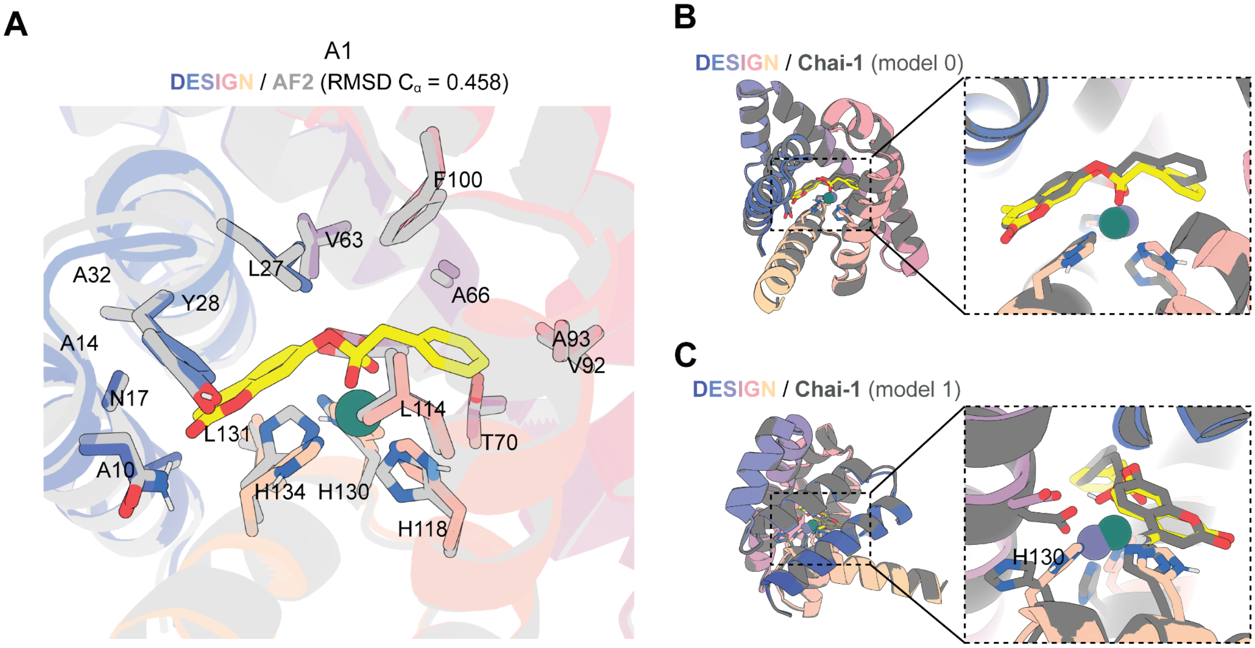

**Fig. S15.** Predicted structures of the A1 (ZETA_1) design. (**A**) Active site of the A1 design model with the pocket residues labeled. The designed structure is colored and the AlphaFold2 *apo*-predicted structure is in gray. The structures are in high agreement, suggesting that A1 has a high degree of preorganization. (**B**) Global and active site comparison of the A1 design model (colored) with model 0 of the Chai-1[28] structure sequence/scaffold) in the presence of zinc and 4MU-PA. Note that the substrate SMILES file was provided to Chai-1, not the tetrahedral intermediate, whereas the A1 design model contains the transition state of the nucleophilic attack of hydroxide to the substrate. The global scaffold fold prediction by Chai-1 is nearly identical to the design model scaffold (RMSD $C_\alpha$ = 0.413). Additionally, the zinc and substrate docking positions by Chai-1 are nearly identical to the design model; model 0 of Chai-1 also predicts the zinc being coordinated by the 3 histidines (H118, H130, H134) in nearly the same orientations as the design model. (**C**) Global and active site comparison of the A1 design model (colored) with model 1 of the Chai-1 structural prediction (gray) in the presence of zinc and 4MU-PA. Note this is shown from the side/back view. Again, the global scaffold fold prediction and substrate docking position by Chai-1 are nearly identical to the design model. Interestingly, Chai-1 predicts a slightly different zinc docking position in model 1 with H130 flipped away from the active site, and D67 is instead coordinating zinc with the other two histidines (H118 and H134). This supports the hypothesis that H130 and D67 are competitively binding zinc, explaining the results observed from the mutant activity and zinc binding experiments. Thus, it is unlikely that D67 is functioning efficiently, if at all, as a general base, providing a rational starting point for potential redesign.

Fig. S16. Structural novelty of the A1 (ZETA_1) design.

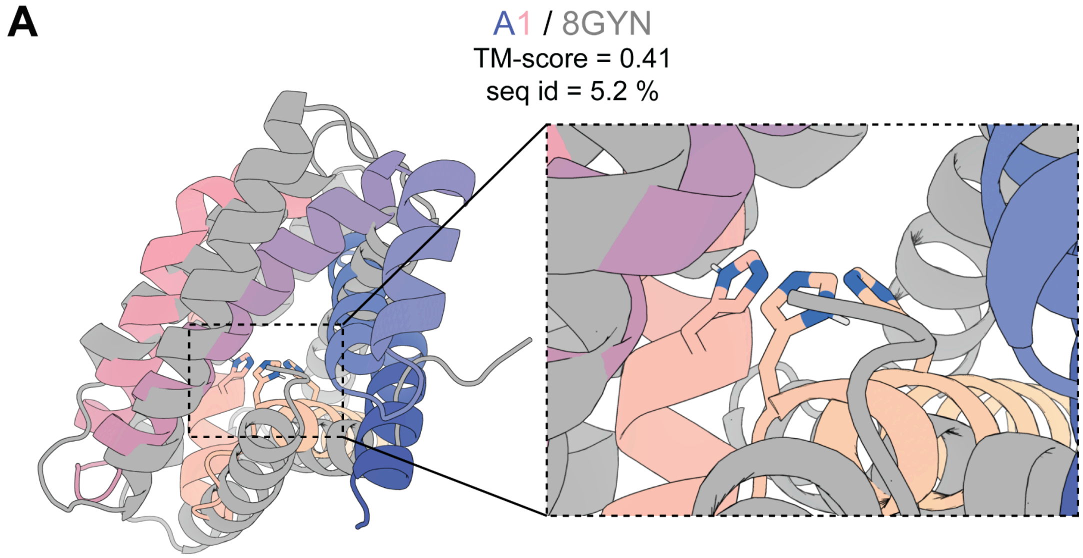

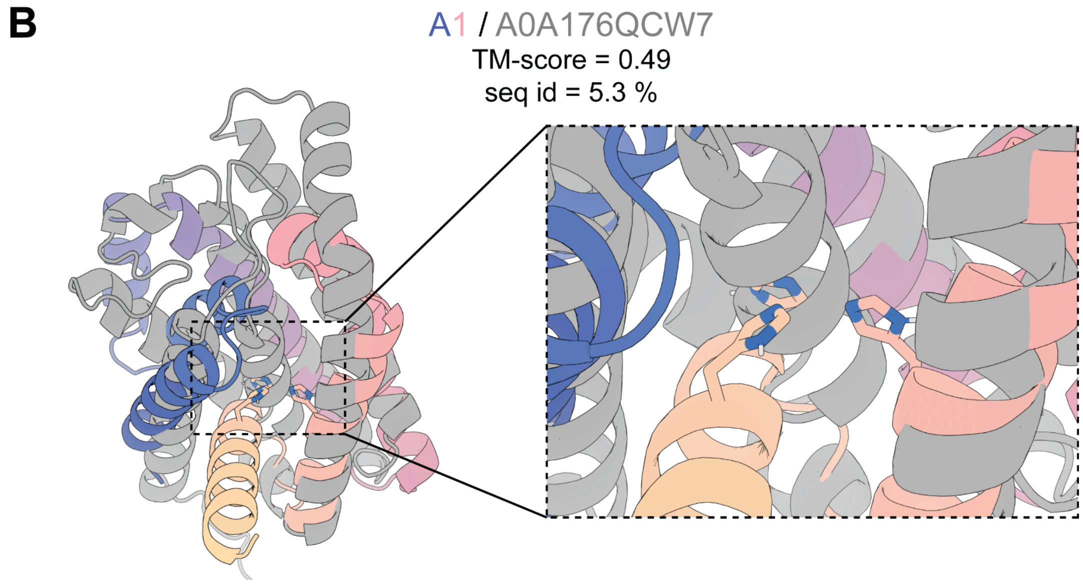

**Fig. S16.** Structural novelty of the A1 (ZETA_1) design. (**A**) Global $C_\alpha$ alignment of A1 with the PDB structure, 8GYM, the most similar structure to A1 in the PDB based on the template modeling (TM) score. Note the alignment, sequence identity, and TM-score are poor, reflecting the novelty of the A1 scaffold. (**B**) Global $C_\alpha$ alignment of A1 with the AlphaFold2 predicted structure, A0A176QCW7, the most similar structure to A1 in the AlphaFold2 database based on TM-score. Note the alignment, sequence identity, and TM-score are poor, further reflecting the novelty of the A1 scaffold.

Fig. S17. SDS-PAGE gel results and SEC spectra for the purified A1 (ZETA_1) knockout mutants expressed at scale.

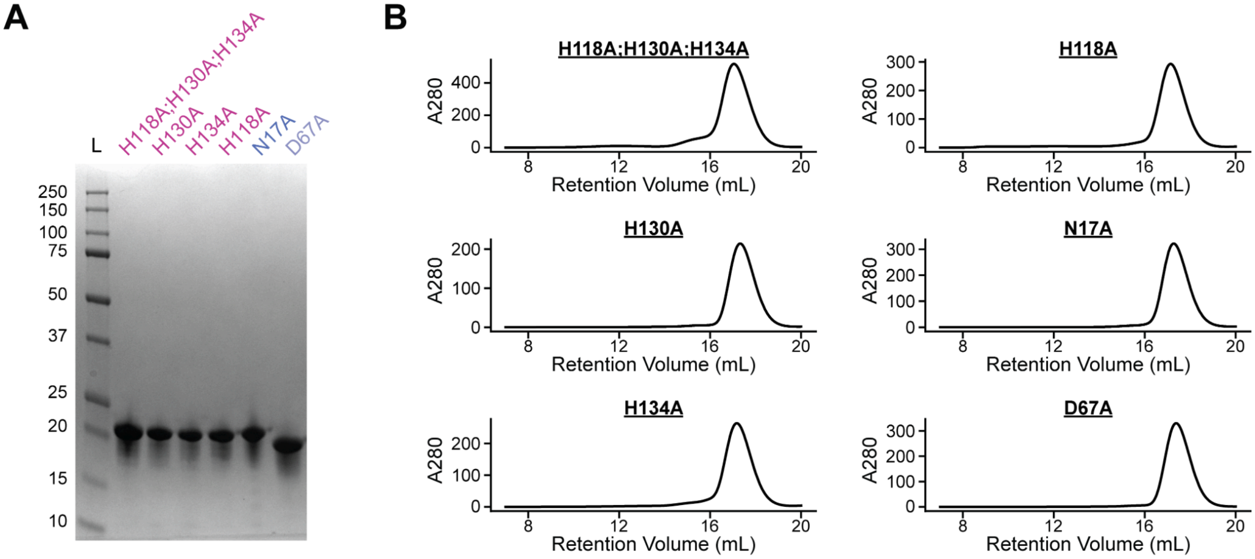

**Fig. S17.** SDS-PAGE gel results and SEC spectra for the purified A1 (ZETA_1) knockout mutants expressed at scale. (**A**) SDS-PAGE gel for the six A1 mutants purified by affinity chromatography and size exclusion chromatography, respectively. Note that no contaminant bands are visible and all of the bands are around their expected mass as judged by the ladder position. (**B**) Size exclusion chromatography A280 traces during elution. All of the mutants demonstrate a clean monomeric peak. Only the fraction corresponding to the highest peak in the monomeric state was used for each design.

Fig. S18. Kinetic characterization and Chai-1 predicted structures of the A1 (ZETA_1) knockout mutants with the proposed role of residue N17.

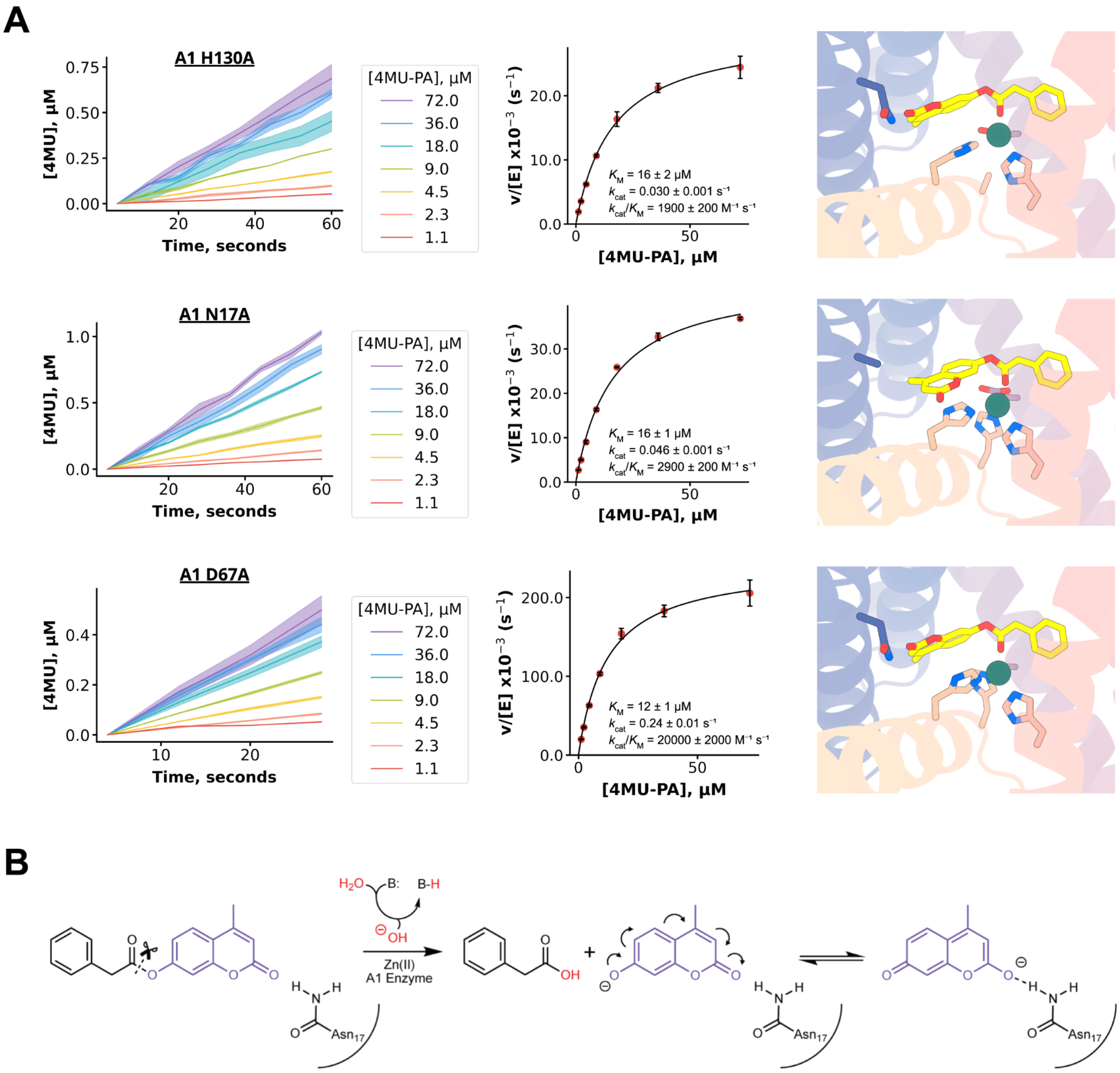

**Fig. S18.** Kinetic characterization and Chai-1[28] predicted structures of the A1 (ZETA_1) knockout mutants with the proposed role of residue N17. (**A**) The progress curves for the measured initial velocities are shown on the left. Michaelis-Menten plots from these data are shown in the middle. The Chai-1 predicted structures and predicted complexes of the protein, zinc, and substrate are shown on the right for each mutant. Chai-1 predicts the zinc and substrate in almost the exact same position and geometry as the A1 design model for each of the active mutants. Additionally, Chai-1 predicts the D67 coordination to zinc in the H130A mutant as hypothesized. Interestingly, the N17A substrate docking position by Chai-1 reveals a slightly shifted coumarin (4MU) conformation, suggesting N17 likely plays a role in positioning the coumarin and possibly a role in stabilizing the coumarin leaving group. (**B**) Proposed stabilization of the coumarin leaving group through resonance and H-bonding to N17.

Fig. S19. Zinc-binding affinity experiments for A1 (ZETA_1) and its knockout mutants.

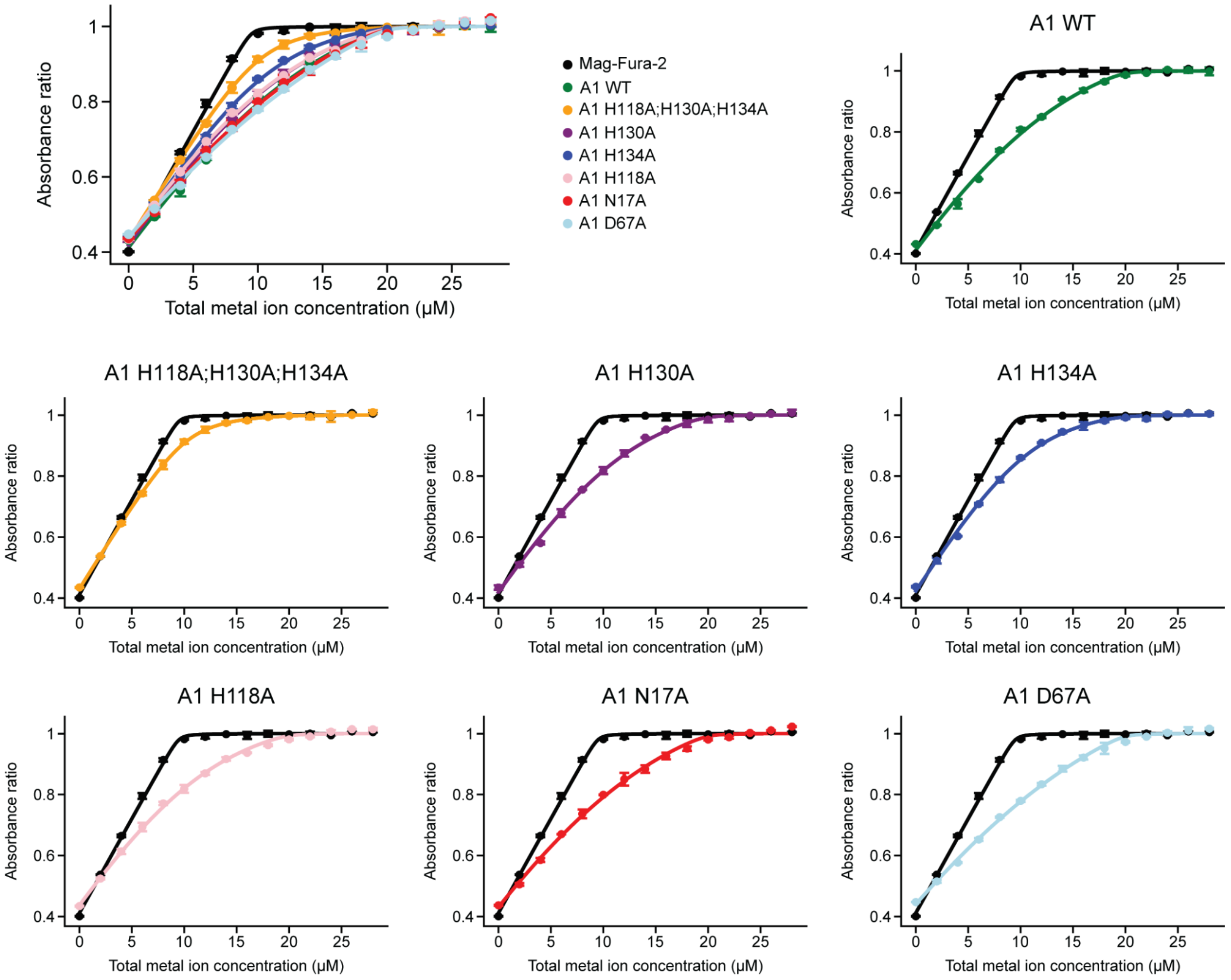

**Fig. S19.** Zinc-binding affinity experiments for A1 (ZETA_1) and its knockout mutants. The plots display the zinc-binding profiles of the wild-type A1 enzyme and its mutants. As the total zinc ion concentration increases, the absorbance ratio plateaus reflecting saturated zinc binding.

Fig. S20. PLACER metrics of ordered designs from each design campaign.

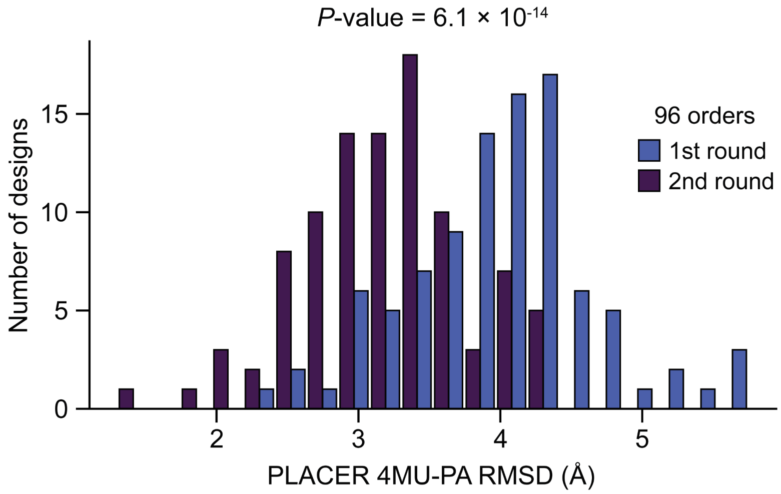

**Fig. S20.** PLACER metrics of ordered designs from both design campaigns. Histogram comparing the distribution of average substrate RMSD between the design model and the predicted PLACER ensembles (PLACER 4MU-PA RMSD) for the 96 designs ordered from the 1st design campaign (blue) and the 96 designs ordered from the 2nd design campaign (purple). The significance was measured using the Wilcoxon rank-sum test, a nonparametric statistical test. The 2nd design campaign featured designs whose substrate positions were predicted by PLACER to be in higher agreement with their design model (lower RMSD), on average.

Fig. S21. Structural panel of the 96 ordered designs spanning 37 unique backbones from the 2nd design campaign.

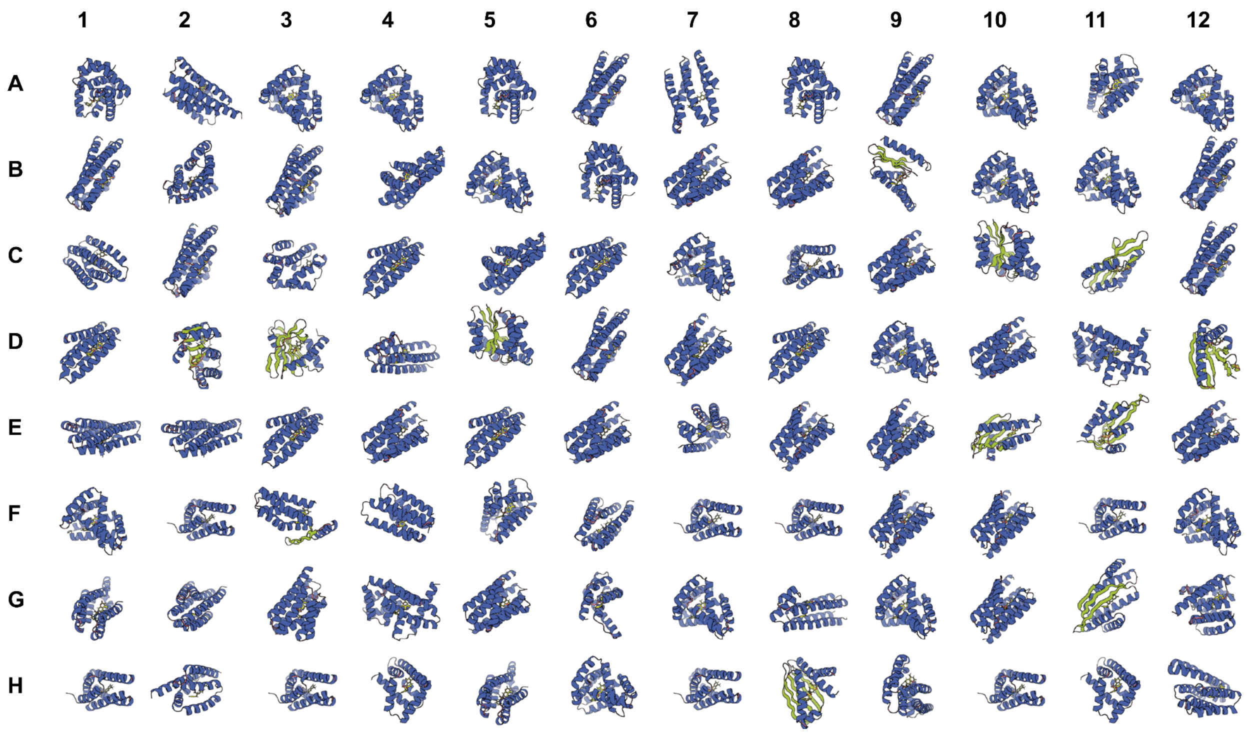

**Fig. S21.** Structural panel of the 96 ordered designs spanning 37 unique backbones from the 2nd design campaign. The secondary structures are colored (green = beta-sheets; blue = alpha-helices). Structures range in length and have a range of diverse folds. Scaffolds were generated in the length range of 110 - 190 amino acids.

Fig. S22. SDS-PAGE gel results for the 96 designs of the 2nd round campaign expressed and purified in parallel.

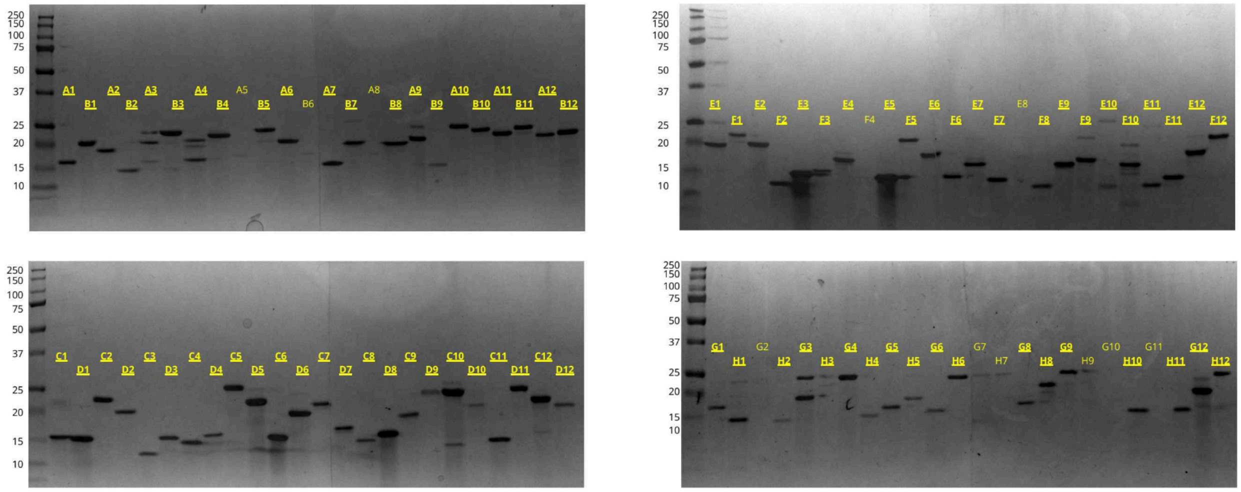

**Fig. S22.** SDS-PAGE gel results for the 96 designs of the 2nd round campaign expressed in *E. coli* and purified in parallel using Strep-Tag affinity chromatography. Designs that had any visible sign of a band were counted as an expressed design (bolded and underlined text). Note that the ladder units are in kDa and that the bottom two gels had an empty spacer well. 87/96 designs show at least one band.

Fig. S23. Michaelis-Menten kinetics of 11 major hits from 2nd design campaign.

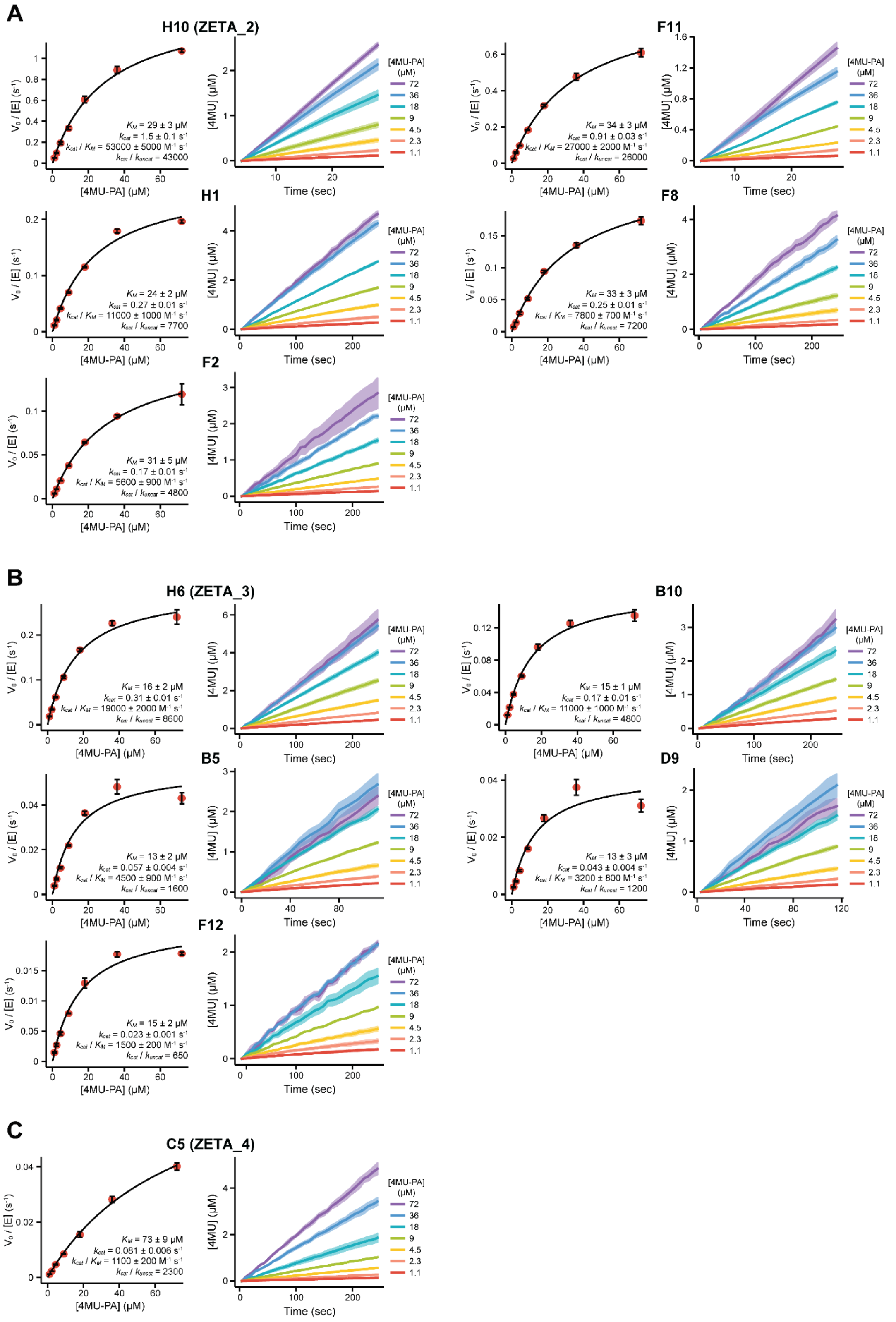

**Fig. S23.** Michaelis-Menten kinetics of 11 major hits from 2nd design campaign. (**A**-**C**) Kinetics of designs grouped by (**A**) ZETA_2 scaffold, (**B**) ZETA_3 scaffold, and (**C**) ZETA_4 scaffold.

Fig. S24. Opposite substrate binding modes in ZETA_2 and ZETA_3 design models resulting from ORI placement.

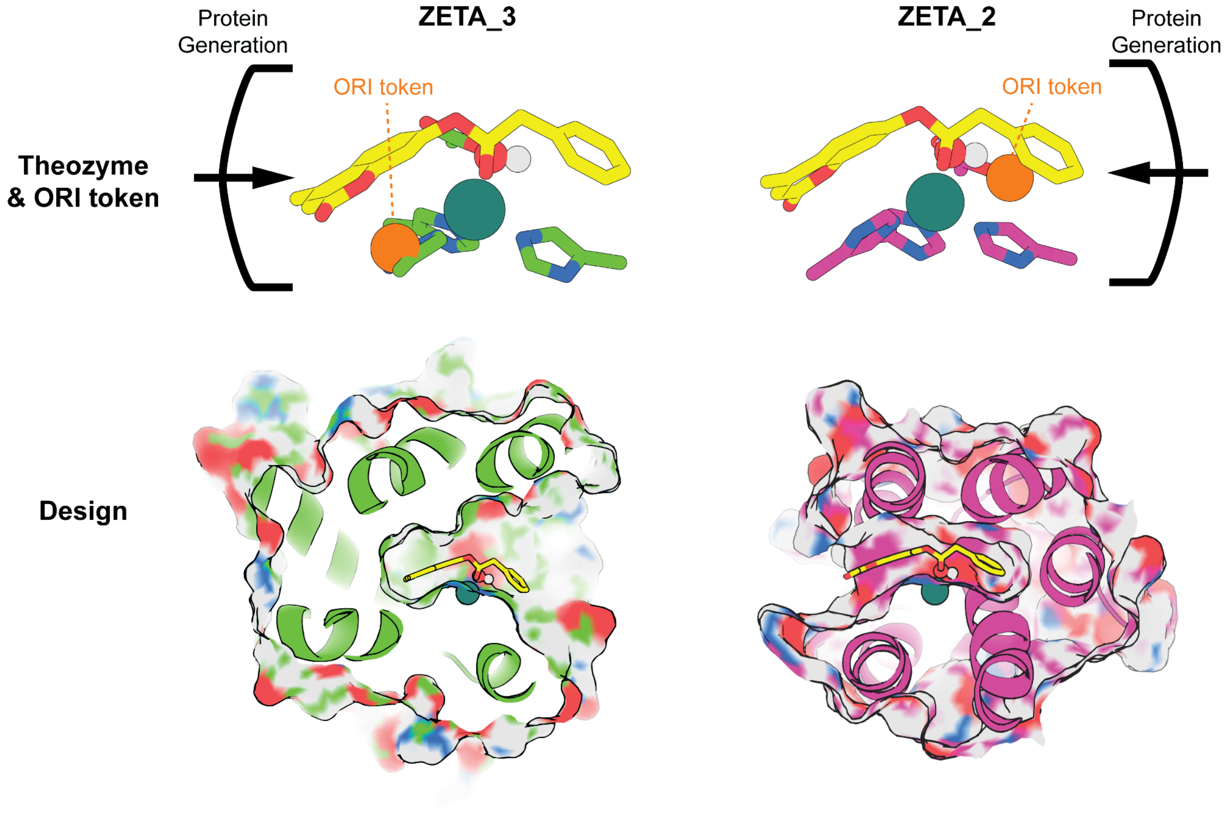

**Fig. S24.** Opposite substrate binding modes in ZETA_2 and ZETA_3 design models result from ORI placement. The ORI token is an optional coordinate fed to the updated RFdiffusion2 which instructs the model on where it should aim to generate the protein center of mass (see SI, section 3.2). Thus, the ORI token can be used to specify which parts of the substrate/theozyme should be more surrounded by protein (buried) on average. ZETA_2 (right) and ZETA_3 (left) are designed to bind the substrate in opposite orientations; this arose from the opposite placement of ORI tokens with respect to the substrate. The bottom panels show a cross-section of the design models for these enzymes with the surface view to observe the designed substrate binding pocket. In ZETA_3 (left) the PA is solvent exposed and the 4MU is buried because the ORI token is next to the 4MU. In ZETA_2 (right) the 4MU is solvent exposed and the PA is buried because the ORI token is next to the PA.

#### 6. SUPPLEMENTAL TABLES

**Table S1.** Michaelis-Menten parameters for the hydrolysis of 4MU-PA by the designed de novo metallohydrolases and mutants.

| Variant | $k_{cat}/K_M$ ($M^{-1}$ $s^{-1}$) | $k_{cat}$ ($s^{-1}$) | $K_M$ (μM) |
|---|---|---|---|
| **1st Design Campaign** | | | |
| A1 (ZETA_1) | 16000 ± 2000 | 0.51 ± 0.03 | 32 ± 4 |
| A8 | 140 ± 10 | 0.012 ± 0.001 | 85 ± 5 |
| B9 | 69 ± 14 | 0.014 ± 0.001 | 200 ± 30 |
| C4 | 100 ± 10 | 0.0028 ± 0.0001 | 27 ± 2 |
| F7 | 35 ± 5 | 0.0035 ± 0.0003 | 100 ± 10 |
| **A1 (ZETA_1) Knockout Mutants** | | | |
| A1 H130A | 1300 ± 100 | 0.020 ± 0.001 | 16 ± 2 |
| A1 N17A | 2000 ± 100 | 0.031 ± 0.001 | 16 ± 1 |
| A1 D67A | 14000 ± 2000 | 0.17 ± 0.01 | 12 ± 1 |
| A1 H118A | Inactive | Inactive | Inactive |
| A1 H134A | Inactive | Inactive | Inactive |
| A1 H118A;H130A;H134A | Inactive | Inactive | Inactive |
| **2nd Design Campaign** | | | |
| H10 (ZETA_2) | 53000 ± 5000 | 1.5 ± 0.1 | 29 ± 3 |
| F11 | 27000 ± 2000 | 0.91 ± 0.03 | 34 ± 3 |
| H1 | 11000 ± 1000 | 0.27 ± 0.01 | 24 ± 2 |
| F8 | 7800 ± 700 | 0.25 ± 0.01 | 33 ± 3 |
| F2 | 5600 ± 900 | 0.17 ± 0.01 | 31 ± 5 |
| H6 (ZETA_3) | 19000 ± 2000 | 0.31 ± 0.01 | 16 ± 2 |
| B10 | 11000 ± 1000 | 0.17 ± 0.01 | 15 ± 1 |
| B5 | 4500 ± 900 | 0.057 ± 0.004 | 13 ± 2 |
| D9 | 3200 ± 800 | 0.043 ± 0.004 | 13 ± 3 |
| F12 | 1500 ± 200 | 0.023 ± 0.001 | 15 ± 2 |
| C5 (ZETA_4) | 1100 ± 200 | 0.081 ± 0.006 | 73 ± 9 |

**Table S2.** Electrospray ionization mass spectrometry results for the characterized hits and A1 mutants.

| Design | (E) Expected Mass (Da) | (O) Observed Mass (Da) | Δ (O - E) |
|---|---|---|---|
| **1st Design Campaign** | | | |
| A1 (ZETA_1) | 18528 | 18397 | -131 |
| A8 | 25301 | 25170 | -131 |
| B9 | 20202 | 20071 | -131 |
| C4 | 18724 | 18593 | -131 |
| F7 | 21023 | 20892 | -131 |
| **A1 (ZETA_1) Knockout Mutants** | | | |
| A1 H118A;H130A;H134A | 18330 | 18199 | -131 |
| A1 H130A | 18462 | 18331 | -131 |
| A1 H134A | 18462 | 18331 | -131 |
| A1 H118A | 18462 | 18331 | -131 |
| A1 N17A | 18485 | 18354 | -131 |
| A1 D67A | 18484 | 18353 | -131 |
| **2nd Design Campaign** | | | |
| H10 (ZETA_2) | 15748 | 15616 | -132 |
| H6 (ZETA_3) | 22046 | 21915 | -131 |
| C5 (ZETA_4) | 22585 | 22454 | -131 |

Note 1: In all cases the observed mass corresponded to the loss of N-terminal methionine (-131 Da).
Note 2: The mass spectrometer has an error of approximately ±1 Da.

**Table S3.** Comparing the catalytic efficiencies of different designed and engineered (metallo)enzymes.

| | **Substrate*** | **pH** | $k_{cat}$ (s$^{-1}$) | $k_{cat}/K_M$ (M$^{-1}$ s$^{-1}$) | $k_{cat}/k_{uncat}$ | **Ref.** |
|---|---|---|---|---|---|---|
| ZETA_1 | 4MU-PA | 8.0 | 0.51 | $1.6 \times 10^{4}$ | $1.4 \times 10^{4}$ | |
| ZETA_2 | 4MU-PA | 8.0 | 1.5 | $5.3 \times 10^{4}$ | $4.3 \times 10^{4}$ | |
| ZETA_3 | 4MU-PA | 8.0 | 0.31 | $1.9 \times 10^{4}$ | $8.6 \times 10^{3}$ | |
| ZETA_4 | 4MU-PA | 8.0 | 0.081 | $1.1 \times 10^{3}$ | $2.3 \times 10^{3}$ | |
| **Computationally Designed Zinc Esterases (from scratch with novel scaffolds; no laboratory evolution)** | | | | | | |
| MID1 §,‖ | *p*NPA | 8.0 | 0.13 | 40 | $9.4 \times 10^{3}$ | 29 |
| **Computationally Designed/Engineered Zinc Esterases (from existing scaffolds)** | | | | | | |
| (TRIL9CL23H)$_3$ ‖ | *p*NPA | 8.0 | 0.0054 | 3.1 | $3.4 \times 10^{2}$ | 30 |
| $^{A104}$AB3 §,‖ | *p*NPA | 9.0 | 0.0068 | 8 | $6.2 \times 10^{1}$ | 31 |
| Ac-IHIHIQI-CONH$_2$ ‖ | *p*NPA | 8.0 | 0.026 | 62 | $1.6 \times 10^{3}$ | 32 |
| **Computationally Designed Esterases (from scratch with novel scaffolds; no laboratory evolution)** | | | | | | |
| win11 | 4MU-Ac | 7.4 | 0.0197 | $3.9 \times 10^{3}$ | $5.1 \times 10^{3}$ | 33 |
| **Computationally Designed/Engineered Esterases (from existing scaffolds)** | | | | | | |
| FR29 ‖,** | *p*NPP | 7.5 | 0.0041 | 34 | $5.5 \times 10^{2}$ | 34 |
| ECH13 ‖,** | *p*NPP | 7.5 | 0.018 | 309 | $2.4 \times 10^{3}$ | 34 |
| ECH19 ‖,** | *p*NPP | 7.5 | 0.01 | 227 | $1.4 \times 10^{3}$ | 34 |
| AlleyCatE ‖ | (*R*)-*p*NPP | 7.5 | 0.004 | 5.5 | $5.4 \times 10^{2}$ | 35 |
| CC-Hept-Cys-His-Glu ‖,** | *p*NPA | 7.0 | 0.0005 | 3.7 | $1.4 \times 10^{2}$ | 36 |
| PZD2 ‖ | *p*NPA | 7.0 | 0.00046 | 2.7 | $1.3 \times 10^{2}$ | 37 |
| **Engineered Esterases (from combinatorial libraries of peptides)** | | | | | | |
| S-824 ‖ | *p*NPA | 7.3 | 0.0055 | 1.8 | $7.4 \times 10^{2}$ | 38 |
| **Computationally Designed/Engineered Non-Hydrolytic Enzymes (from scratch)** | | | | | | |
| G$_4$-DF$_{tet}$ † | 4AP | 7.0 | 0.021 | 25 | — | 39 |
| **Computationally Designed Non-Hydrolytic Enzymes (from existing scaffolds; no laboratory evolution)** | | | | | | |
| DA_20_00 † | CBBC/DMAA | 7.4 | 0.000028 | 0.06 ‡ | 4.1 ‡ | 40 |
| BH32 † | F2PP | — | ~0.017 | ~330 | — | 41 |
| LuxSit † | DTZ | 8.0 | ~0.025 | ~$1.4 \times 10^{3}$ | — | 42 |
| RAD29 † | Methodol | 7.5 | 0.0314 | 290 | $5 \times 10^{6}$ | 43 |
| RA95 † | Methodol | 7.5 | 0.0020 | 0.053 | $4.8 \times 10^{3}$ | 44 |
| cRA-50 † | Methodol | — | 2.2 | 180 | — | 45 |
| dnHEM1 † | Amplex Red | 7.2 | 9.5 | 260 | — | 46 |
| HG3 † | 5-NBZ | 7.25 | 0.68 | 425 | $5.9 \times 10^{5}$ | 47 |
| KE70 † | 5-NBZ | 7.25 | 0.16 | 80 | $1.4 \times 10^{5}$ | 48 |
| **Computational Sequence-Redesigned/-Optimized Enzymes (from existing active enzymes)** | | | | | | |
| HG259 † | 5-NBZ | 7.0 | — | $13 \times 10^{4}$ | — | 49 |
| HG630 † | 5-NBZ | 7.0 | — | $15 \times 10^{4}$ | — | 49 |
| HG649 † | 5-NBZ | 7.0 | — | $32 \times 10^{4}$ | — | 49 |
| KE703b † | 5-NBZ | 7.0 | — | $5.5 \times 10^{3}$ | — | 49 |
| KGSADH$_{V18C}$ † | 3-HPA | 7.4 | 7.0 | $4.0 \times 10^{3}$ | — | 50 |
| **Catalytic Antibodies** | | | | | | |
| 43C9 ‖ | *p*NP-ester | 9.3 | 25 | $4.7 \times 10^{5}$ | $2.7 \times 10^{4}$ | 51 |
| D2.4 ‖ | *p*NP-ester | 8.3 | 0.22 | $9.4 \times 10^{3}$ | $5.4 \times 10^{3}$ | 52 |
| CNJ157 ‖ | *p*NP-ester | 8.0 | 0.04 | 360 | $2.5 \times 10^{3}$ | 53 |
| 48G7 ‖ | *p*NP-ester | 8.2 | 0.092 | 230 | $5.8 \times 10^{3}$ | 54 |
| CNJ206 ‖ | *p*NPA | 8.0 | 0.0067 | 83 | $4.2 \times 10^{2}$ | 55 |
| **Natural Enzymes** | | | | | | |
| *S. solfataricus* lipase †† | *p*NPB | 6.0 | 1100 | $8.8 \times 10^{6}$ | — | 56 |
| *C. albicans* lipase ‖ | *p*NPB | 6.0 | 54 | $1.3 \times 10^{5}$ | $1.7 \times 10^{7}$ | 57 |
| hCA II ‖ | *p*NPA | 8.0 | 53 | $2.5 \times 10^{3}$ | $3.3 \times 10^{6}$ | 58 |
| *Staph. xylosus* lipase †† | *p*NPB | 9.0 | 0.061 | 140 | — | 59 |

Note 1: Table adapted from previous tables.[29,36]

Note 2: Catalytic antibodies and native metallohydrolases are included for reference.

Note 3: According to the sorting in this table, ZETA_1-4 would be classified under `Computationally Designed Zinc Esterases (from scratch with novel scaffolds; no laboratory evolution)`

* Substrate abbreviation. 4MU-PA: 4-Methylumbelliferyl phenylacetate; 4MU-Ac: 4-Methylumbelliferyl acetate; *p*NPA: *p*-nitrophenylacetate; *p*NPB: *p*-nitrophenylbutyrate; *p*NPP: *p*-nitrophenyl-2-propanoate; *p*NP-ester: 5-((4-(2-(4-nitrophenoxy)-2-oxoethyl)phenyl)amino)-5-oxopentanoic acid (43C9), p-nitrophenyl-N-glycylglutarate (D2.4), 4-(5-(4-nitrophenoxy)-5-oxopentanamido)butanoic acid (CNJ157), 6-(4-nitrophenoxy)-6-oxohexanoic acid (48G7). 5-NBZ: 5-nitro-1,2-benzisoxazole; DTZ: diphenylterazine; CBBC/DMAA: 4-carboxybenzyl trans-1,3-butadiene-1-carbamate and N,N-dimethylacrylamide; F2PP: fluorescein 2-phenylpropanoate; 4AP: 4-aminophenol; 3-HPA: 3-hydroxypropionaldehyde

** Only the catalytic parameters for enzyme acylation were reported; the deacylation step is considerably slower and limits overall turnover efficiency.

† Non-esterase enzymes. $G_4$-$DF_{tet}$: diiron $O_2$-dependent phenol oxidase; DA_20_00: Diels-Alderase; BH32: Morita-Baylis-Hillman reaction enzyme catalyst; LuxSit: luciferase; RA95, cRA-50, RAD29: retroaldolase; dnHEM1: heme peroxidase; HG series, KE series: kemp eliminase; $KGSADH_{Y68T}$: alpha-ketoglutaric semialdehyde dehydrogenase.

|| Rate enhancement ($k_{cat}/k_{uncat}$) was calculated using $k_{buffer}$ for *p*NPA hydrolysis from a previous measurement[60]: pH 9.0, 1.1 × $10^{-4}$ $s^{-1}$; pH 8.5, 4.1 × $10^{-5}$ $s^{-1}$; pH 8.0, 1.6 × $10^{-5}$ $s^{-1}$; pH 7.5, 7.4 × $10^{-6}$ $s^{-1}$; pH 7.0, 3.5 × 10-6 $s^{-1}$; pH 6, 2.9 × $10^{-6}$ $s^{-1}$ (for pH 6.0, relative rate constant was abstracted from another previous measurement[61]).

‡ $k_{cat}/K_{M\text{-}diene}\ K_{M\text{-}dienophile}$ for the diels-alderase; $k_{cat}/k_{uncat}$ has units of M in this instance.

~ Estimated values for (1) LuxSit since only the $K_M$ = 18.3 μM and $V_{max}$ = 0.0032 photon $s^{-1}$ molecule$^{-1}$ were reported, where enzyme concentration nor luminescent quantum yield were reported to determine $k_{cat}$. The successful mutagenic variant LuxSit-i had $V_{max}$ = 0.36 photon $s^{-1}$ molecule$^{-1}$ and $k_{cat}$ = 2.5 $s^{-1}$ reported; thus, we assumed the $k_{cat}$ for LuxSit was also two orders of magnitude lower since its $V_{max}$ was two orders of magnitude lower and $V_{max}$ is proportional to $k_{cat}$. (2) Estimated values for BH32 using the published extended data figure 1 showing a Michaelis-Menten plot; $k_{cat}/K_M$ was estimated as the initial slope.

§ Normalized to a single zinc site.

†† Kinetic parameters determined at elevated temperatures: 70 °C for *S. solfataricus* lipase; 42 °C for *Staph. xylosus* lipase

#### 7. SUPPLEMENTAL MOVIES

Note: Movies will be available upon publication.

**Movie S1.** RFdiffusion2 trajectory for ZETA_1 (A1).

**Movie S2.** ChemNet predictions for substrate ensemble of ZETA_1 (A1).

**Movie S3.** ChemNet predictions for catalytic residue ensemble of ZETA_1 (A1).

**Movie S4.** ChemNet predictions for substrate ensemble of the inactive design H8.

**Movie S5.** ChemNet predictions for catalytic residue ensemble of the inactive design H7.
